# Identification of key genes and signaling pathway in the pathogenesis of Huntington’s disease via bioinformatics and next generation sequencing data analysis

**DOI:** 10.1101/2024.06.20.599879

**Authors:** Basavaraj Vastrad, Chanabasayya Vastrad

## Abstract

Huntington’s disease (HD) is the primary cause of progressive motor deficits, psychiatric symptoms, and cognitive impairment. The exact molecular mechanisms of HD pathogenesis are largely unknown. This investigation aims to identify the hub genes, miRNA and TFs in HD and explore their potential molecular regulatory network. Next generation sequencing (NGS) dataset (GSE105041) was extracted from the Gene Expression Omnibus (GEO) database. An integrated bioinformatics pipeline including identification of differentially expressed genes (DEGs), Gene ontology and REACTOME pathway enrichment analysis, protein-protein interaction (PPI) network and module analysis, miRNA-hub gene regulatory network analysis and TF-hub gene regulatory network analysis, and receiver operating characteristic curve (ROC) analysis were applied to identify hub genes, miRNA and TFs, and key drivers of HD pathogenesis. We identified 958 DEGs in the discovery phase, consisting of 479 up regulated genes and 479 down regulated genes. GO and REACTOME enrichment analyses of the 479 up regulated genes and 479 down regulated genes showed that they were mainly involved in multicellular organismal process, developmental process, signaling by GPCR and MHC class II antigen presentation. Further analysis of the PPI network using Cytoscape and PEWCC plugins identified 10 hub genes, including LRRK2, MTUS2, HOXA1, IL7R, ERBB3, EGFR, TEX101, WDR76, NEDD4L and COMT. Possible target miRNAs and TFs, including hsa-mir-1292-5p, hsa-mir-4521, ESRRB and SREBF1, were predicted by constructing a miRNA-hub gene regulatory network analysis and TF-hub gene regulatory network. This investigation used bioinformatics methods to explore the molecular pathogenesis of HD, and identified potential molecular markers. These molecular markers might provide novel ideas and methods for the early diagnosis, treatment, and monitoring of HD.

## Introduction

Huntington’s disease (HD) is a rare, inherited, neurodegenerative disorder that manifests as progressive motor deficits, psychiatric symptoms, and cognitive impairment [1]. It is estimated to affect 2.7 per 100,000 individuals of the global population [2]. HD prevalence observed in childhood, adolescence, and adulthood, but in most cases, this occurs in middle aged adulthood [3]. HD is characterized by involuntary movements, slight difficulty with executive functions, and depressed mood [4]. Several risk factors have been proposed to cause HD, including genetic inheritance [5], environmental variables [6], inflammation [7], hypertension [8], cardiovascular diseases [9] and diabetes mellitus [10]. Despite emerging proof providing considerable mechanistic insight into this condition, the exact molecular mechanism of HD is still being debated. As HD has not clear picture of pathogenesis and a not satisfactory response to drug treatment, it is necessary to explore the molecular mechanism of HD to develop effective diagnosis, prognosis and target treatments.

Several investigations has described that significant molecular biomarkers and signaling pathways in HD were identified as prognostic or therapeutic factors, such as Huntingtin (HTT) protein [11], arachidonate 5-lipoxygenase (ALOX5) [12], type II CRISPR RNA-guided endonuclease Cas9 (CRISPR/Cas9) [13], interleukin-6 (IL6) [14], RE1 silencing transcription Factor (REST) [15], insulin/IGF1 (IIS) signaling pathway [16], hippo signaling pathway [17], PI3K/AKT/FoxO1 signaling pathway [18], cAMP/CREB/BDNF signaling pathway [19] and GLP-1 Receptor/PI3K/Akt/BDNF signaling pathway [20]. The identification of novel biomarkers and signaling pathway might be helpful to improve the clinical outcome of HD patients.

Bioinformatics and next generation sequencing (NGS) data analysis are a well-orchestrated tool for screening prognosis and diagnosis relevant biomarkers, which can contribute to the advancement of HD treatment [21]. At current, NGS data downloaded from Gene Expression Omnibus (GEO) database [https://www.ncbi.nlm.nih.gov/geo/] [22] can be used to detect genes expression levels and to provide the technical support for monitoring mRNA expression and cell function prediction in various diseases [23–24].

During the last decades, NGS technology and bioinformatics analysis have been widely used to screen genetic alterations at the genome level, which have helped us identify the differentially expressed genes (DEGs) and functional pathways involved in the progression of HD. Thus, in the current investigation, NGS dataset (GSE105041) [25] from GEO was downloaded and analyzed to obtain DEGs between HD and normal control samples. Subsequently, Gene Ontology (GO), REACTOME pathway enrichment analysis protein-protein interaction (PPI) network analyses, miRNA-hub gene regulatory network analyses and TF-hub gene regulatory network analyses were performed to help us understand the molecular mechanisms underlying progression of HD. Finally, candidate hub genes were verified by receiver operating characteristic curve (ROC) analysis. In short, these results indicated that the biomarkers might have potential value in prognosis, diagnosis and therapeutic targets of HD.

## Materials and Methods

### Next generation sequencing (NGS) data source

GSE105041 [25] NGS dataset was downloaded from the GEO database. The GSE105041 dataset was generated utilizing the GPL16791 Illumina HiSeq 2500 (Homo sapiens) platform and contained 28 HD samples and 20 normal control samples.

### Identification of DEGs

The DEGs between HD and normal control samples were screened using DESeq2 [26] . DESeq2 is an R Bioconductor tool that allows users to identify DEGs across experimental conditions. The adjusted P-values (adj. P) and Benjamini and Hochberg false discovery rate were applied to provide a balance between discovery of statistically significant genes and limitations of false-positives [27]. logFC (fold change) > 1.966 for up regulated genes, logFC (fold change) < -0.7343 for down regulated genes and adj. P-value <0.05 were considered statistically significant. Significant DEGs were visualized using volcano plot and heatmaps. Both visualizations were completed by“ggplot2” and “gplot” an R Bioconductor packages for data analysis and visualization.

### GO and pathway enrichment analyses of DEGs

Gene Ontology (GO) annotation (http://www.geneontology.org) [28] and REACTOME (https://reactome.org/) [29] pathway enrichment analysis were performed for DEGs. GO Annotation such as analysis of biological processes (BP), cell components (CC), and molecular functions (MF). REACTOME is an online database for pathway enrichment analysis of a large amount of genetic information. GO functional annotation analysis and REACTOME pathway enrichment analysis were performed in the website g:Profiler (http://biit.cs.ut.ee/gprofiler/) [30] import our gene name into the website for enrichment analysis. The significant enrichment threshold was set as p-value < 0.05, GO functions and pathways with the lowest p-values were selected as the top.

### Construction of the PPI network and module analysis

The protein-protein interaction (PPI) network was created from the “Integrated Interactions Database (IID)” (http://iid.ophid.utoronto.ca/search_by_proteins/) [31] database for DEGs and is designed to discover the interaction of DEGs and their interactions with other genes, and Cytoscape (v 3.10.2) (http://www.cytoscape.org/) [32] was used to edit the graphics. The node degree [33], betweenness [34], stress [35] and closeness [36] algorithms of Network Analyzer in Cytoscape was used to explore hub genes. The plug-in PEWCC Cytoscape software is an APP for clustering a given network based on topology to find densely connected regions in PPI network [37]. The most significant module in the PPI networks was identified using PEWCC. Subsequently, the GO and REACTOME pathway enrichment analyses and for genes in this module were performed using g:Profiler.

### Construction of the miRNA-hub gene regulatory network

miRNAs constitute major control modes of gene expression, negatively regulate target genes by combining with the 3′ untranslated region of mRNA. To choose the miRNAs, miRNA-hub gene regulatory network has been visualized by the miRNet (https://www.mirnet.ca/) [38] online platform, which has been widely used as a bioinformatics tool. Constructed a miRNA-hub gene regulatory network graph using Cytoscape (v. 3.10.2) http://www.cytoscape.org/) [32] to show the interactions intuitively.

### Construction of the TF-hub gene regulatory network

TFs constitute major control modes of gene expression, such as transcription and post transcription. To choose the TFs, TF-hub gene regulatory network has been visualized by the NetworkAnalyst database (https://www.networkanalyst.ca/) [39] online platform, which has been widely used as a bioinformatics tool. Constructed a TF-hub gene regulatory network graph using Cytoscape (v. 3.10.2) http://www.cytoscape.org/) [32] to show the interactions intuitively.

### Receiver operating characteristic curve (ROC) analysis

The multivariate modeling with combined selected hub genes were used to identify molecular biomarkers with high sensitivity and specificity for HD diagnosis. One data used as training dataset and other data used as validation dataset iteratively. The receiver operator characteristic curves were plotted and area under curve (AUC) was determined separately to check the performance of each model using the R packages “pROC” [40]. A AUC > 0.8 designated that the model had a excellent fitting effect.

## Results

### Identification of DEGs

GSE105041 was selected and underwent DEGs analysis using “DESeq2” package in R software. 958 DEGs were identified either up or down regulated in all, including 479 up and 479 down regulated genes (|logFC (fold change) > 1.966 for up regulated genes, logFC (fold change) < -0.7343 for down regulated genes and adj. P-value <0.05) (Table 1). As shown in Fig 1, all 958 DEGs were plotted that green ones represented up regulation, red ones indicated down regulation. What’s more, the expression levels of all the DEGs were presented in the heatmap (Fig. 2), and these genes were well clustered between HD and normal control samples.

**Table 1.**
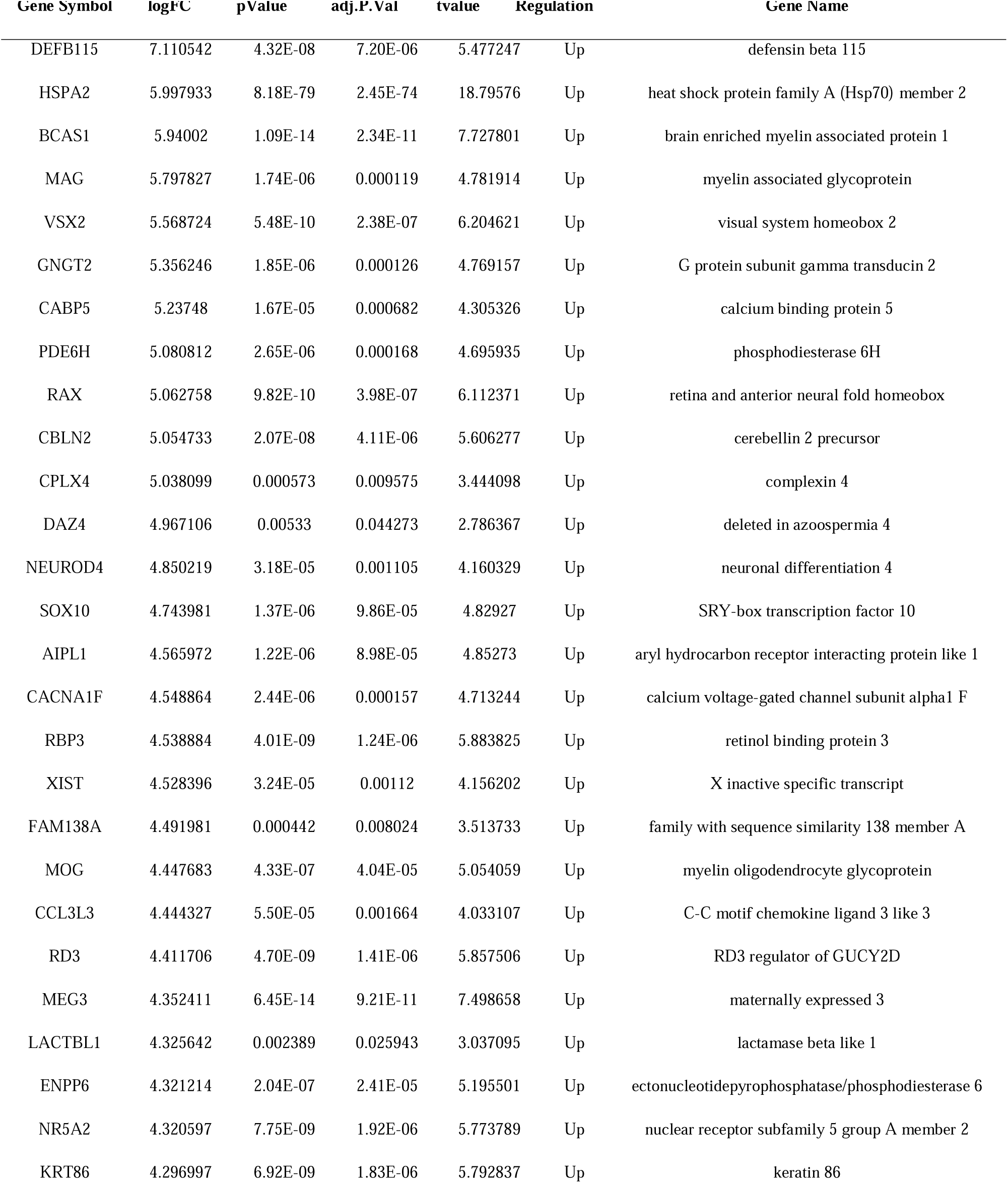
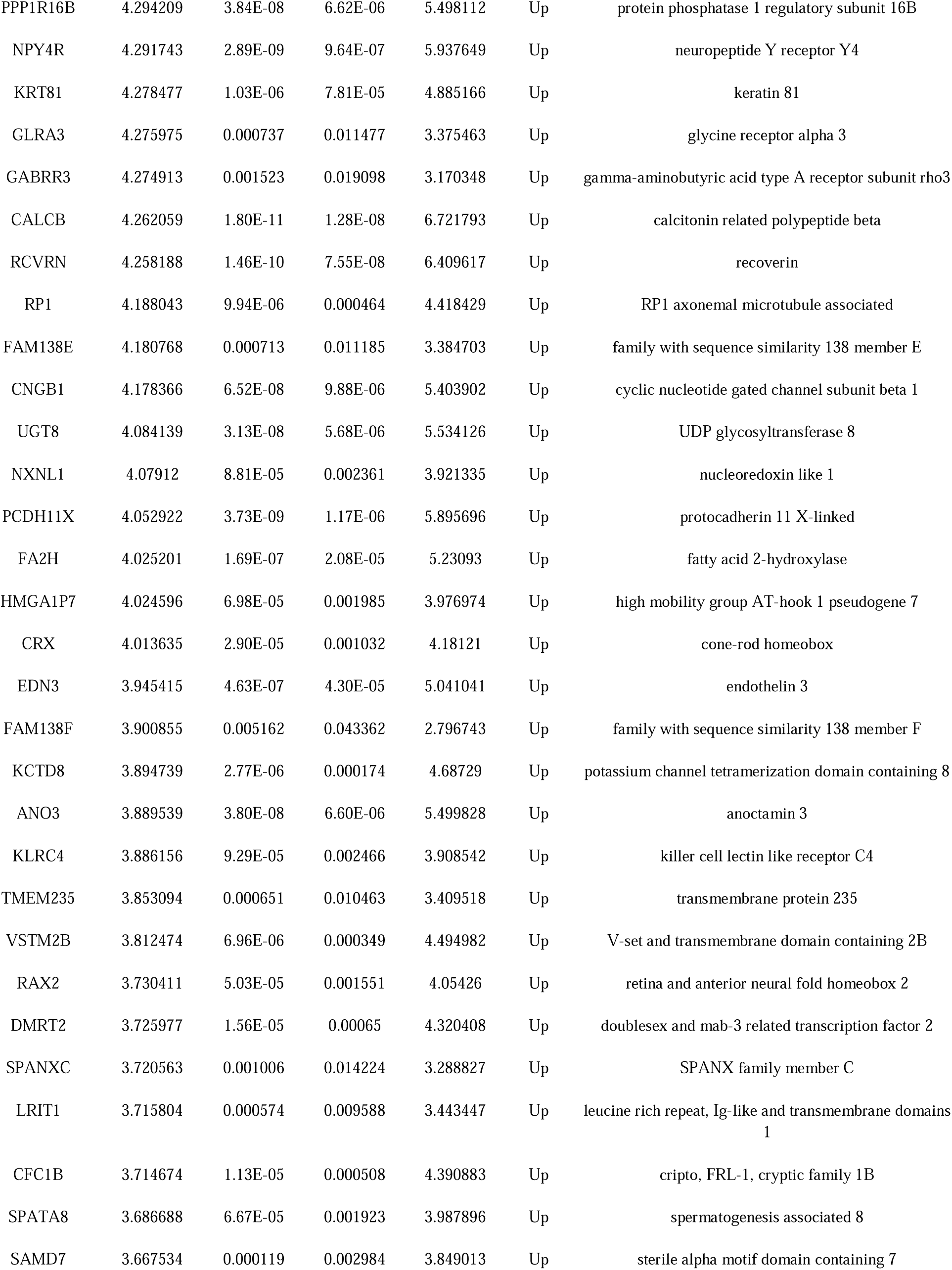
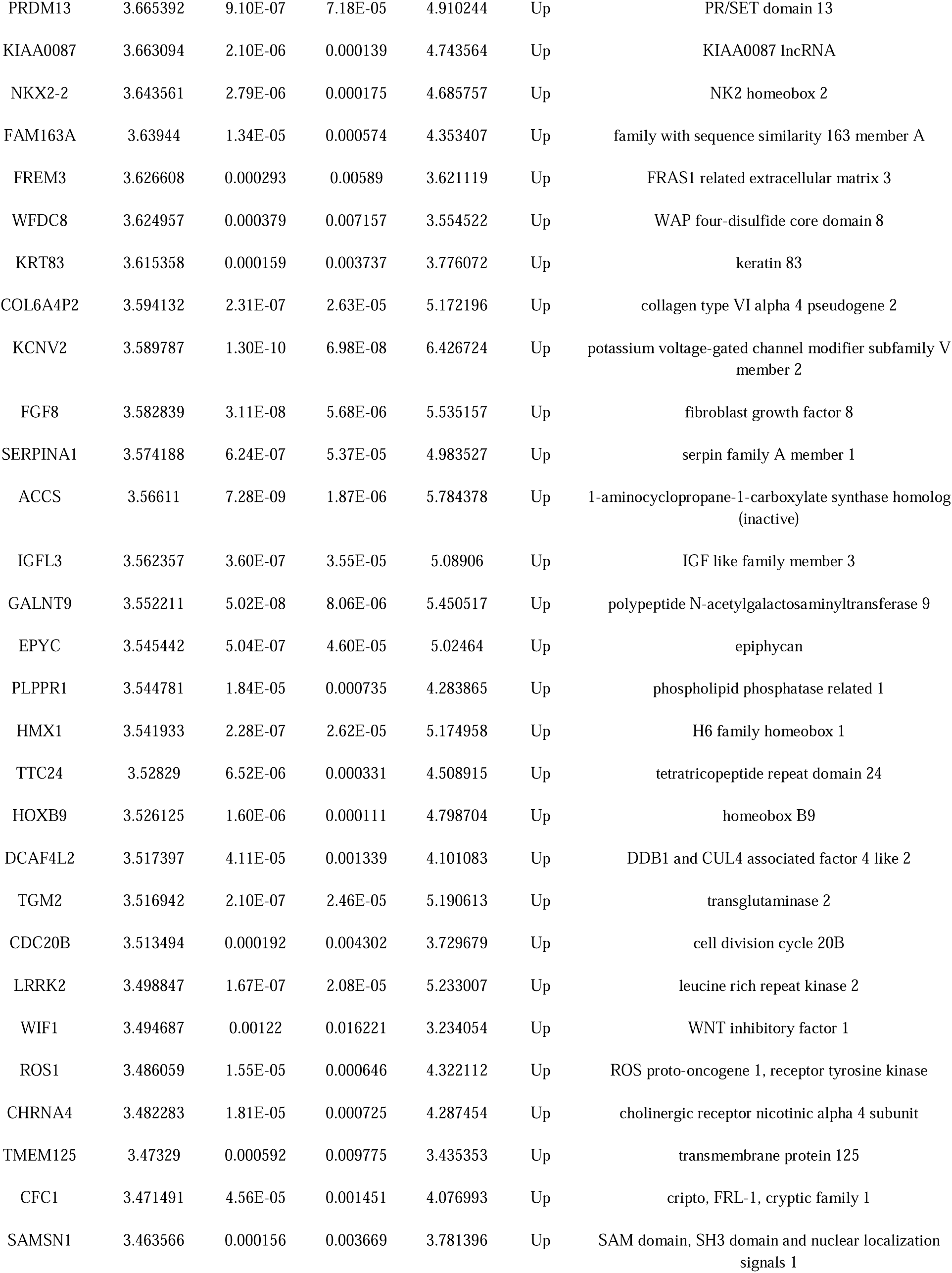
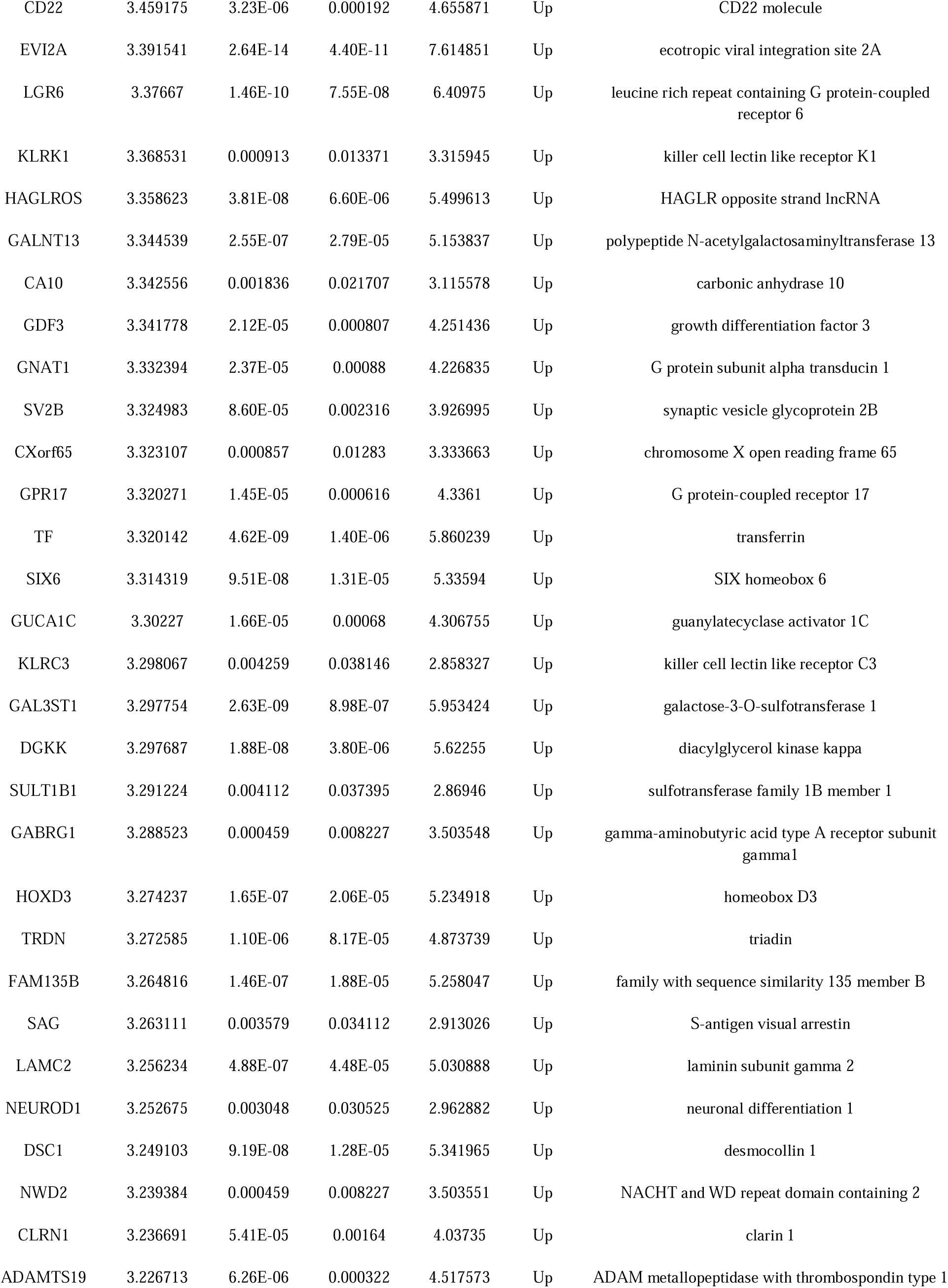
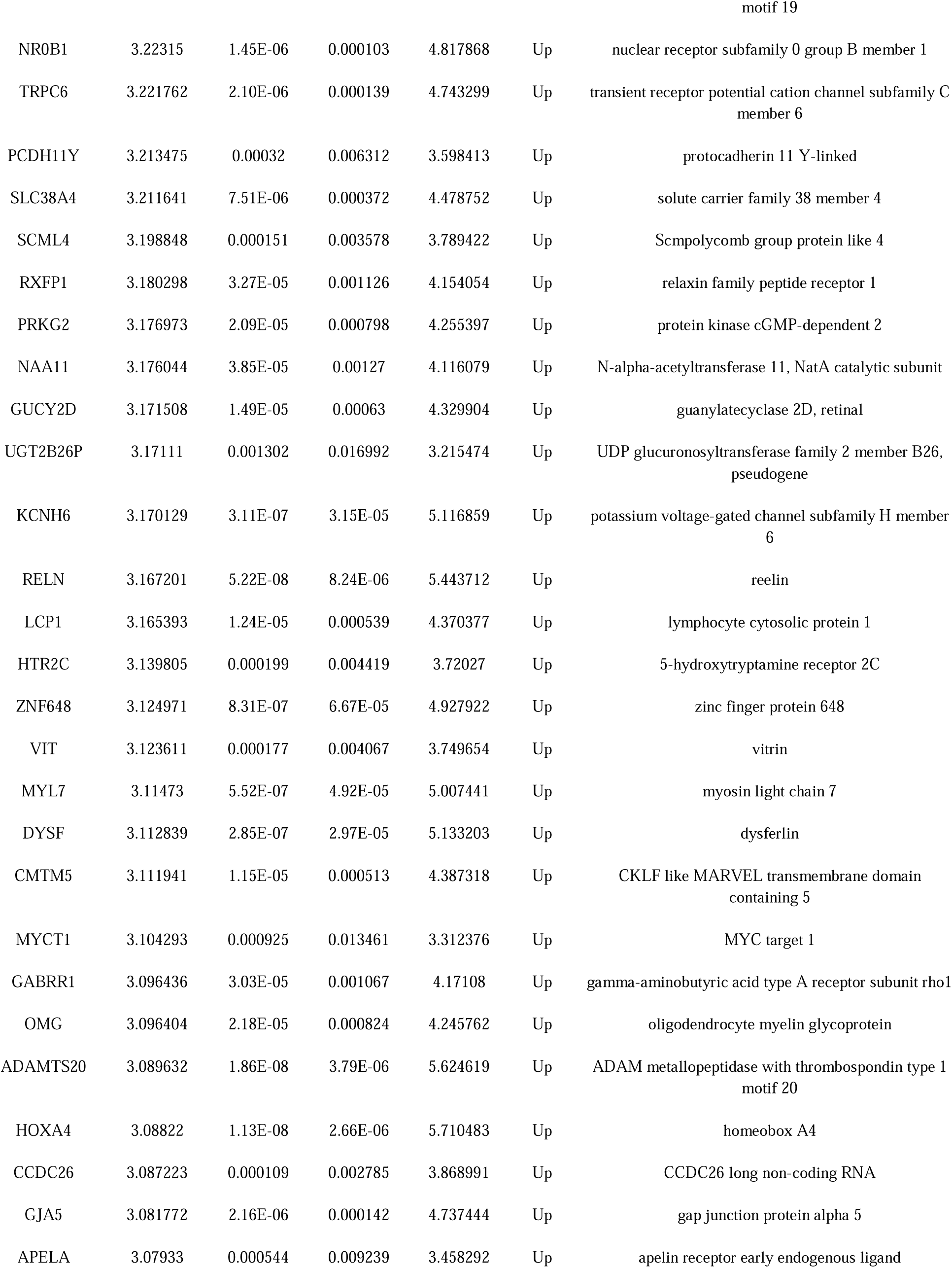
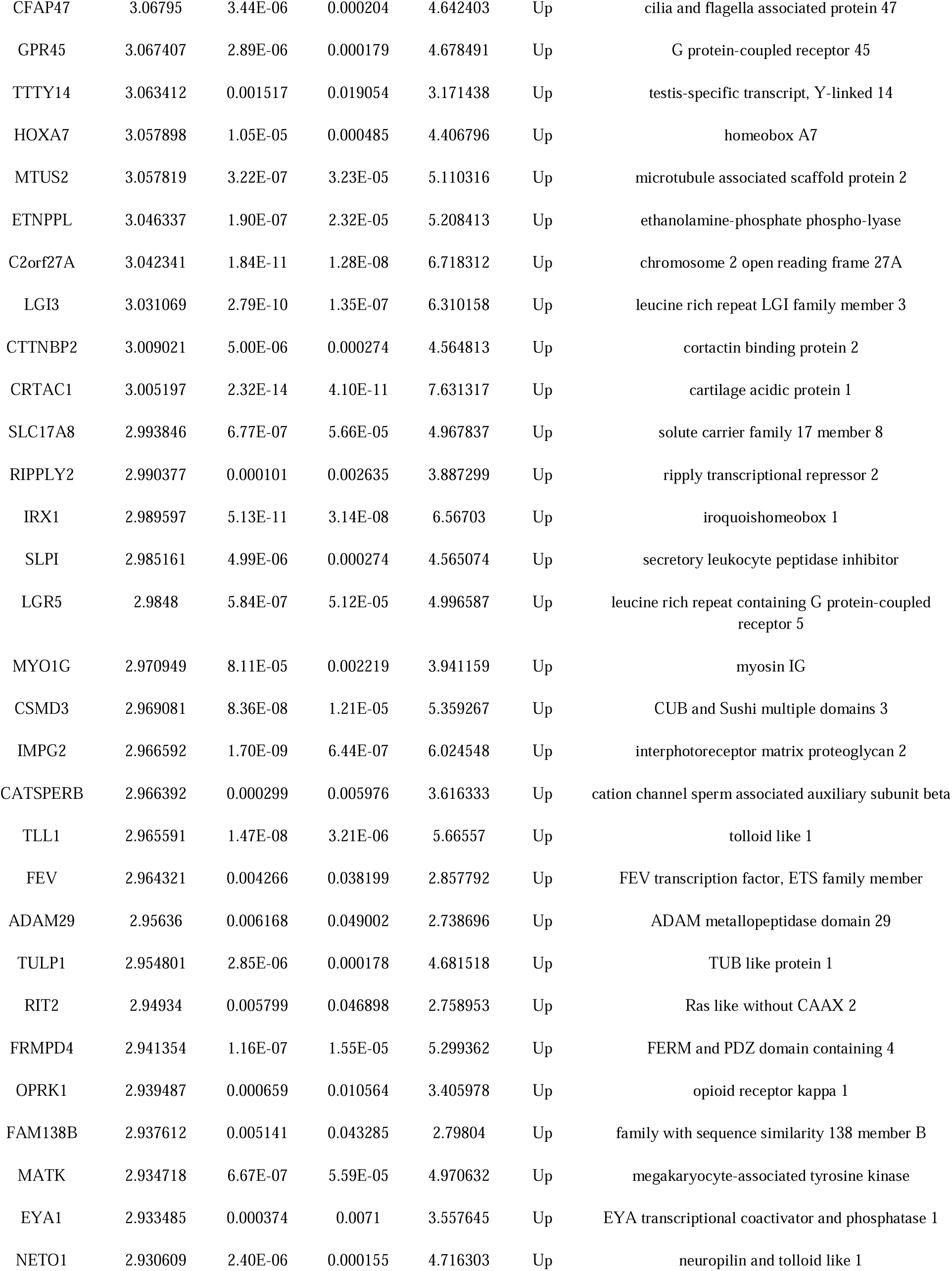
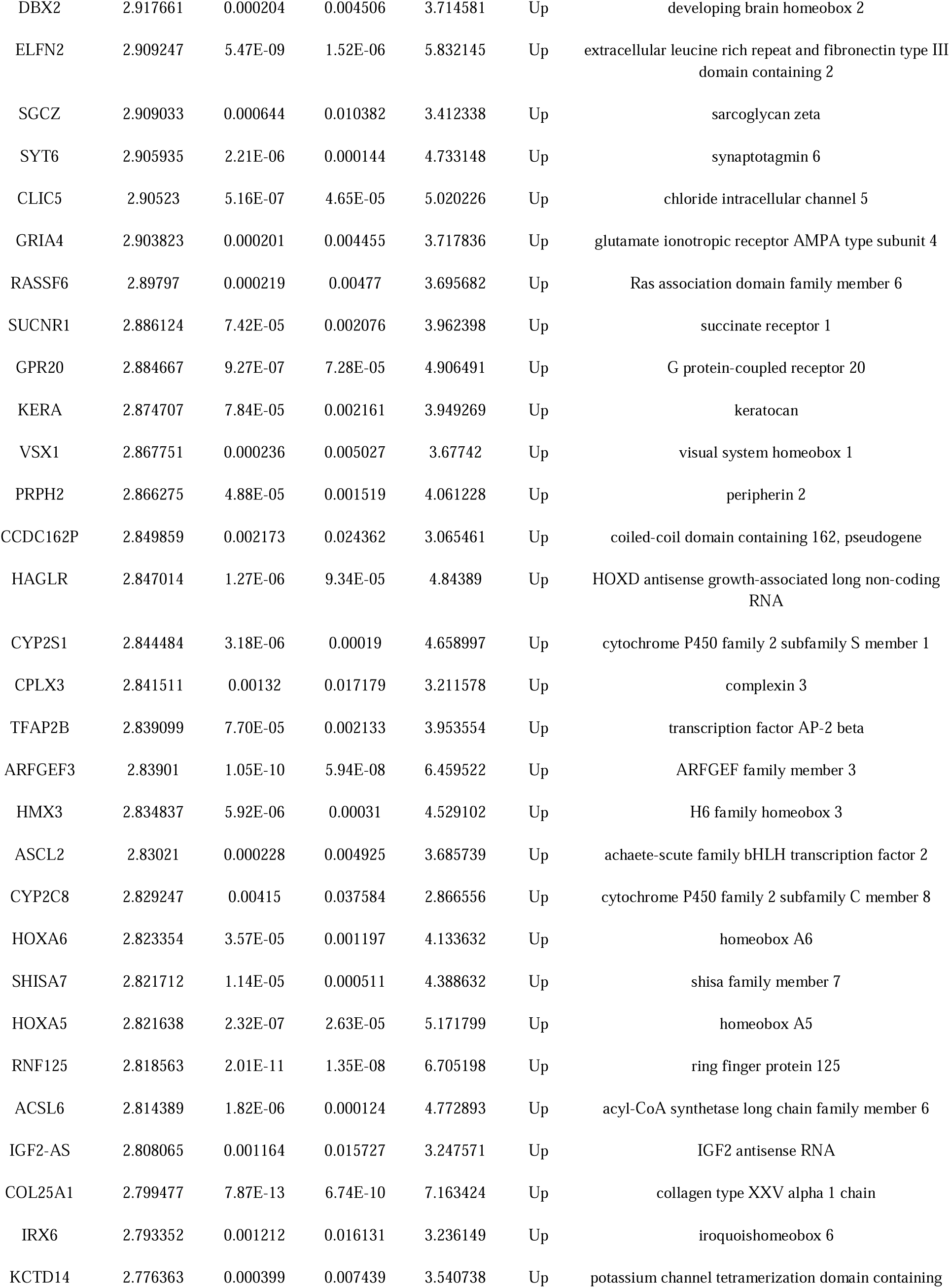
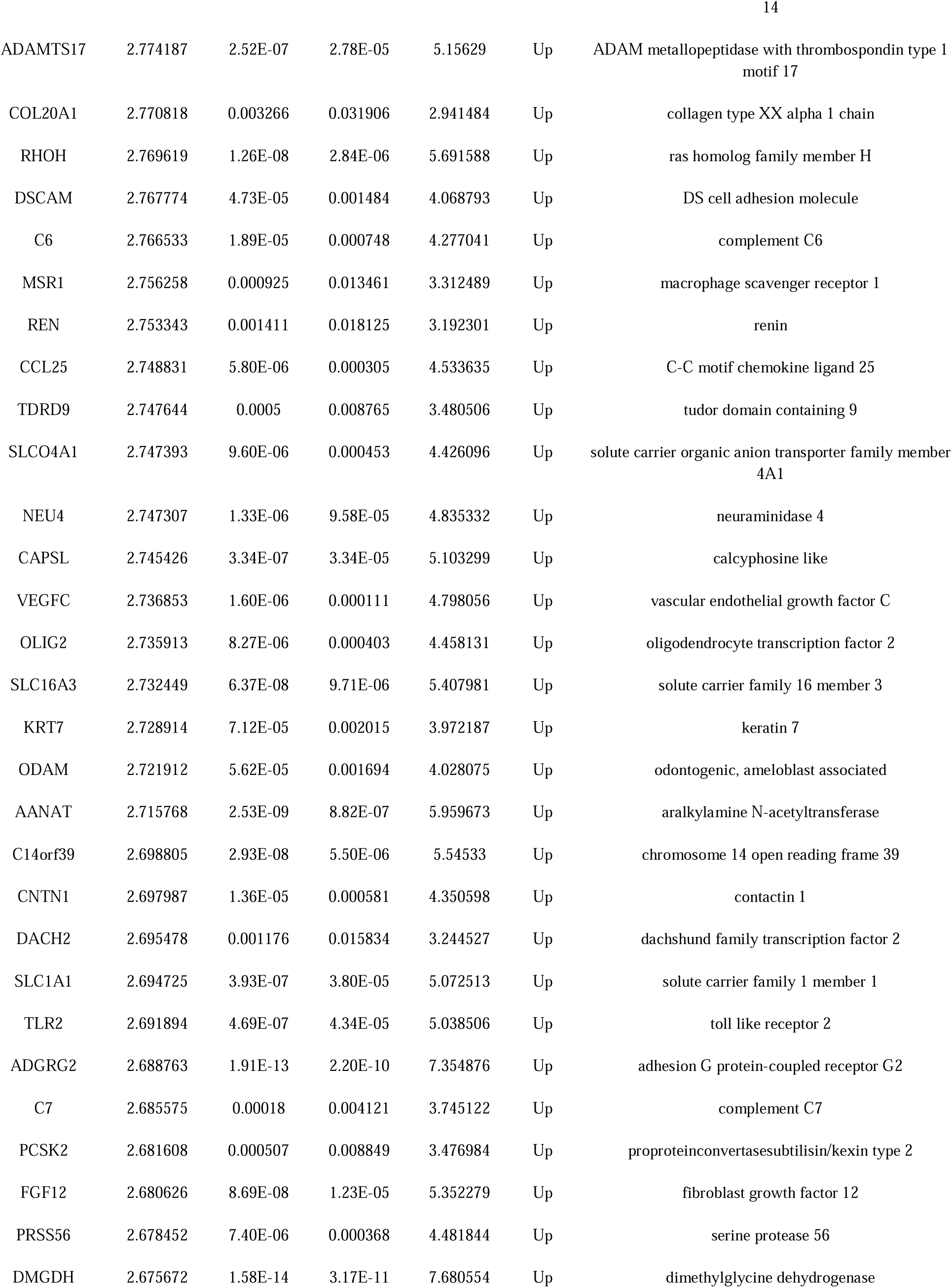
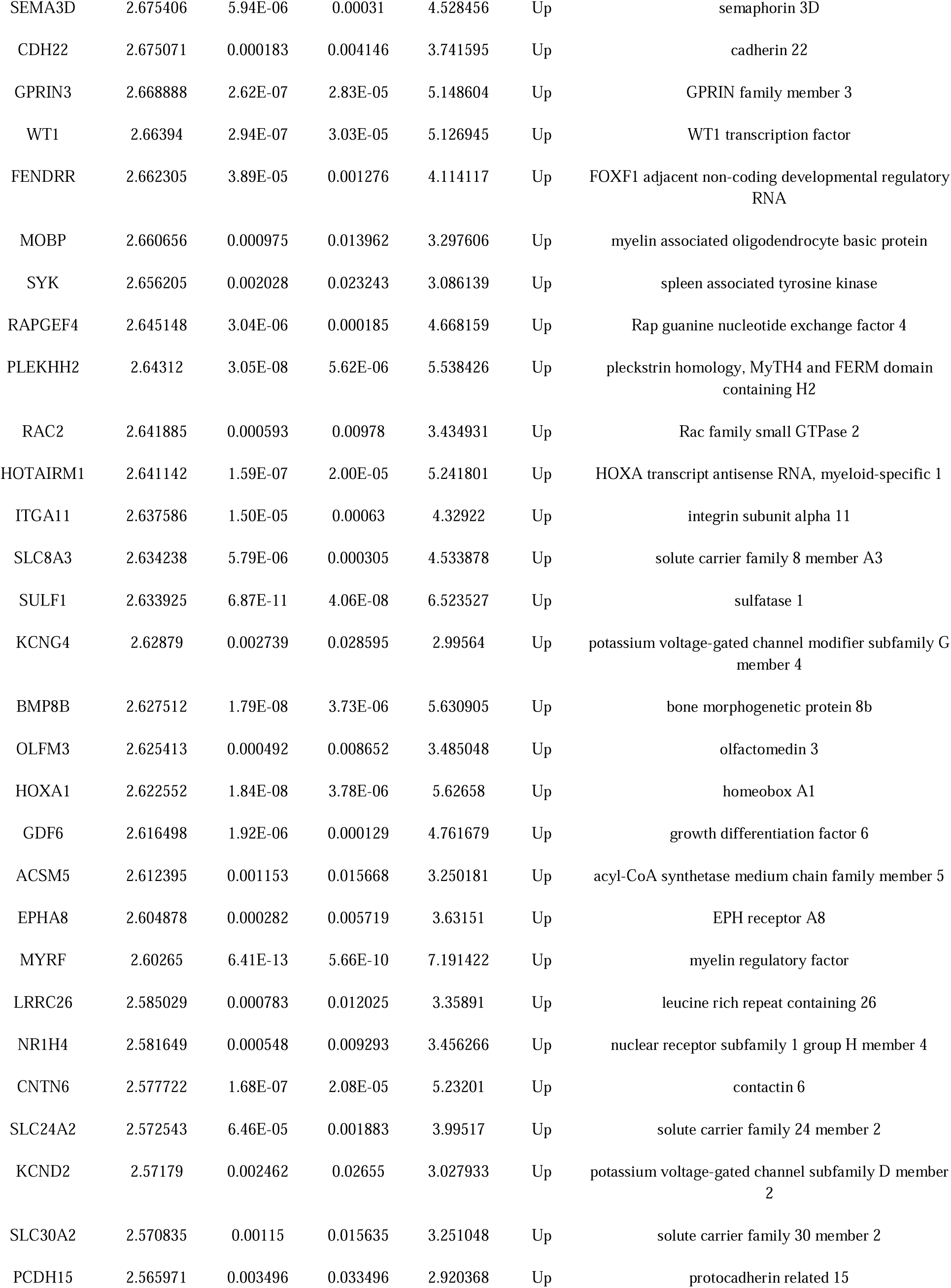
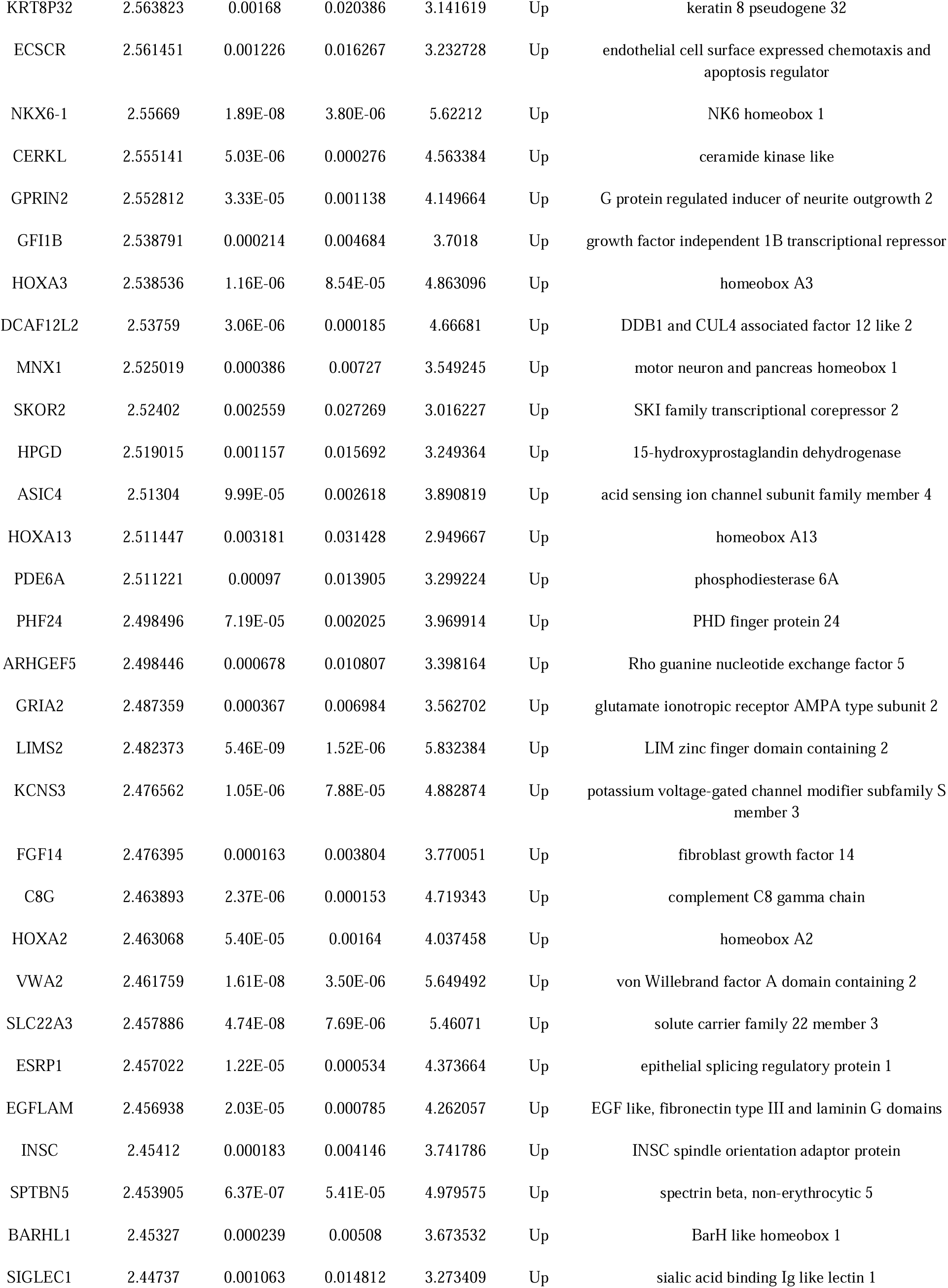
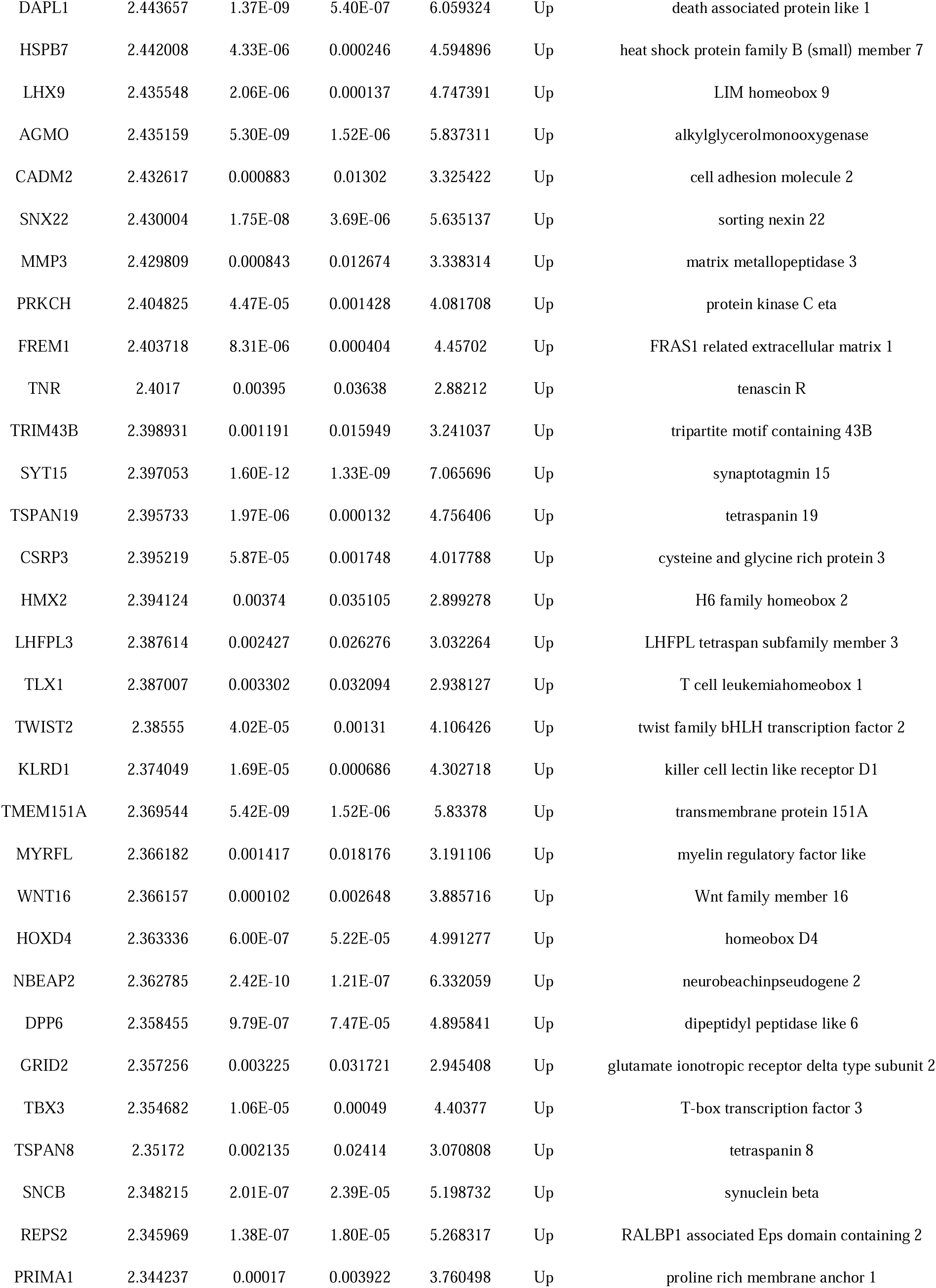
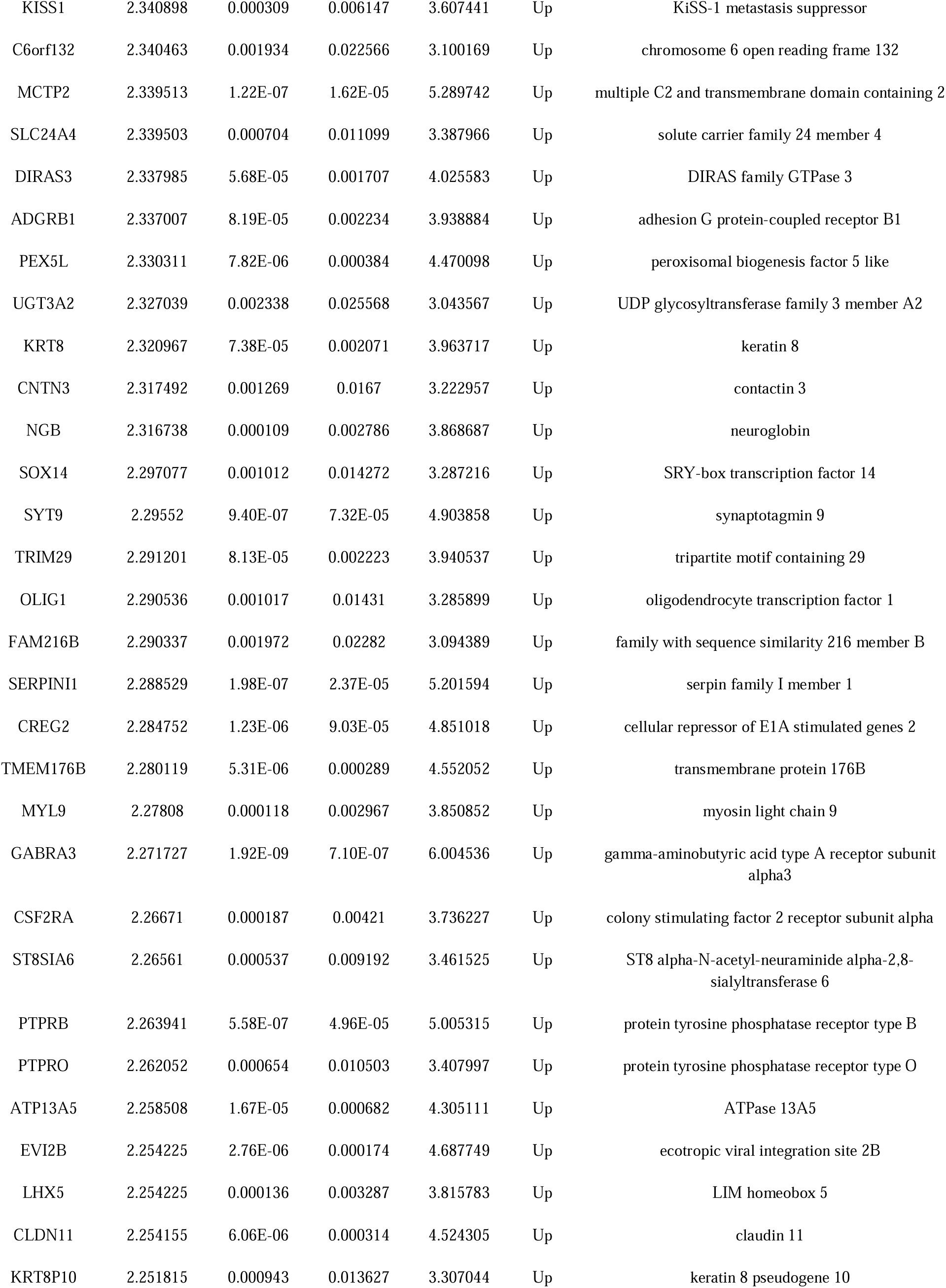
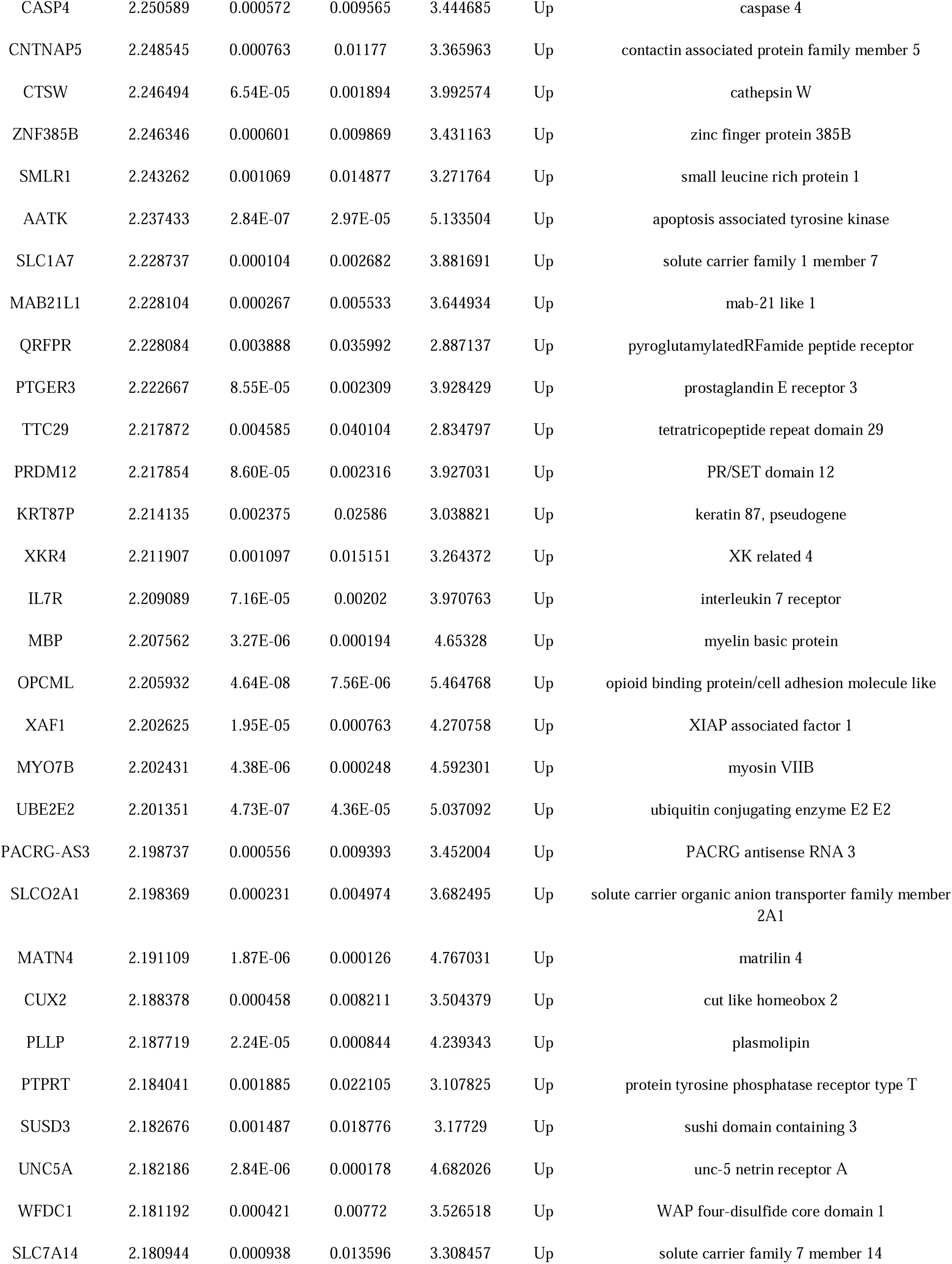
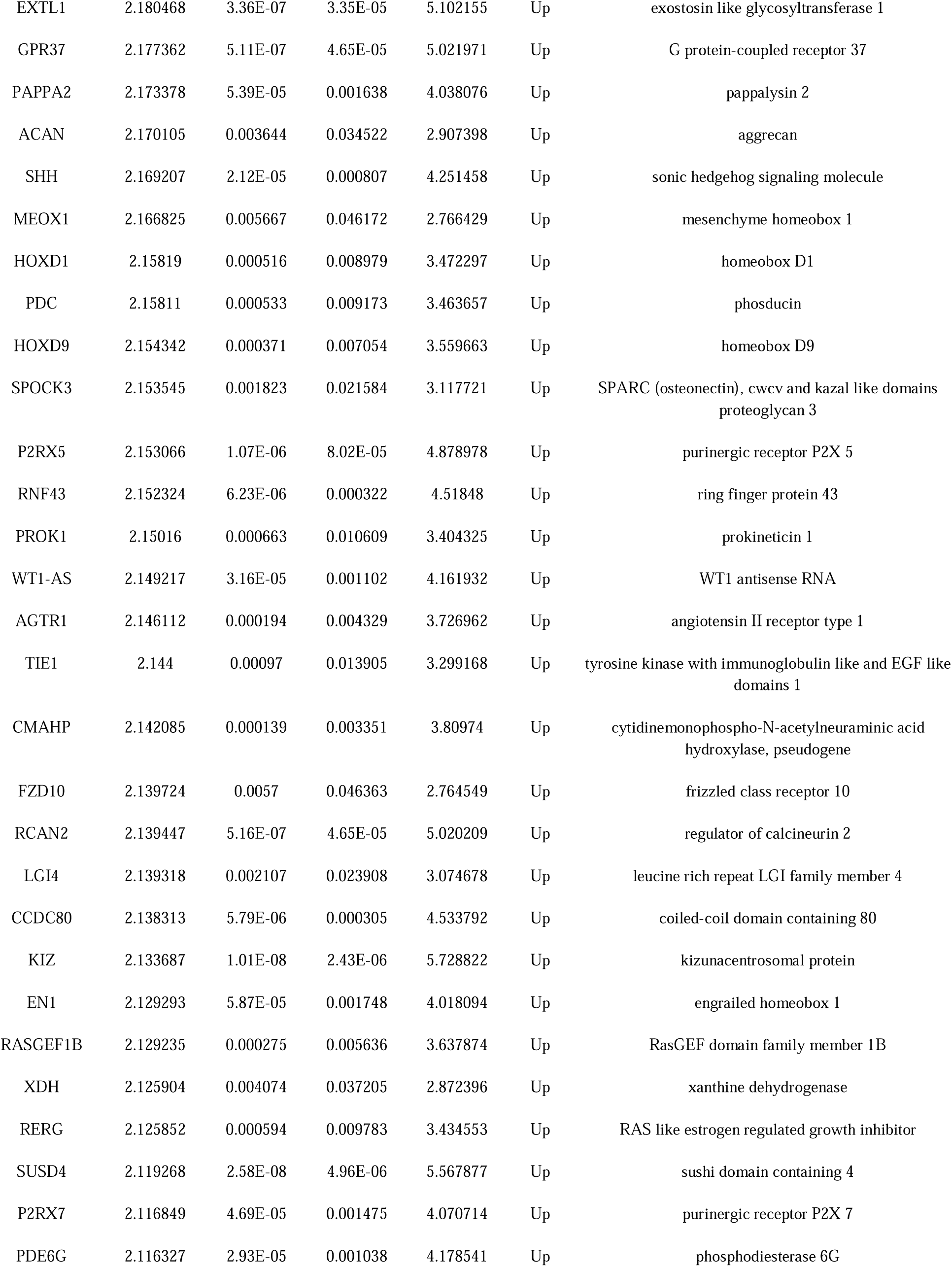
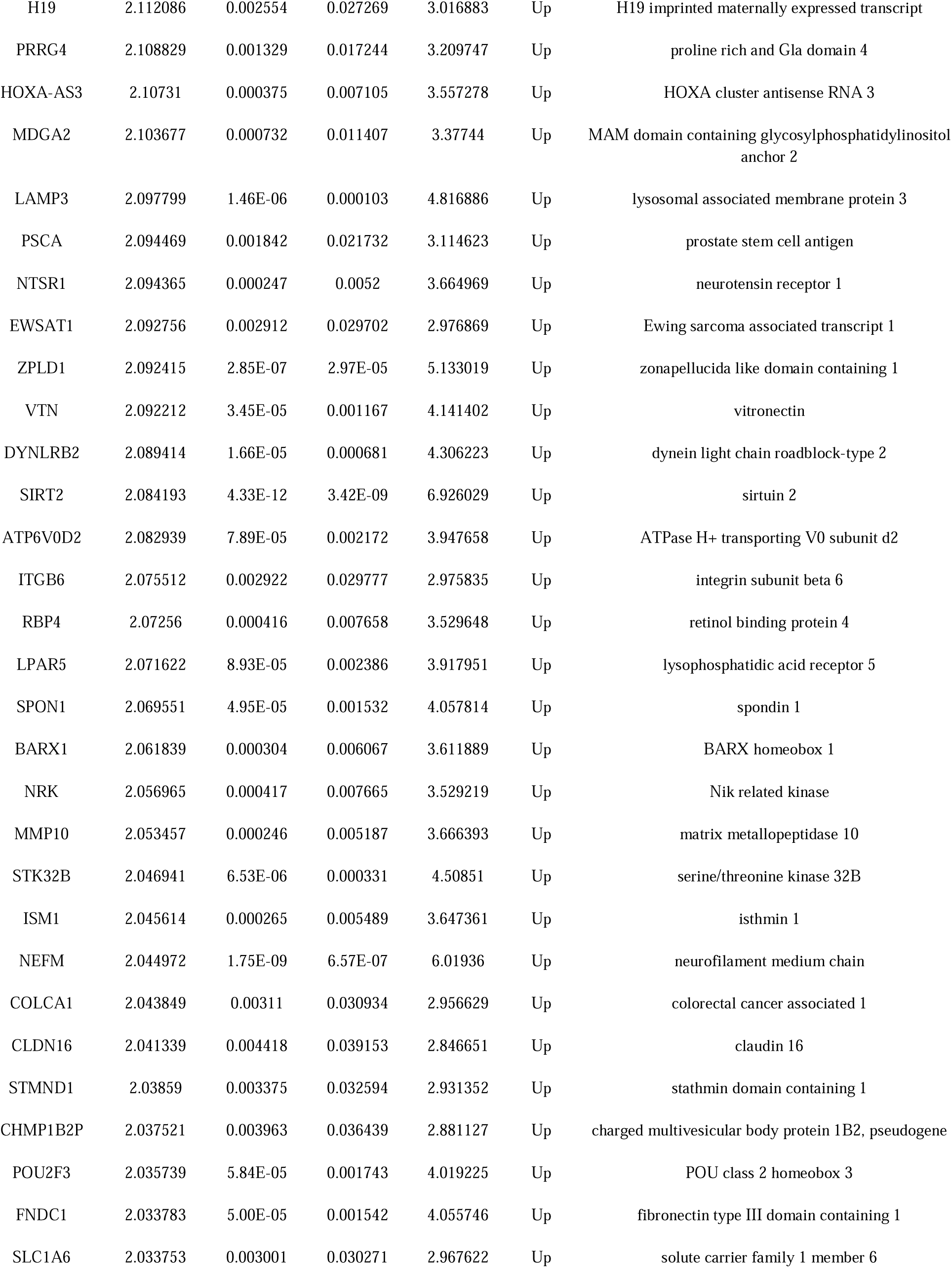
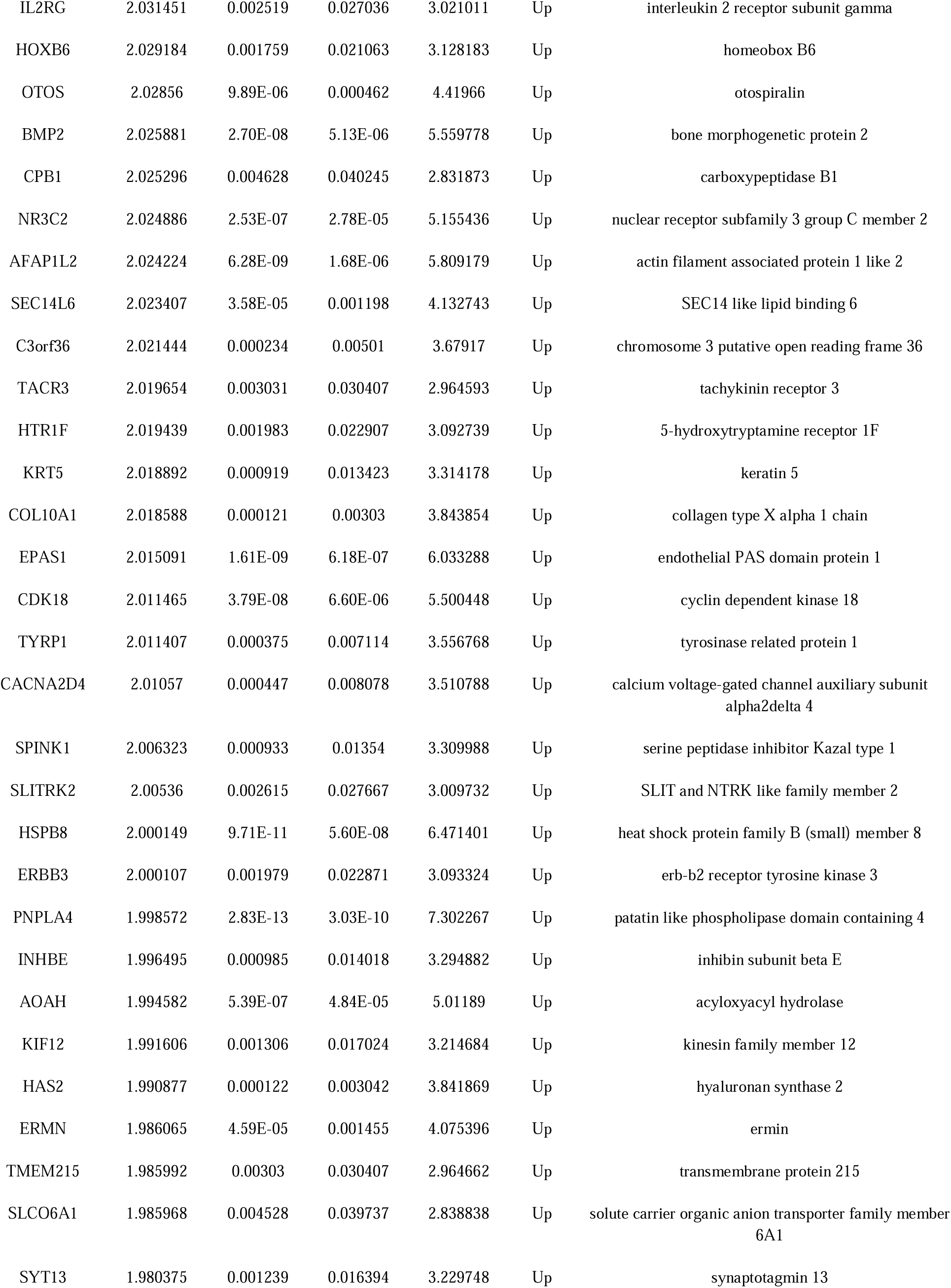
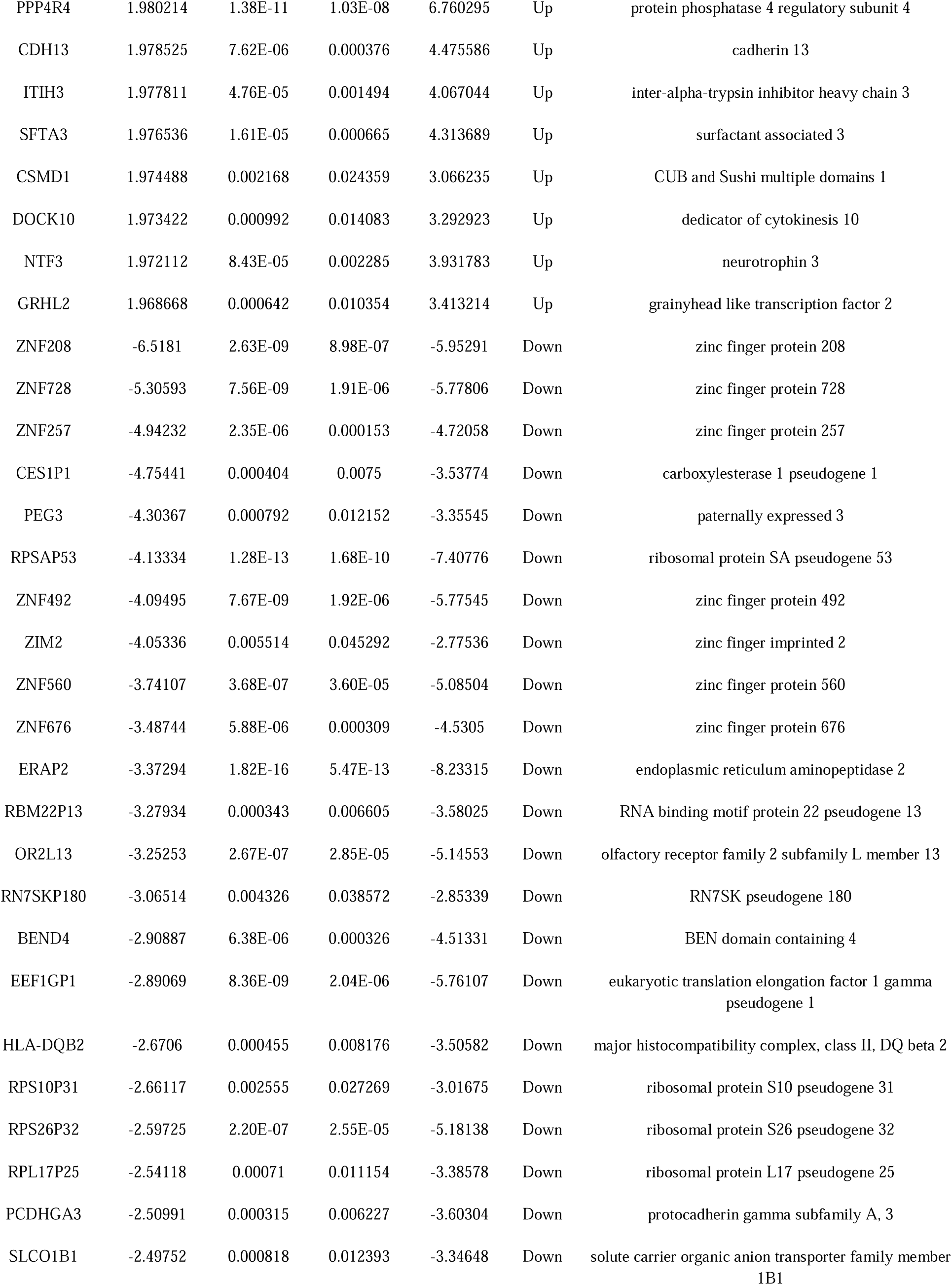
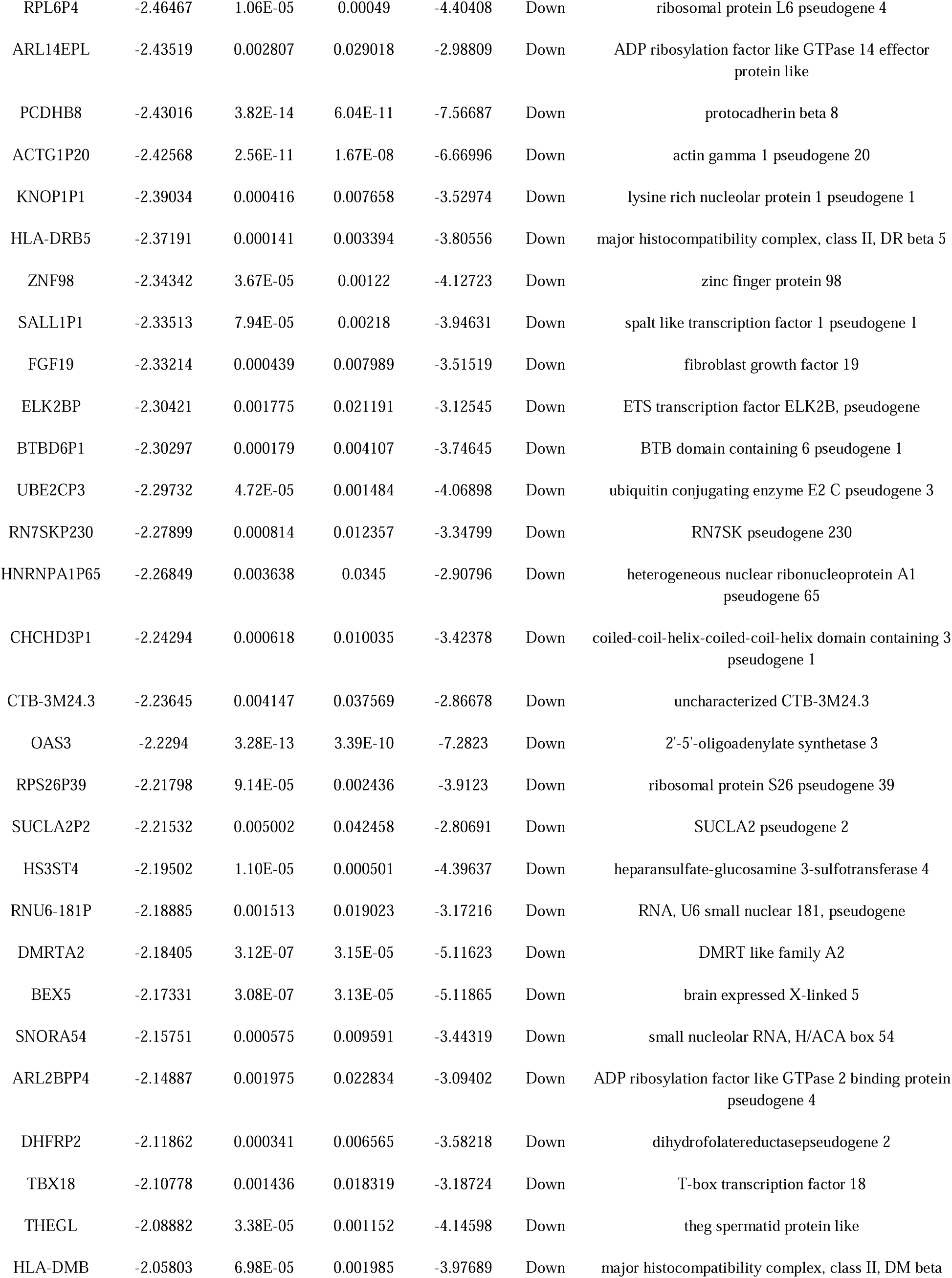
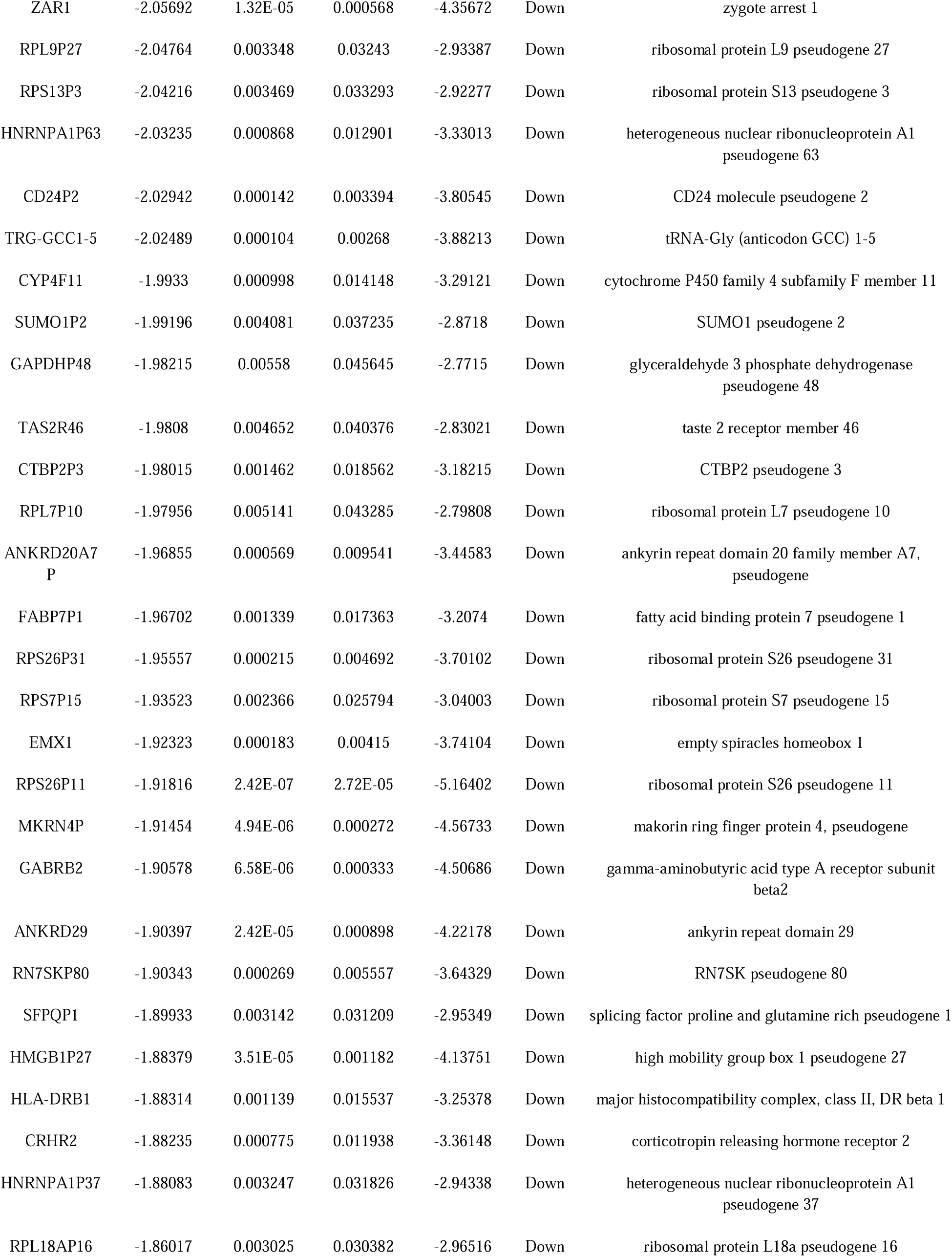
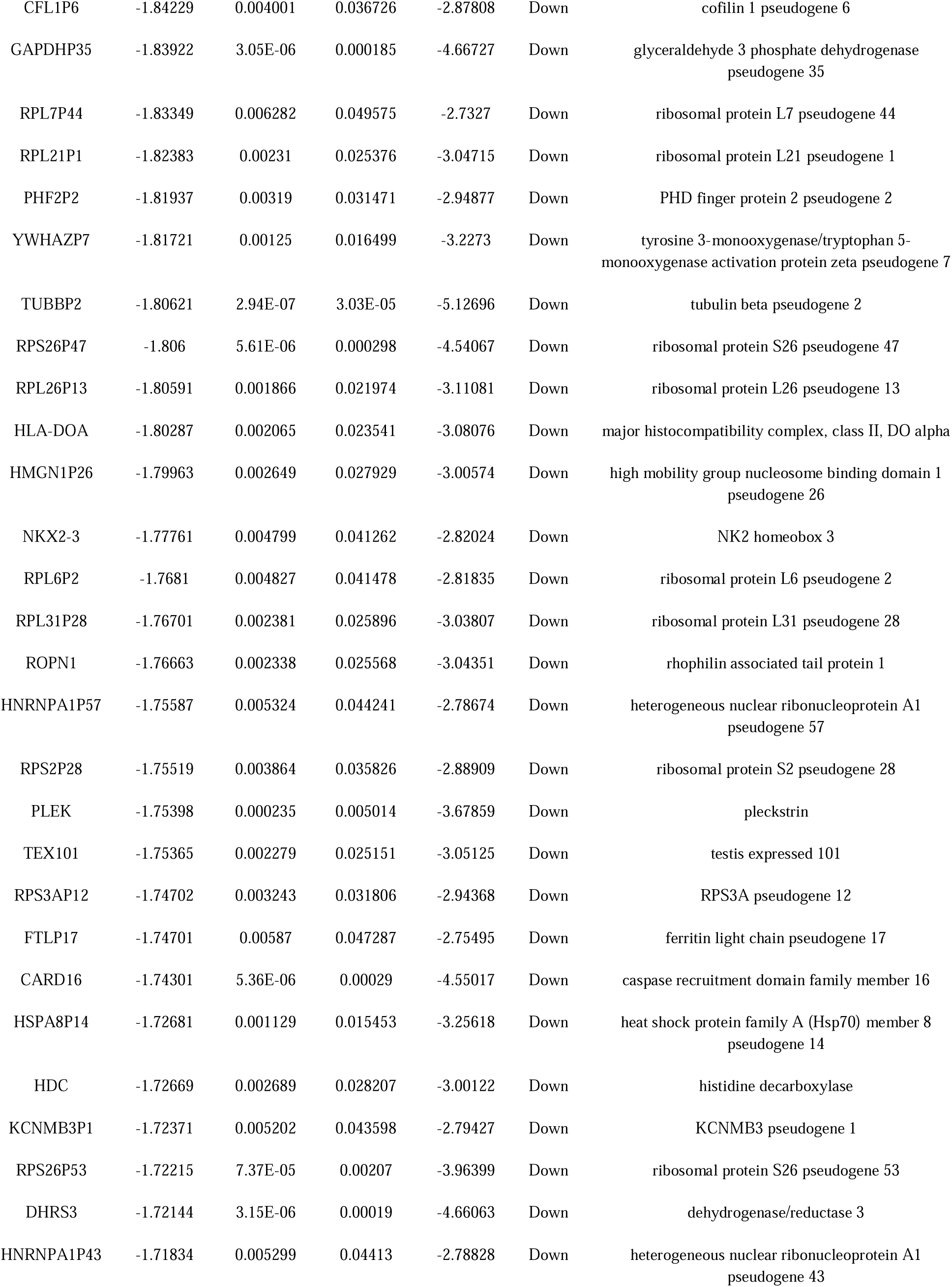
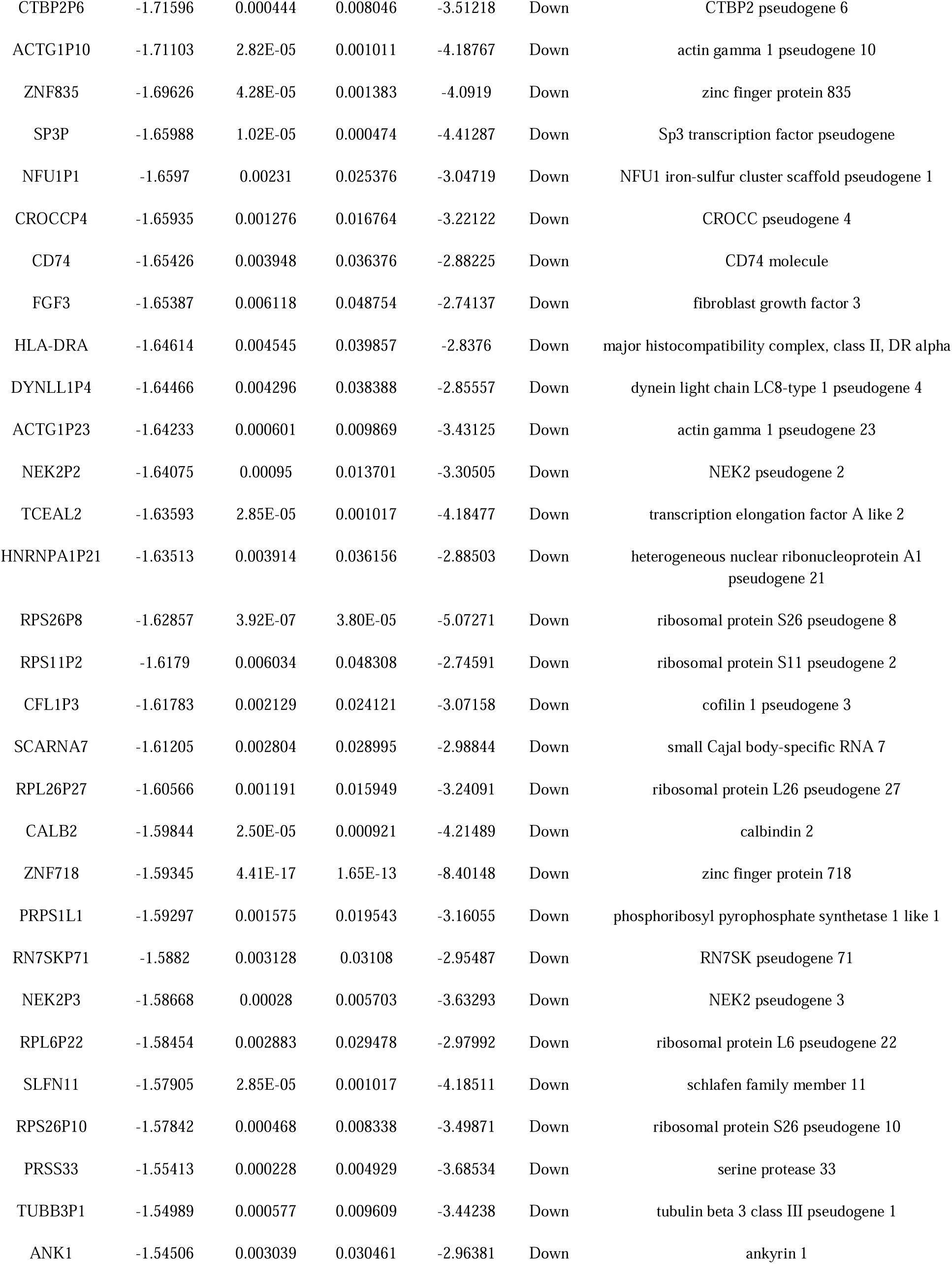
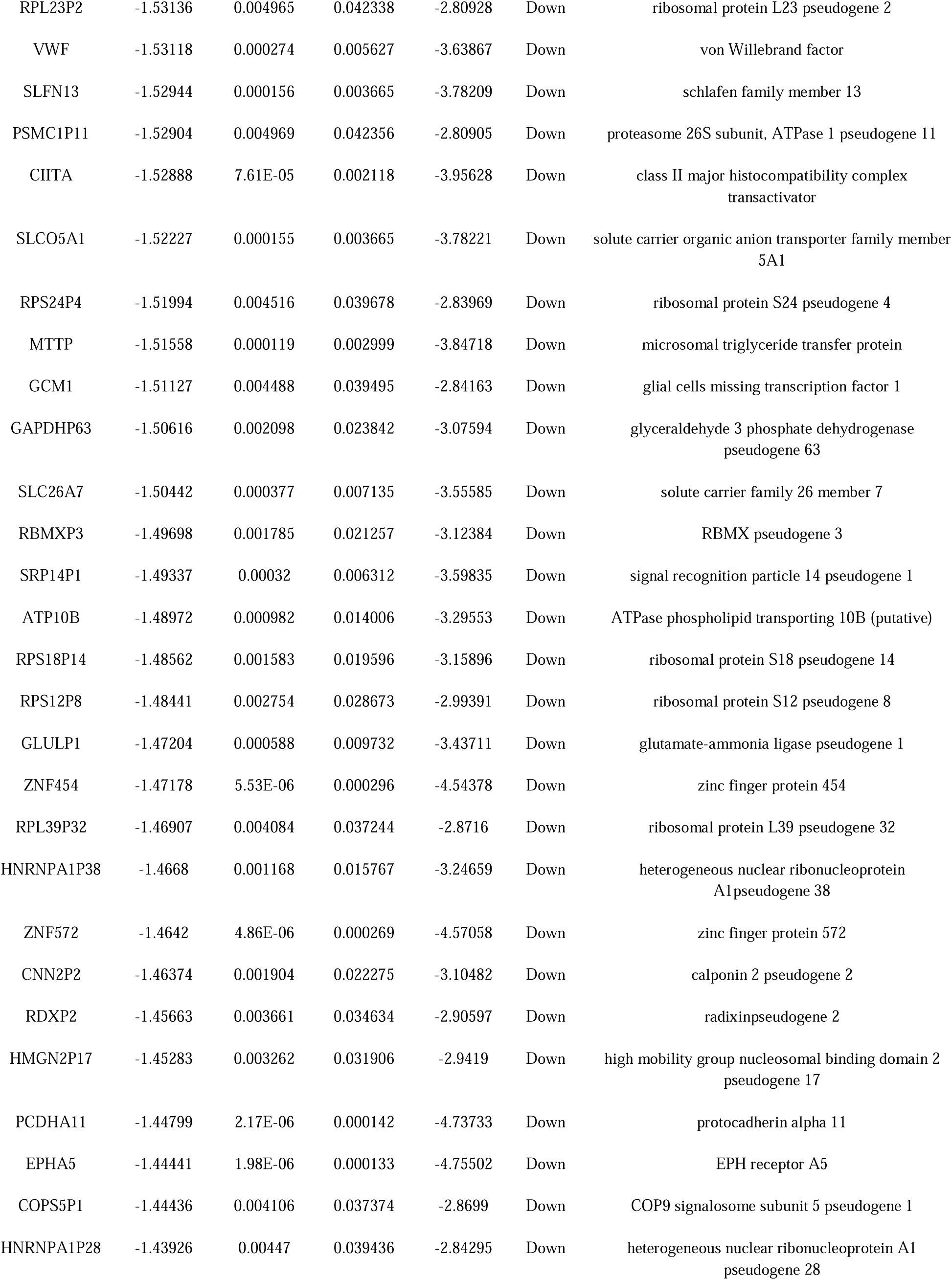
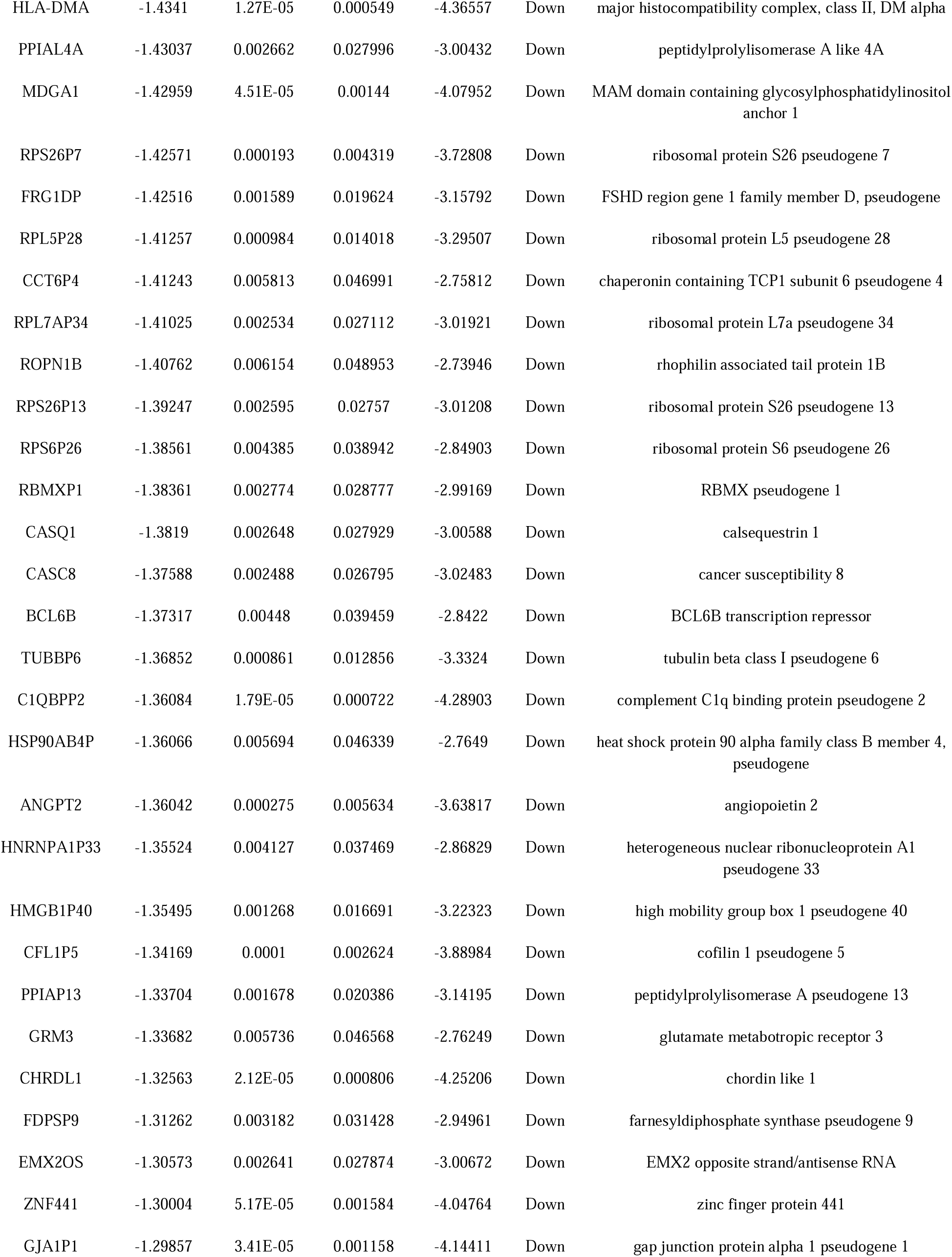
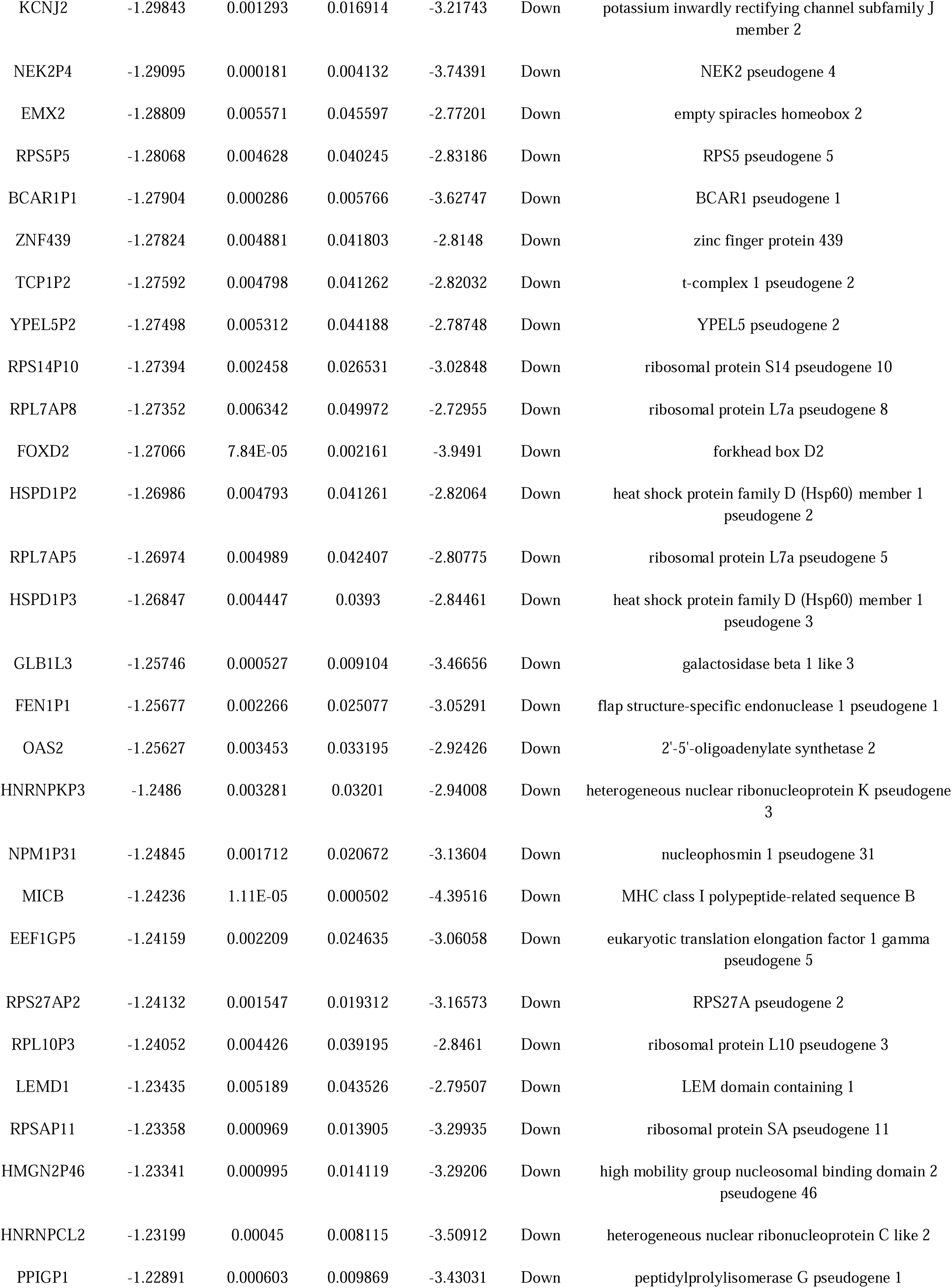
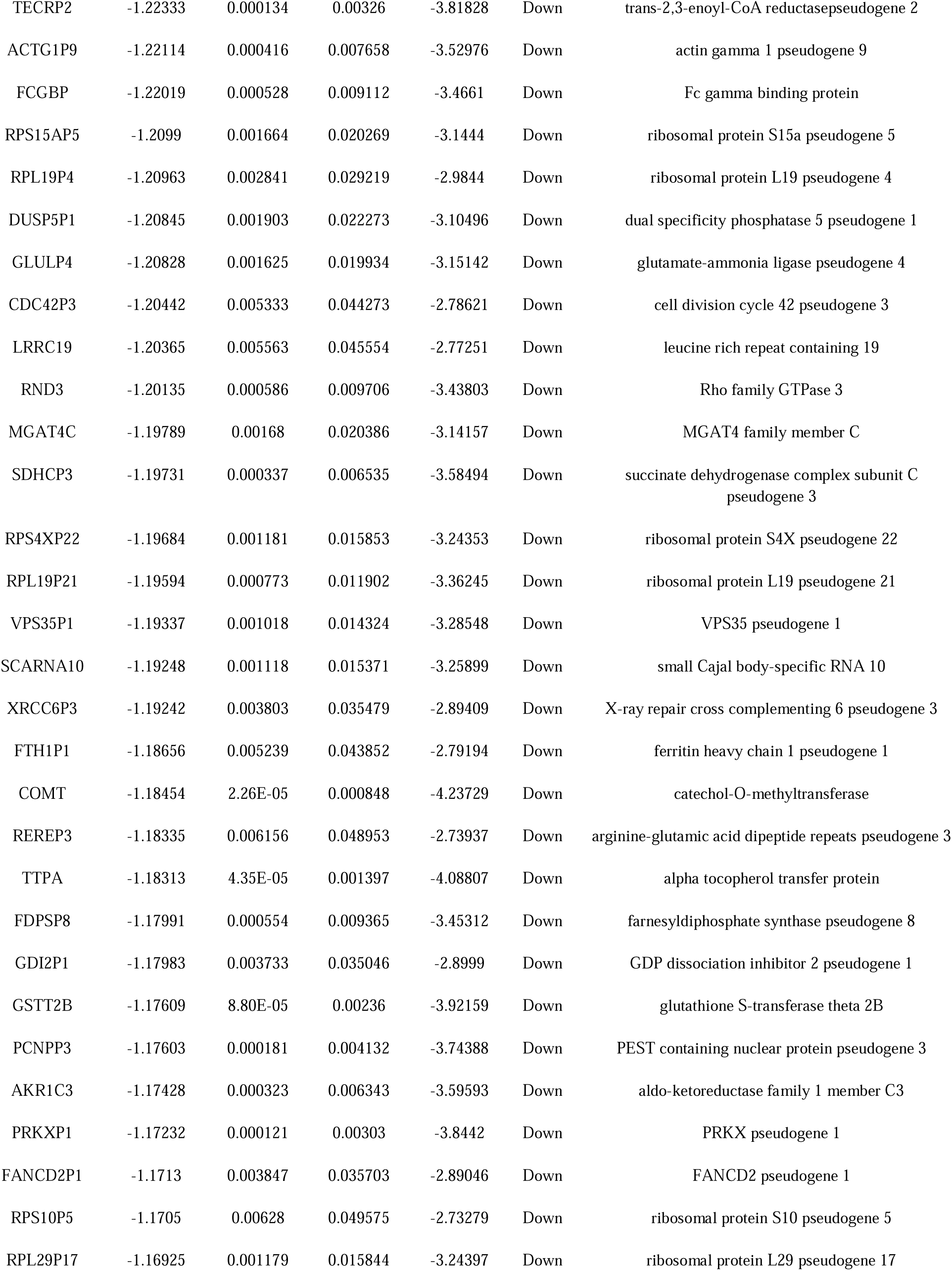
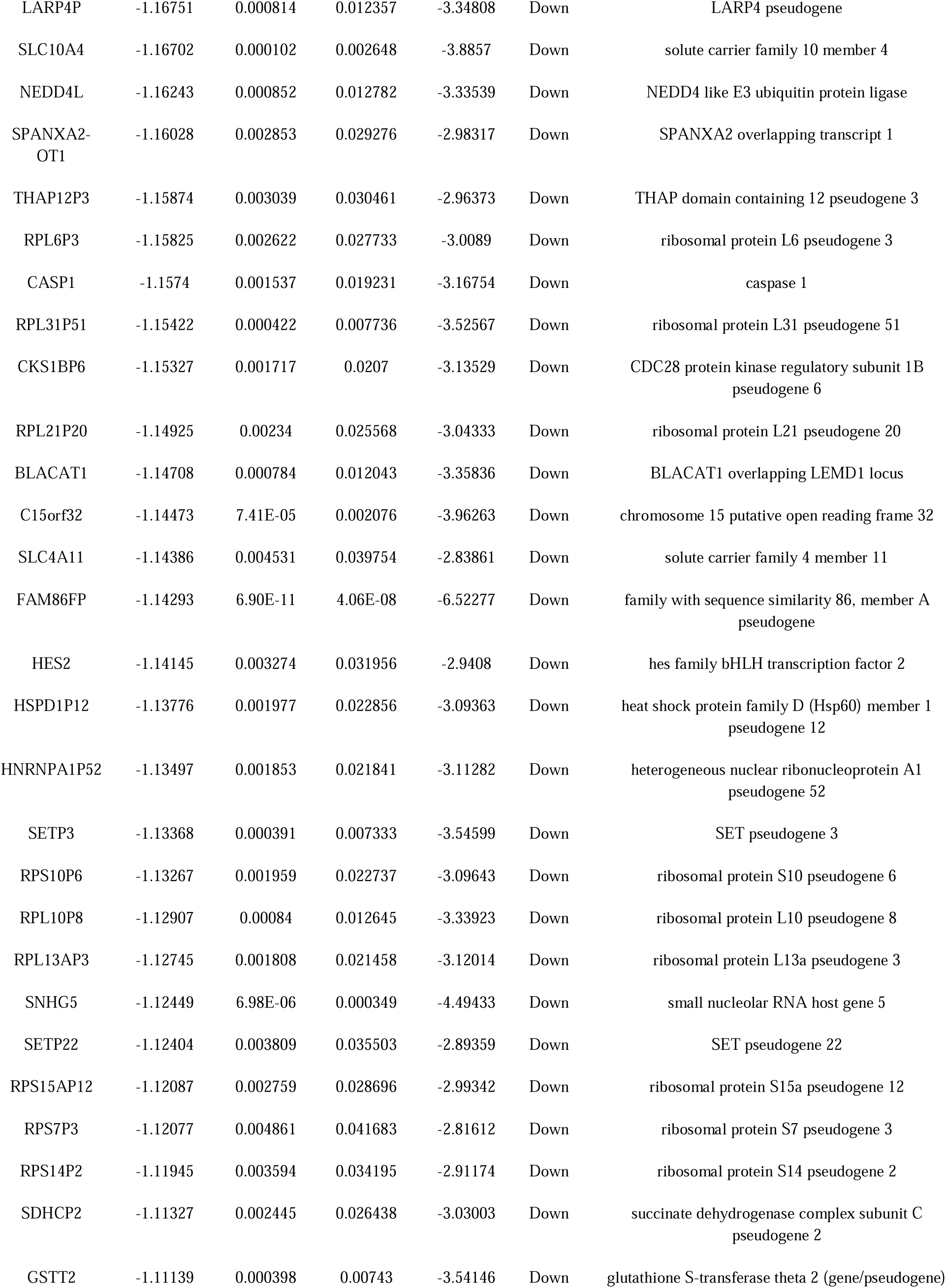
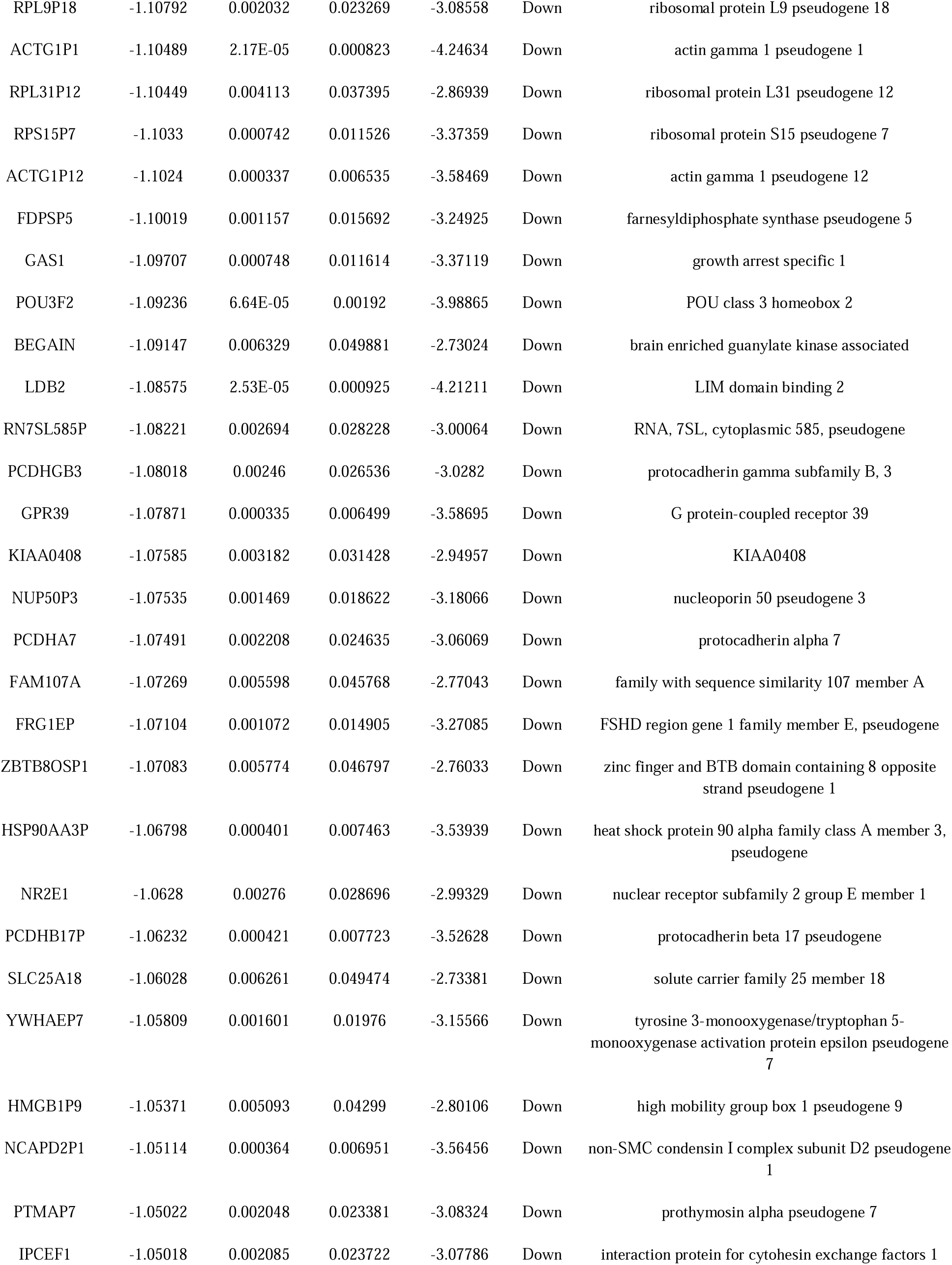
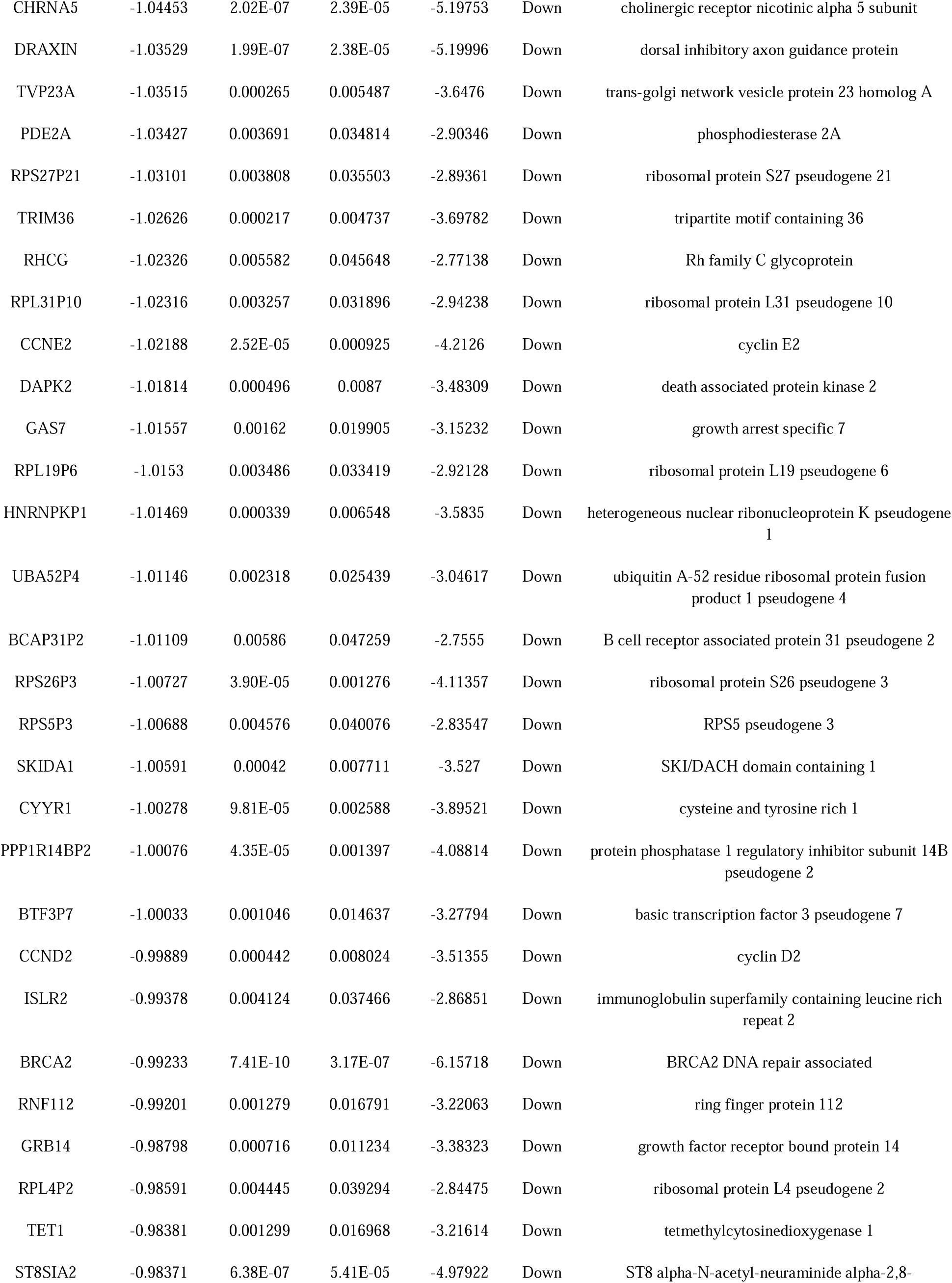
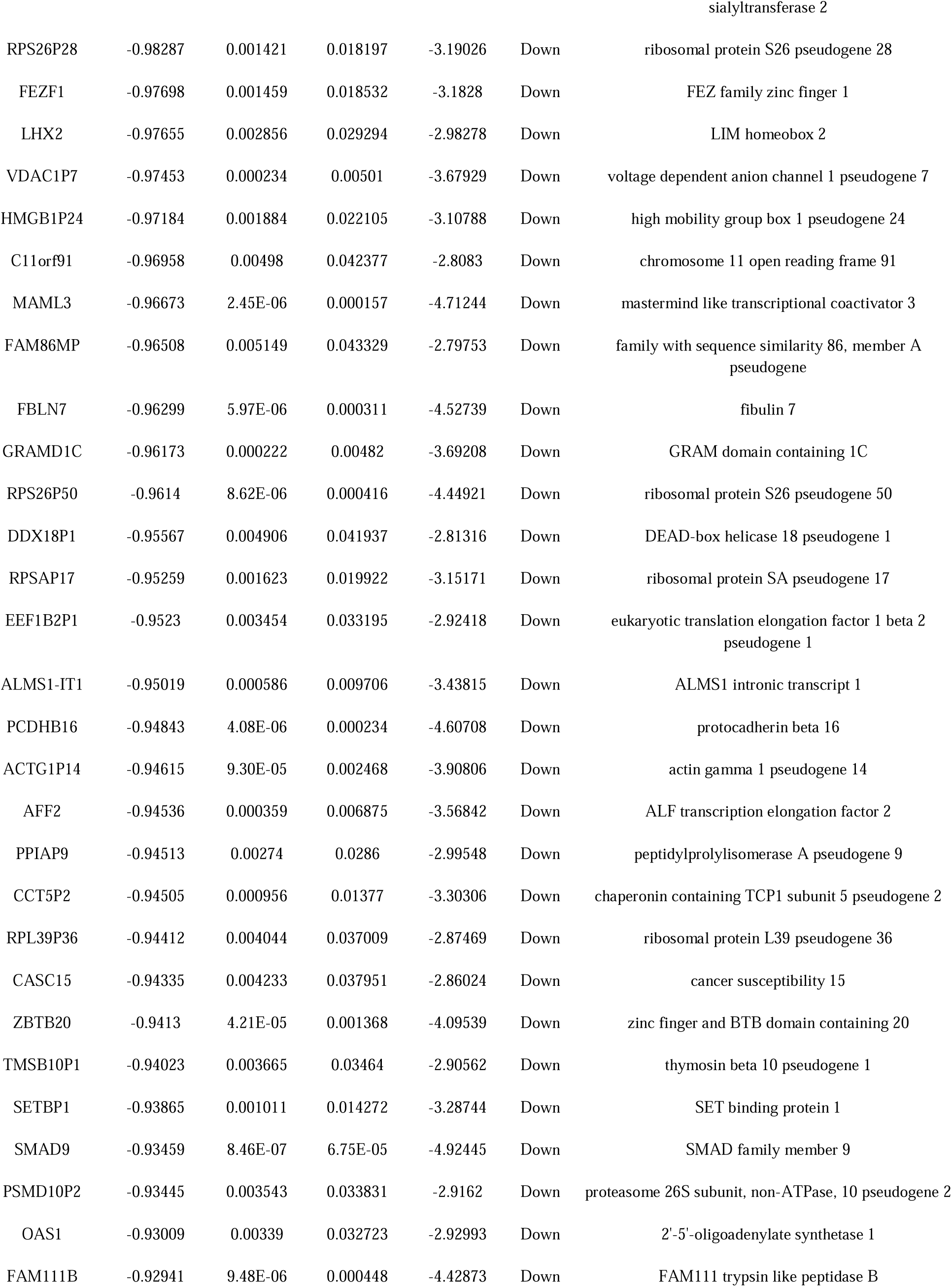
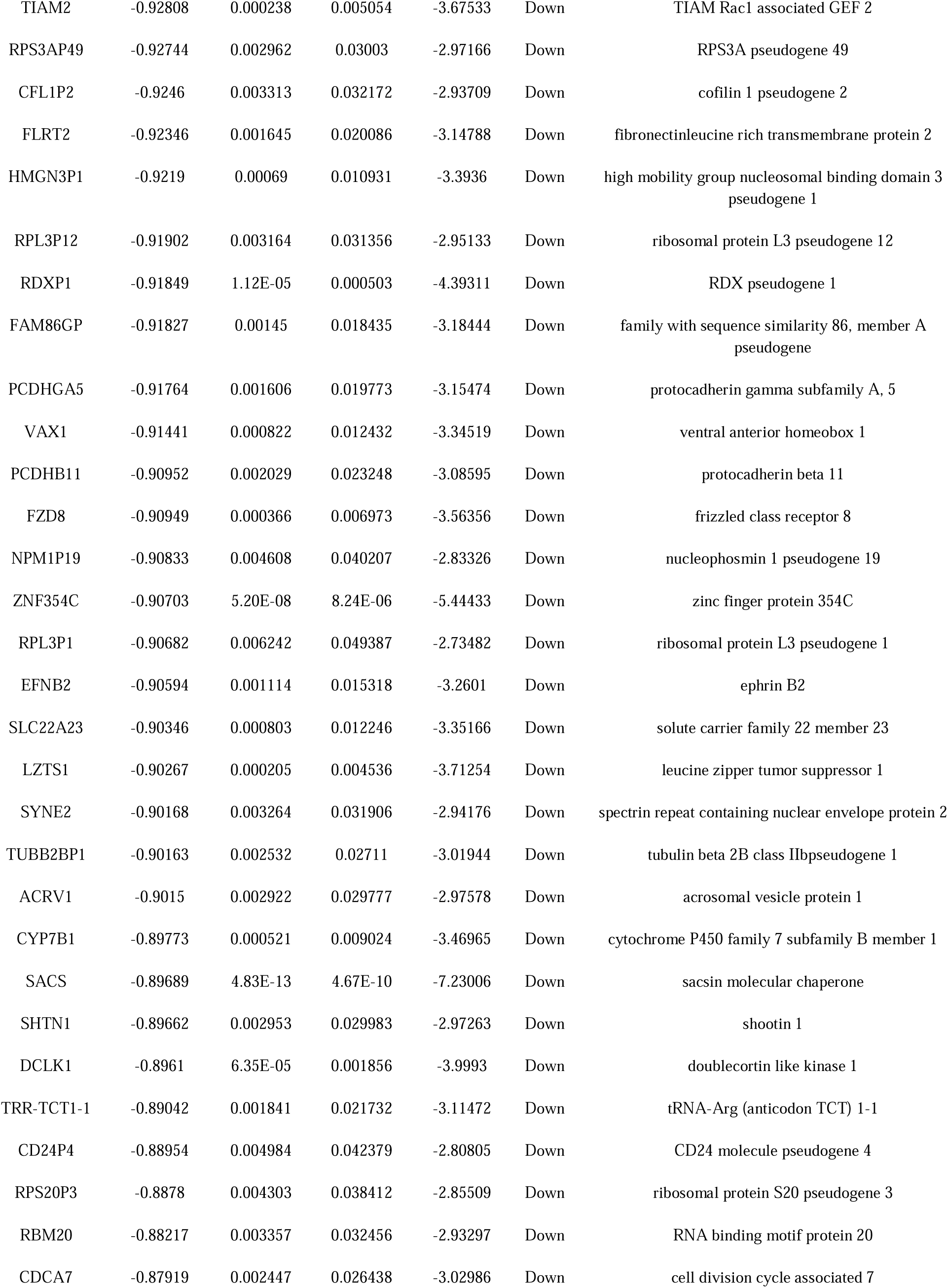
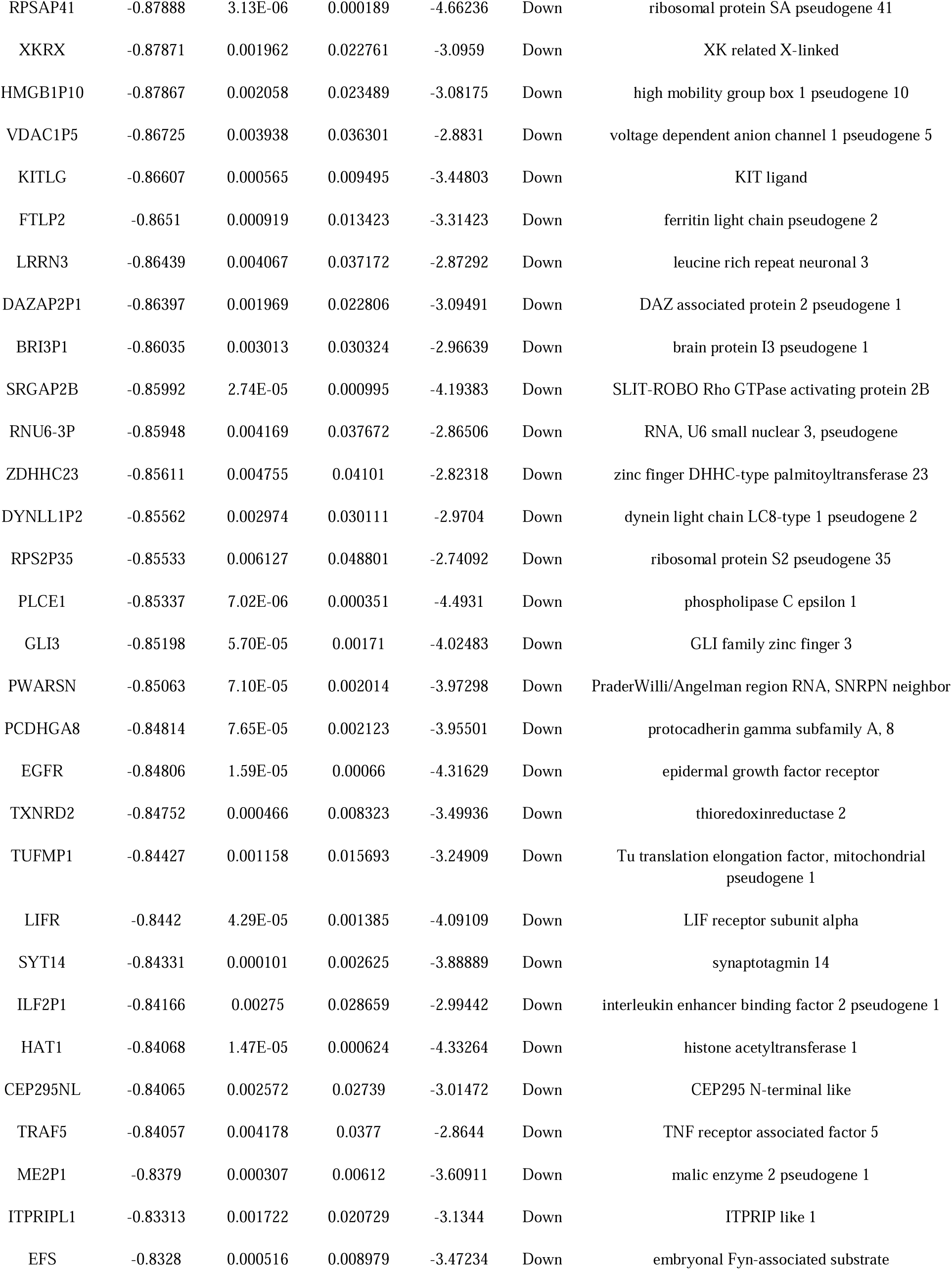
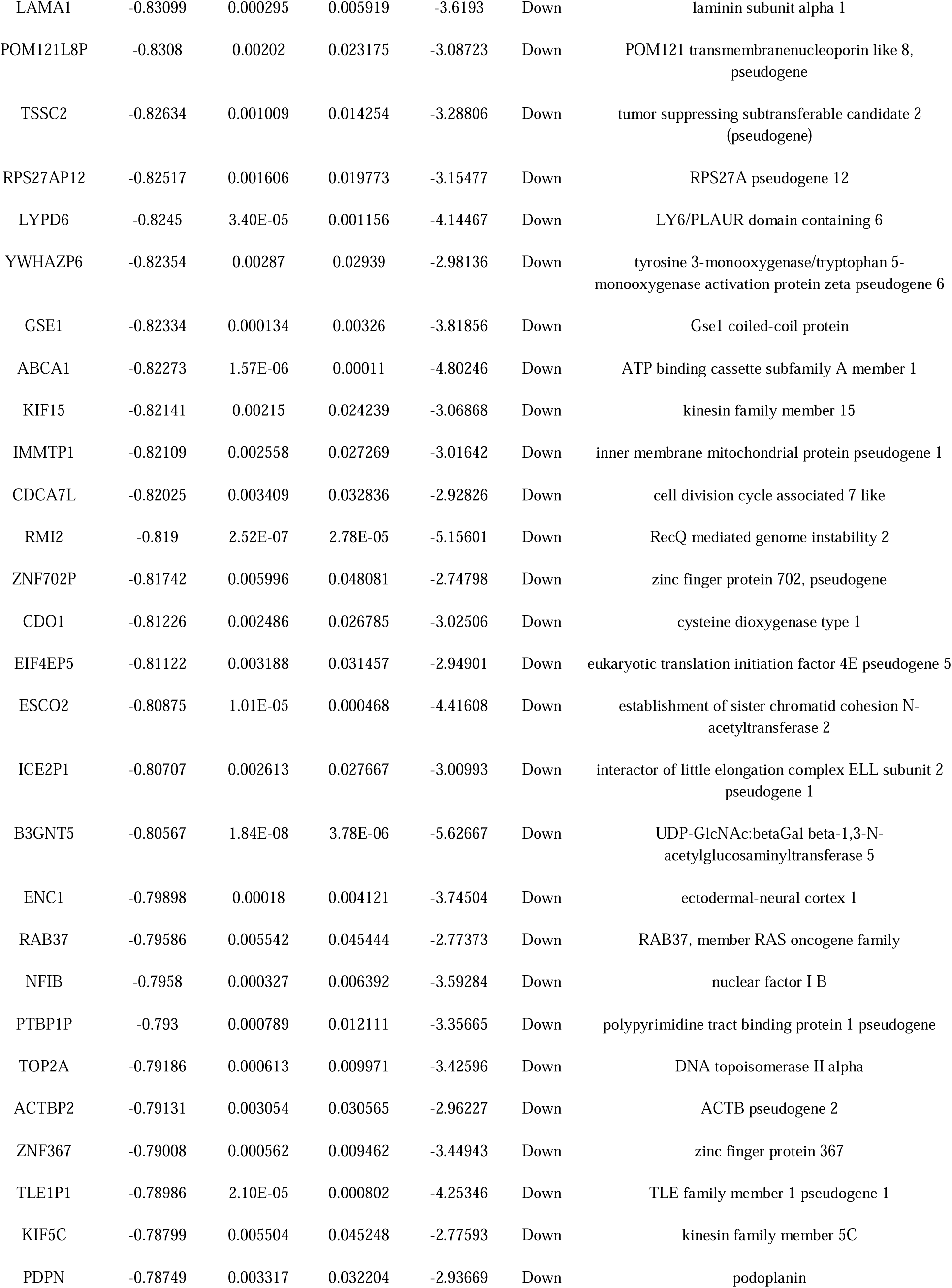
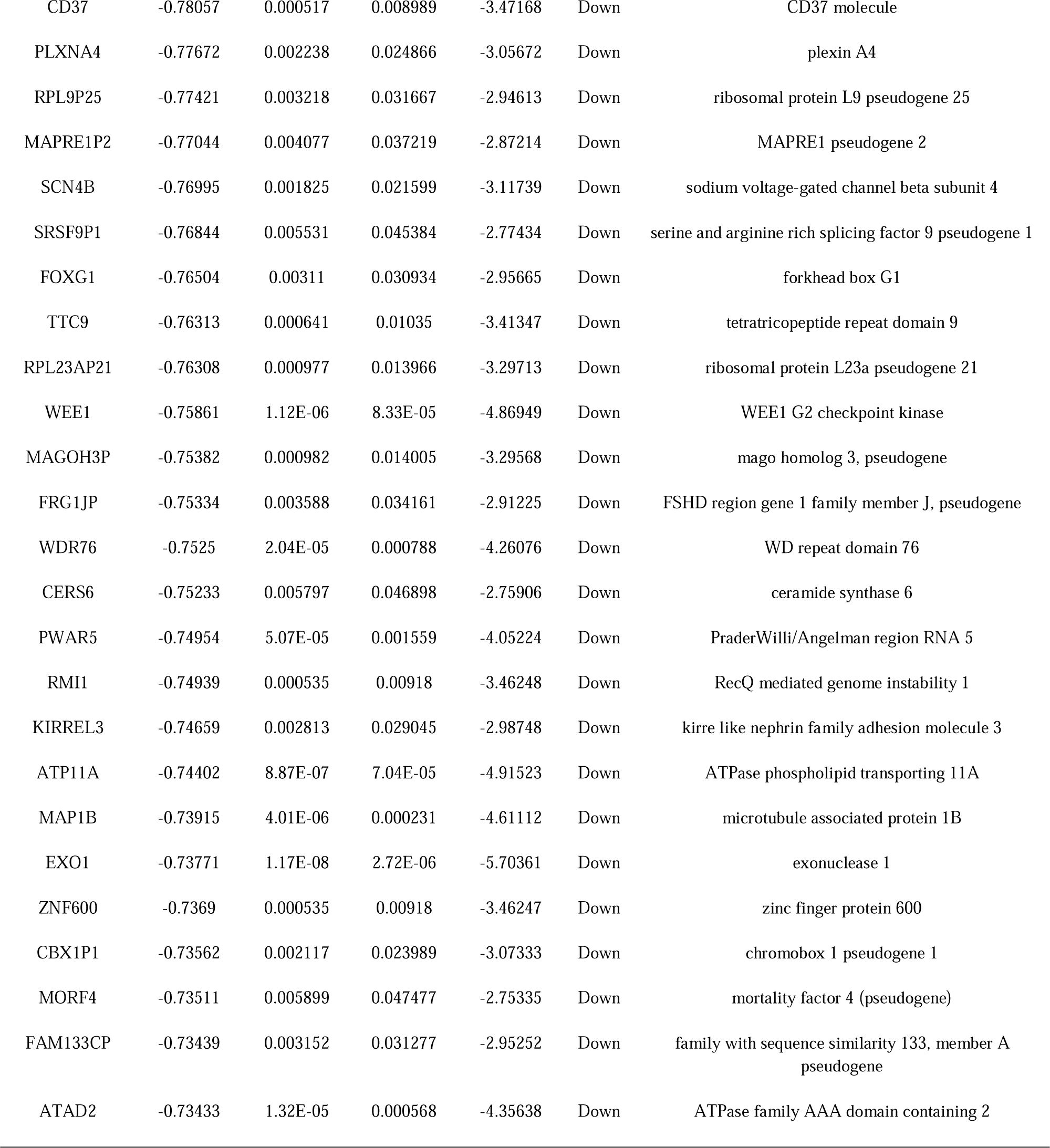
The statistical metrics for key differentially expressed genes (DEGs)

**Fig. 1.**
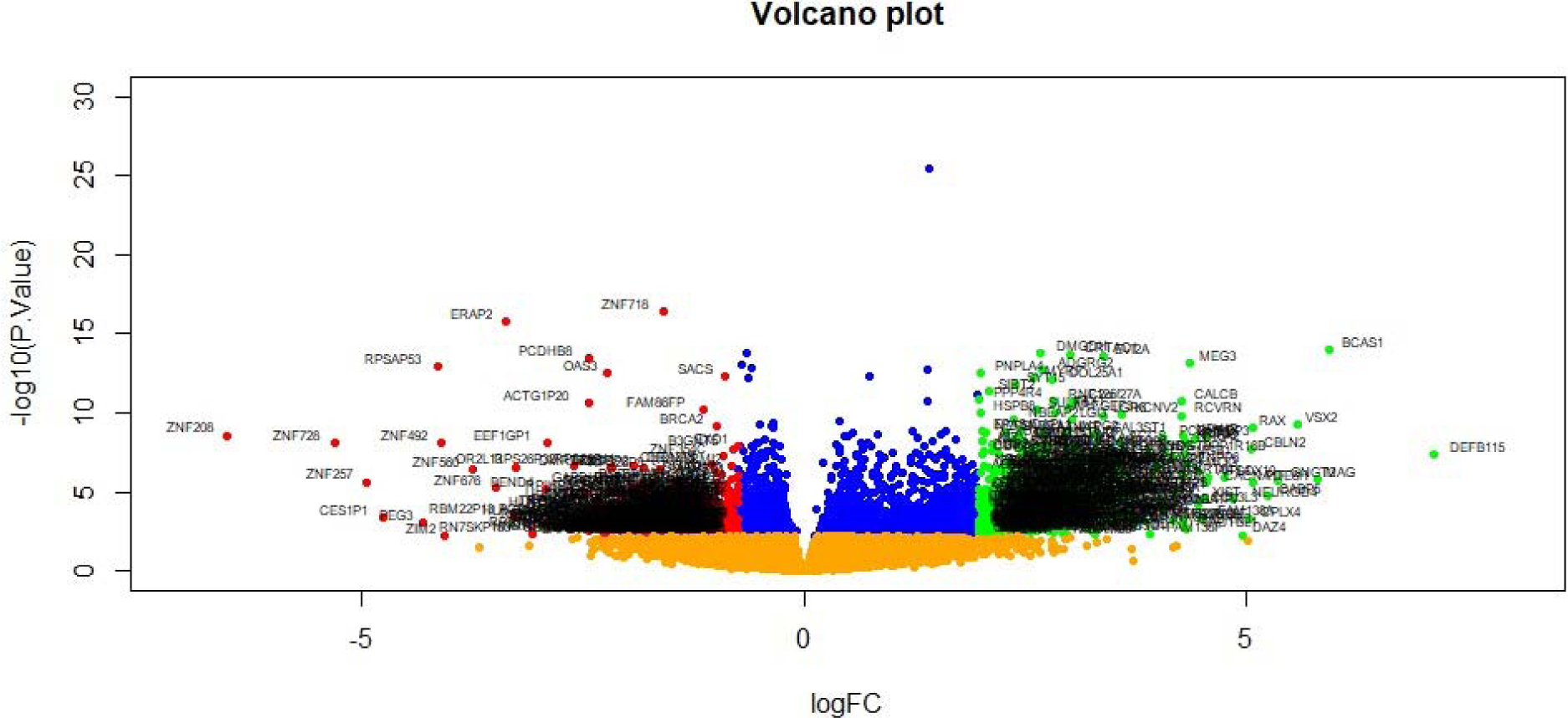
Volcano plot of differentially expressed genes. Genes with a significant change of more than two-fold were selected. Green dot represented up regulated significant genes and red dot represented down regulated significant genes.

**Fig. 2.**
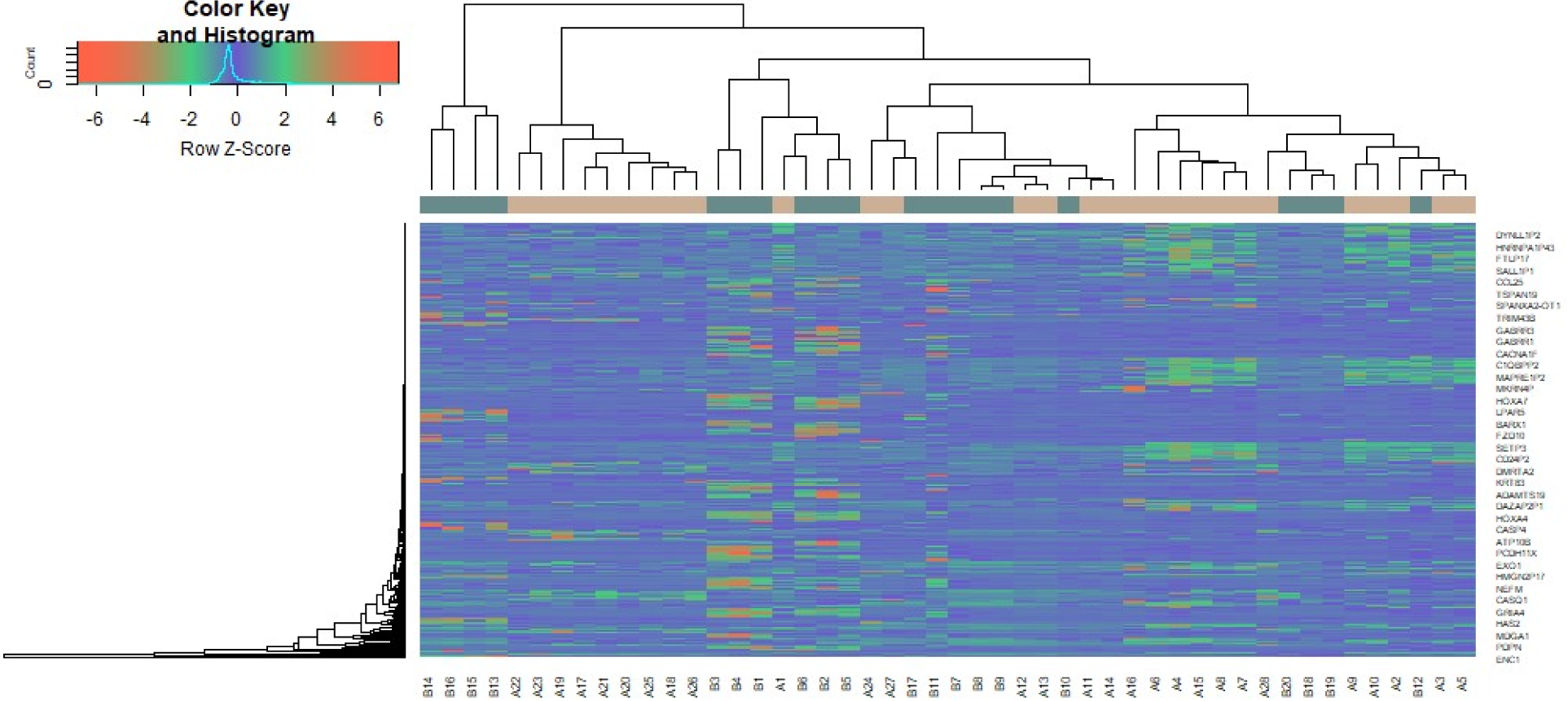
Heat map of differentially expressed genes. Legend on the top left indicate log fold change of genes. (A1 – A28 = HD samples; B1 – B20 = Normal control samples)

### GO and pathway enrichment analyses of DEGs

To obtain a deeper insight into the biological functions of DEGs, GO annotation and REACTOME pathway enrichment analyses were performed. The top enriched GO terms were shown in Table 2. The GO terms were comprised of 3 parts: BP, CC and MF. DEGs of BP were involved in multicellular organismal process, biological regulation and developmental process. CC analysis revealed that DEGs were markedly enriched neuron projection, membrane, cell periphery and cellular anatomical entity. For MF analysis, the top significantly enriched terms were DNA-binding transcription factor activity, molecular transducer activity, ion binding and protein binding. Besides, the enriched REACTOME pathways as presented in Table 3, including signaling by GPCR, Class A/1 (Rhodopsin-like receptors, MHC class II antigen presentation and impaired BRCA2 binding to PALB2.

**Table 2.**
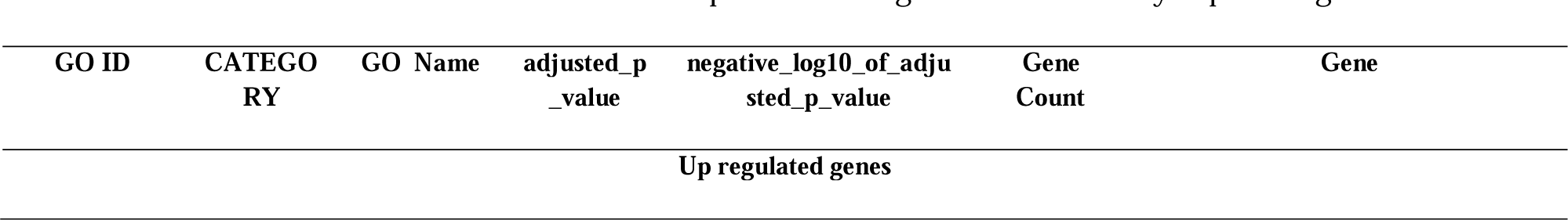
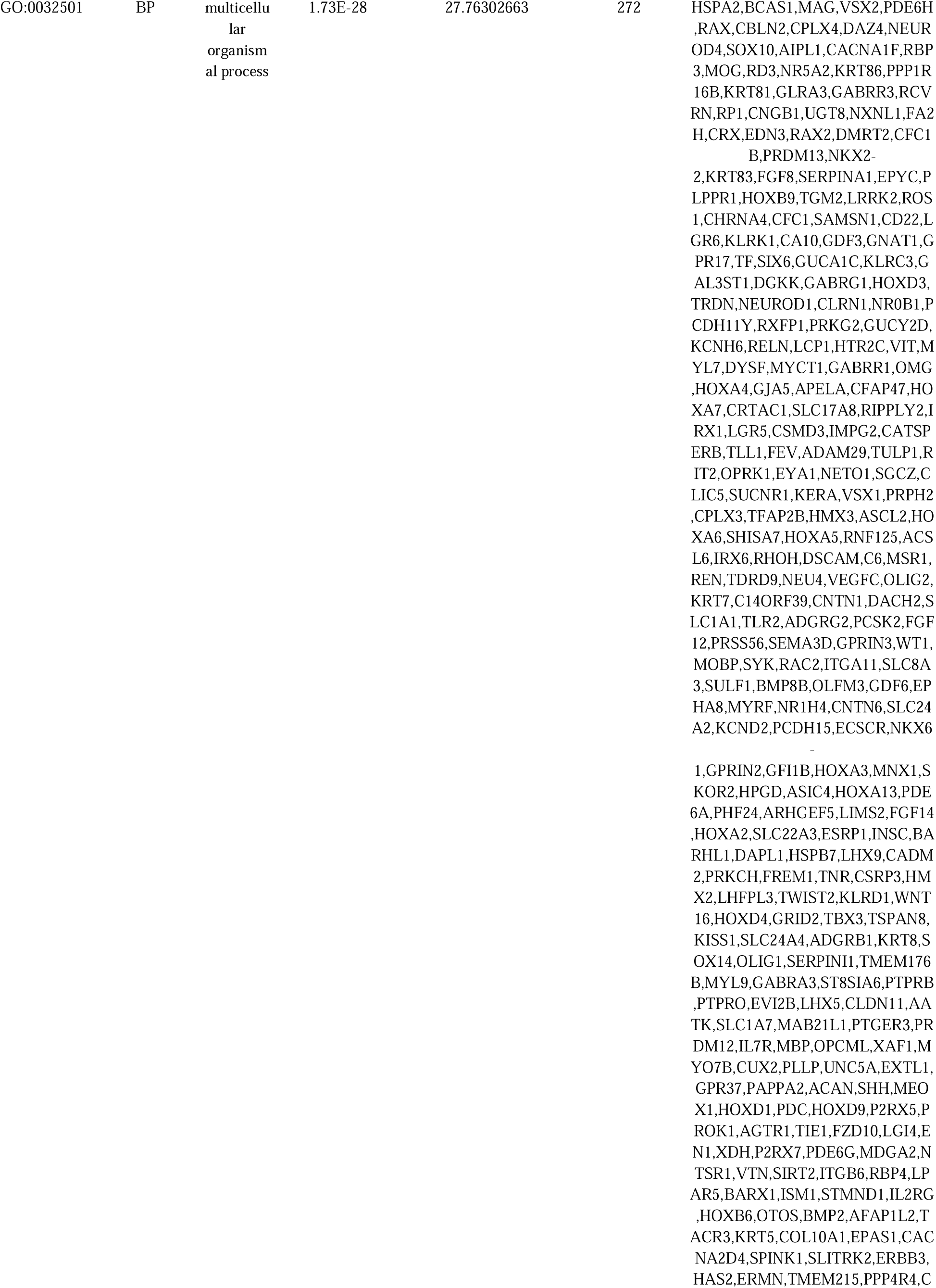
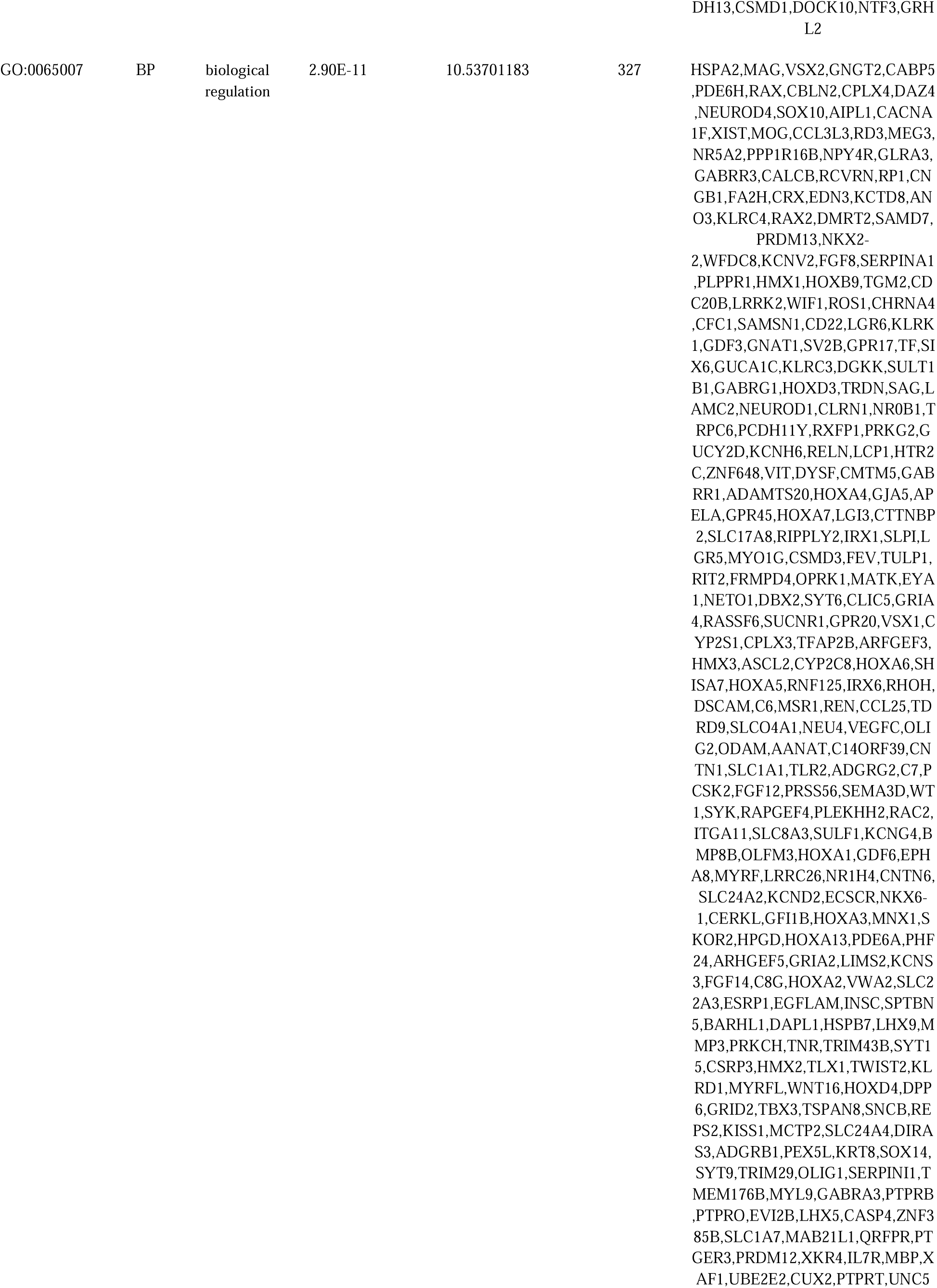
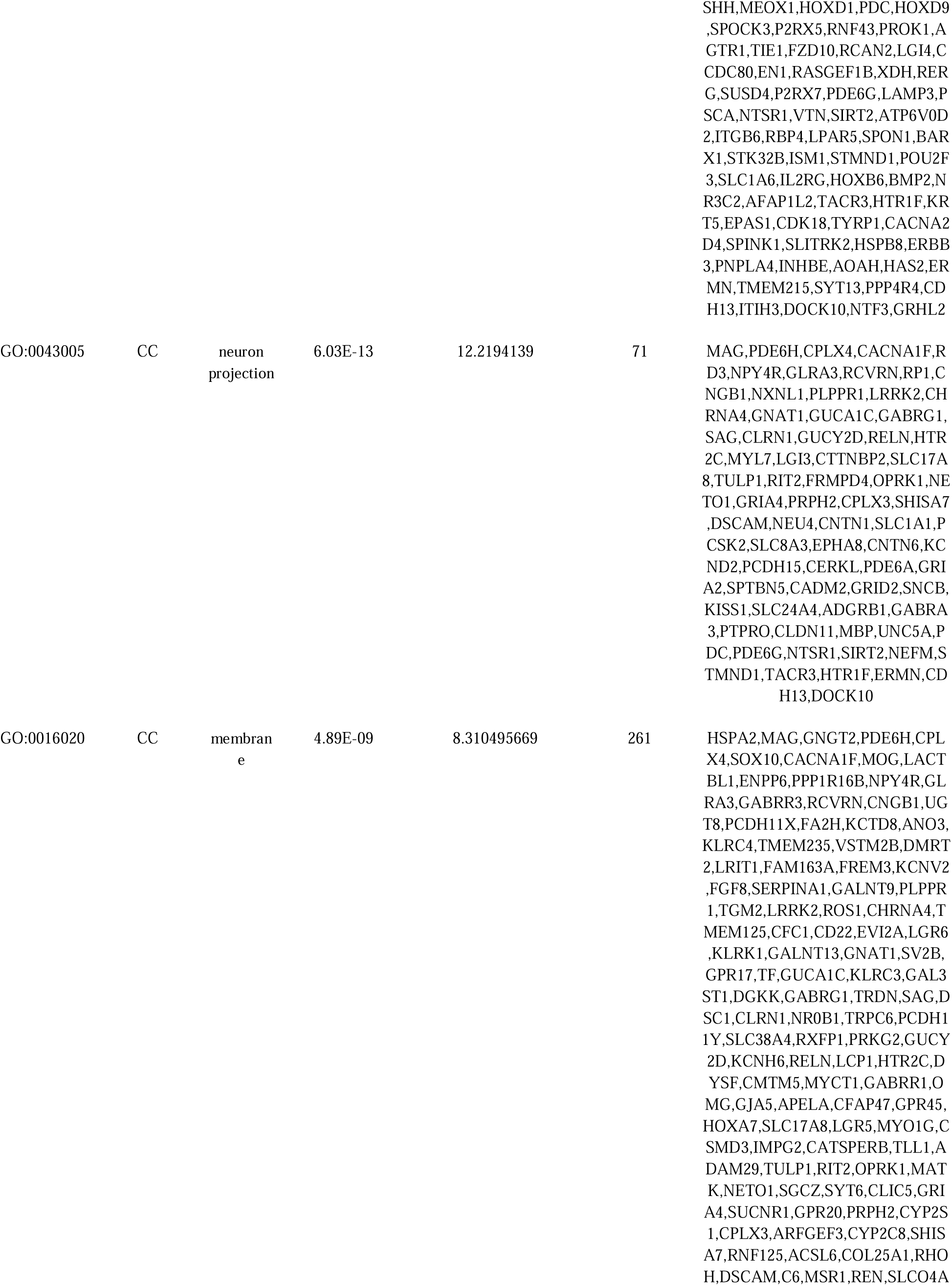
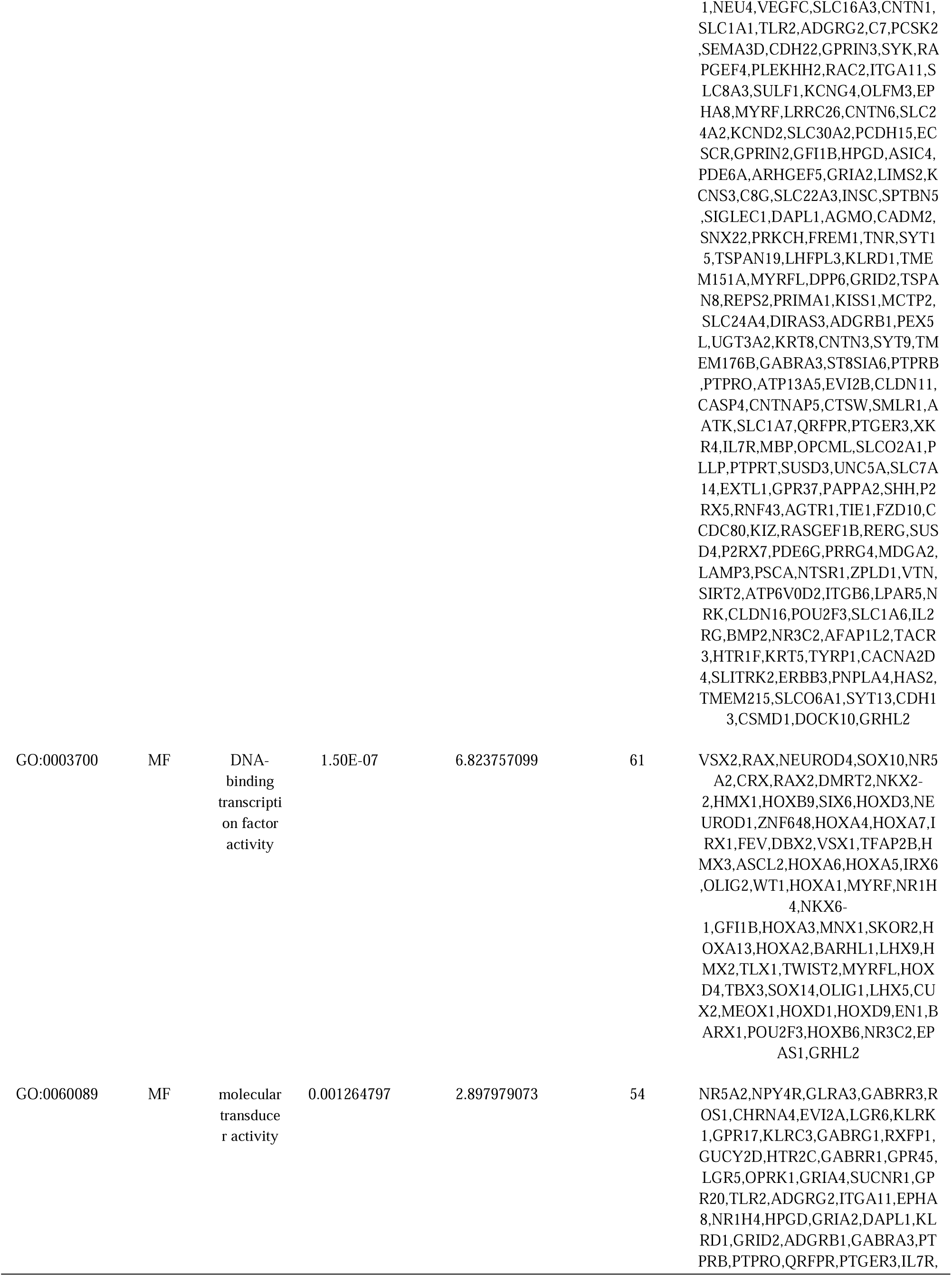
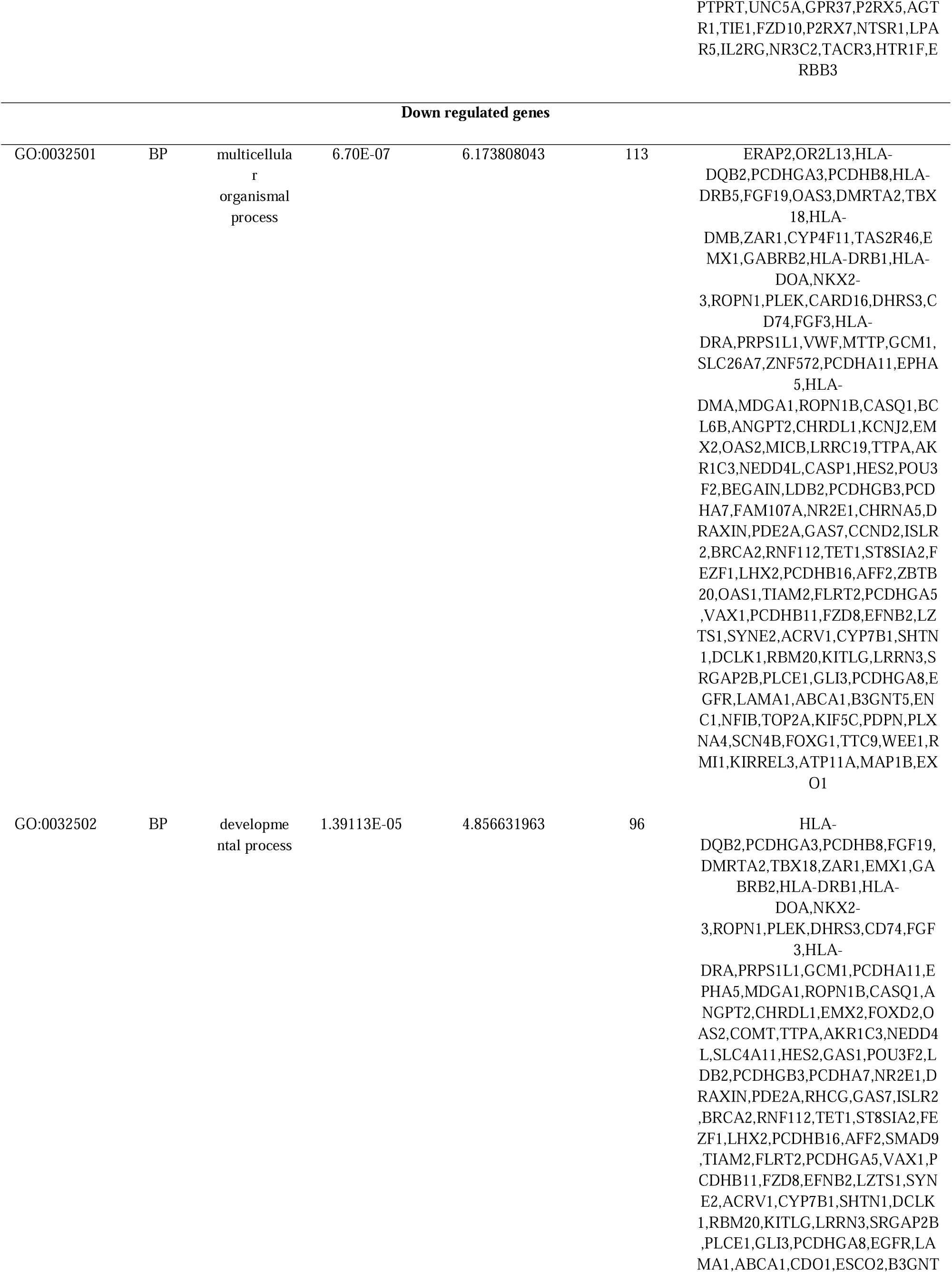
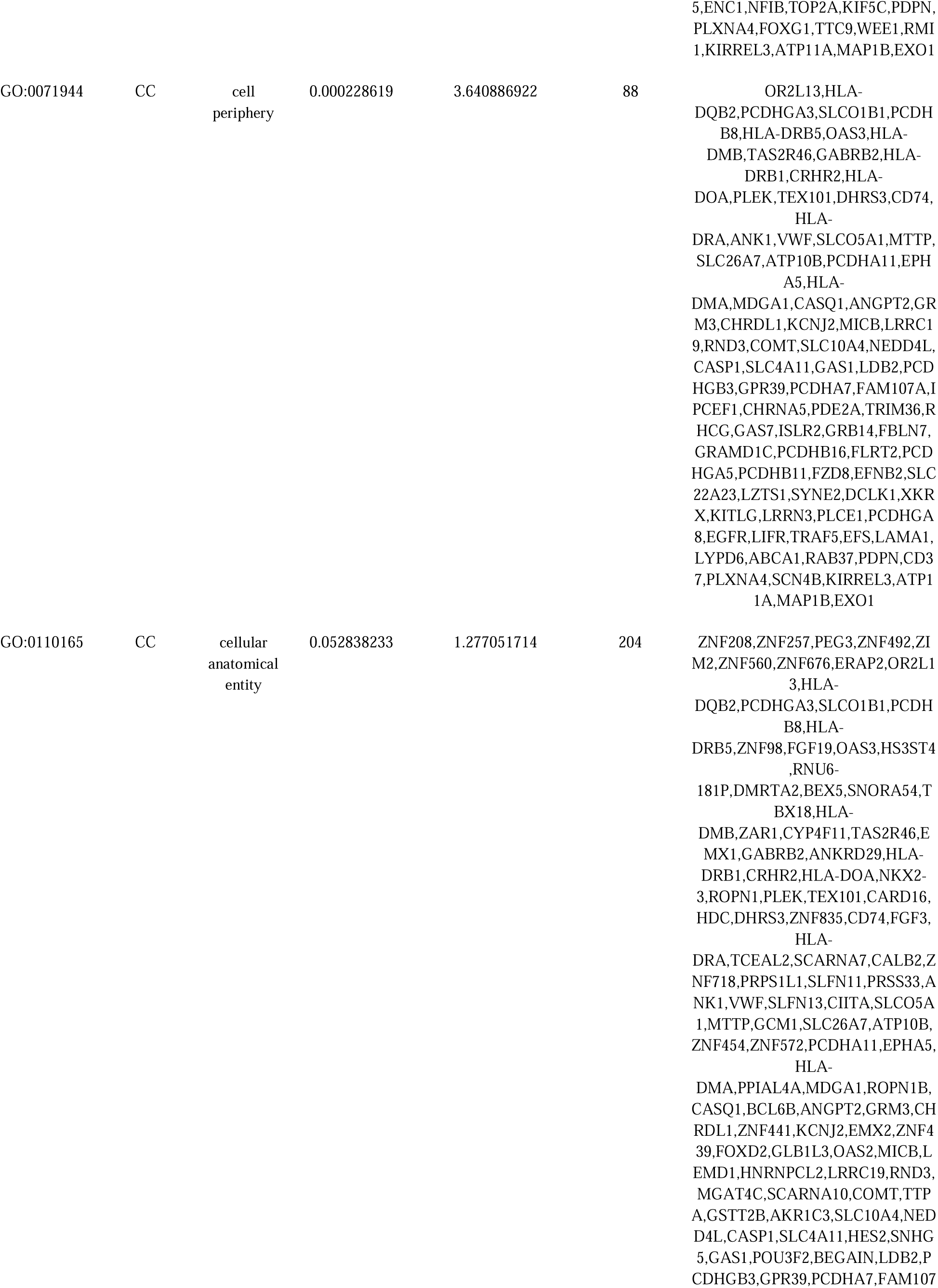
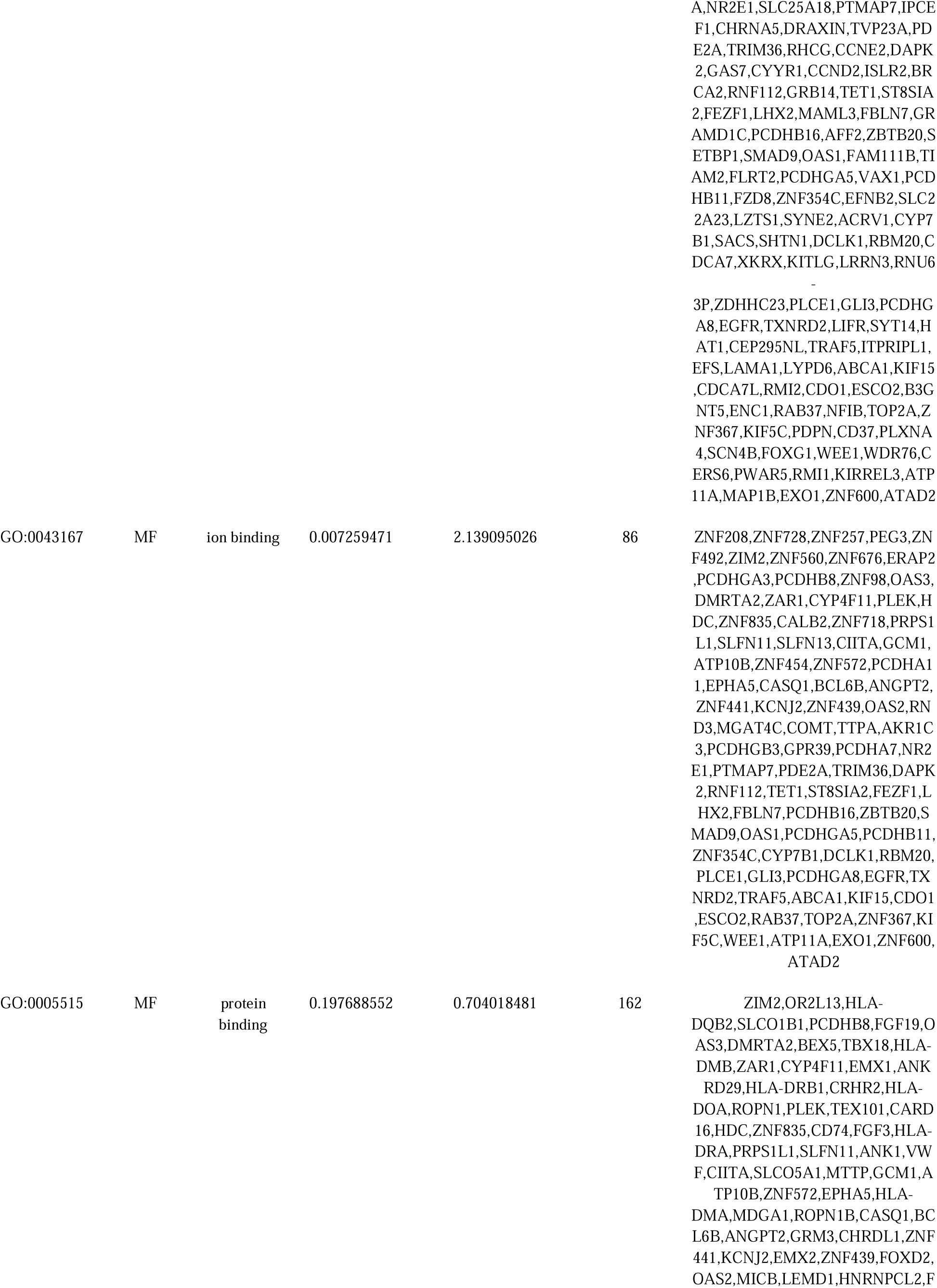
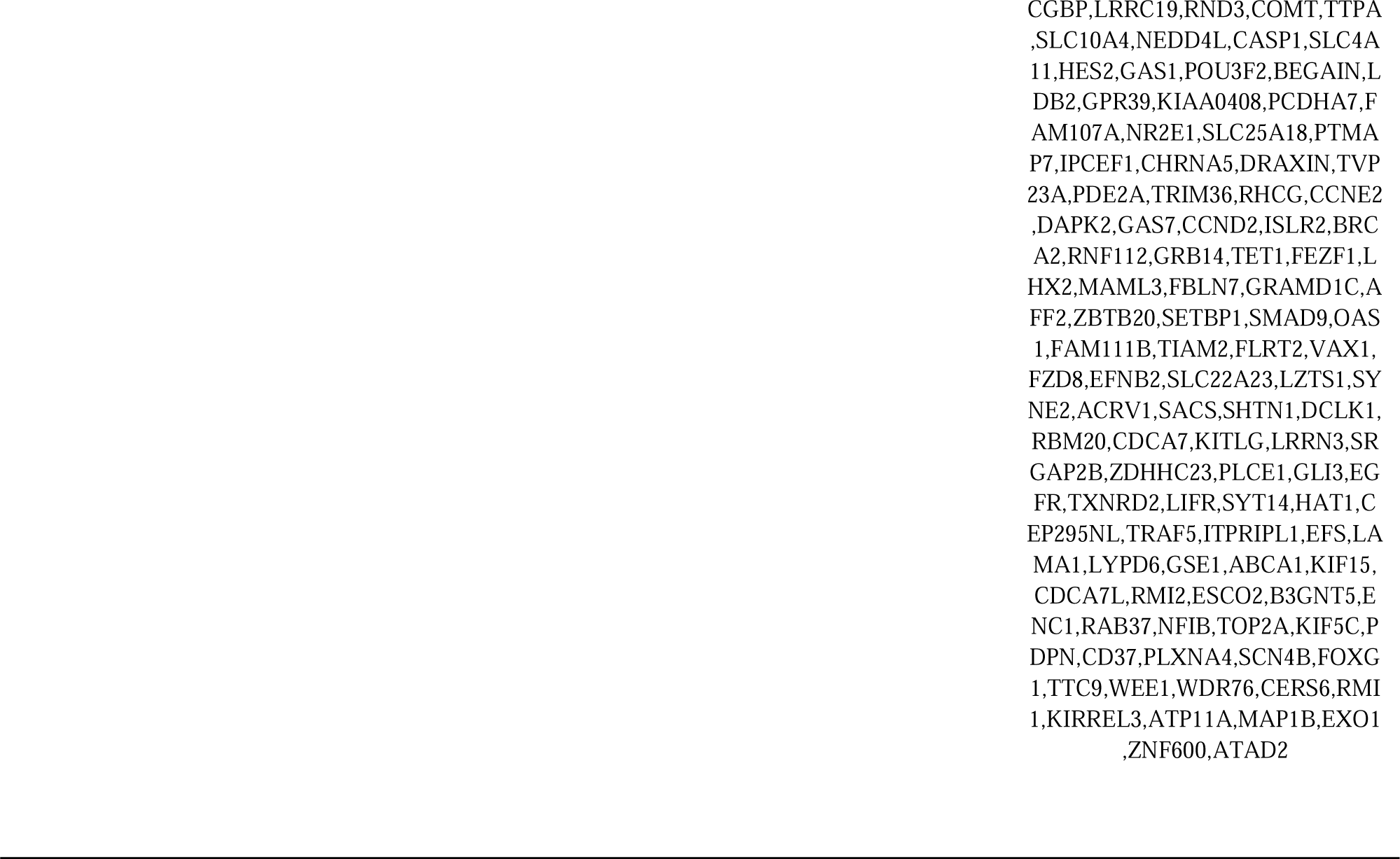
The enriched GO terms of the up and down regulated differentially expressed genes

**Table 3.**
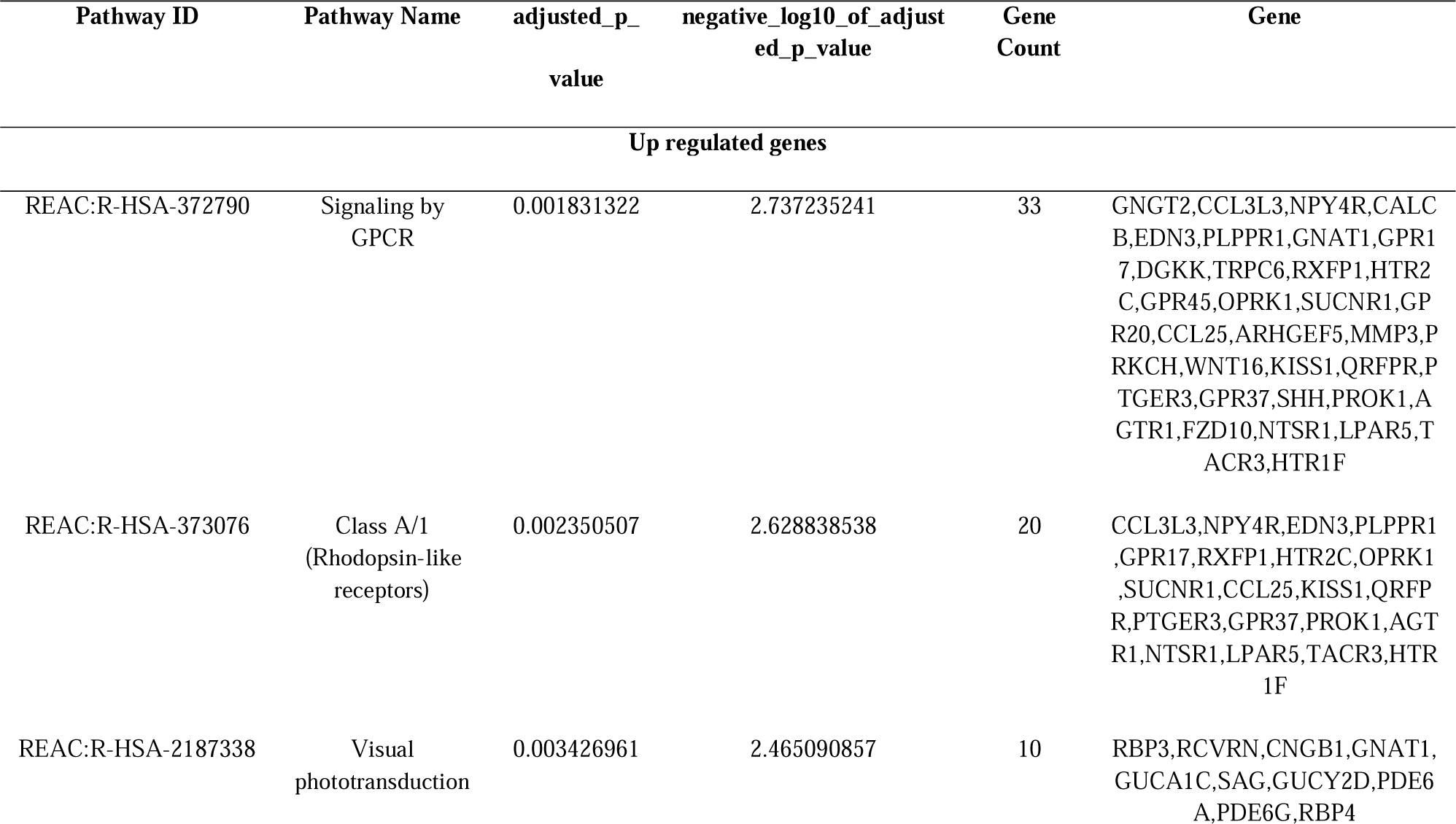
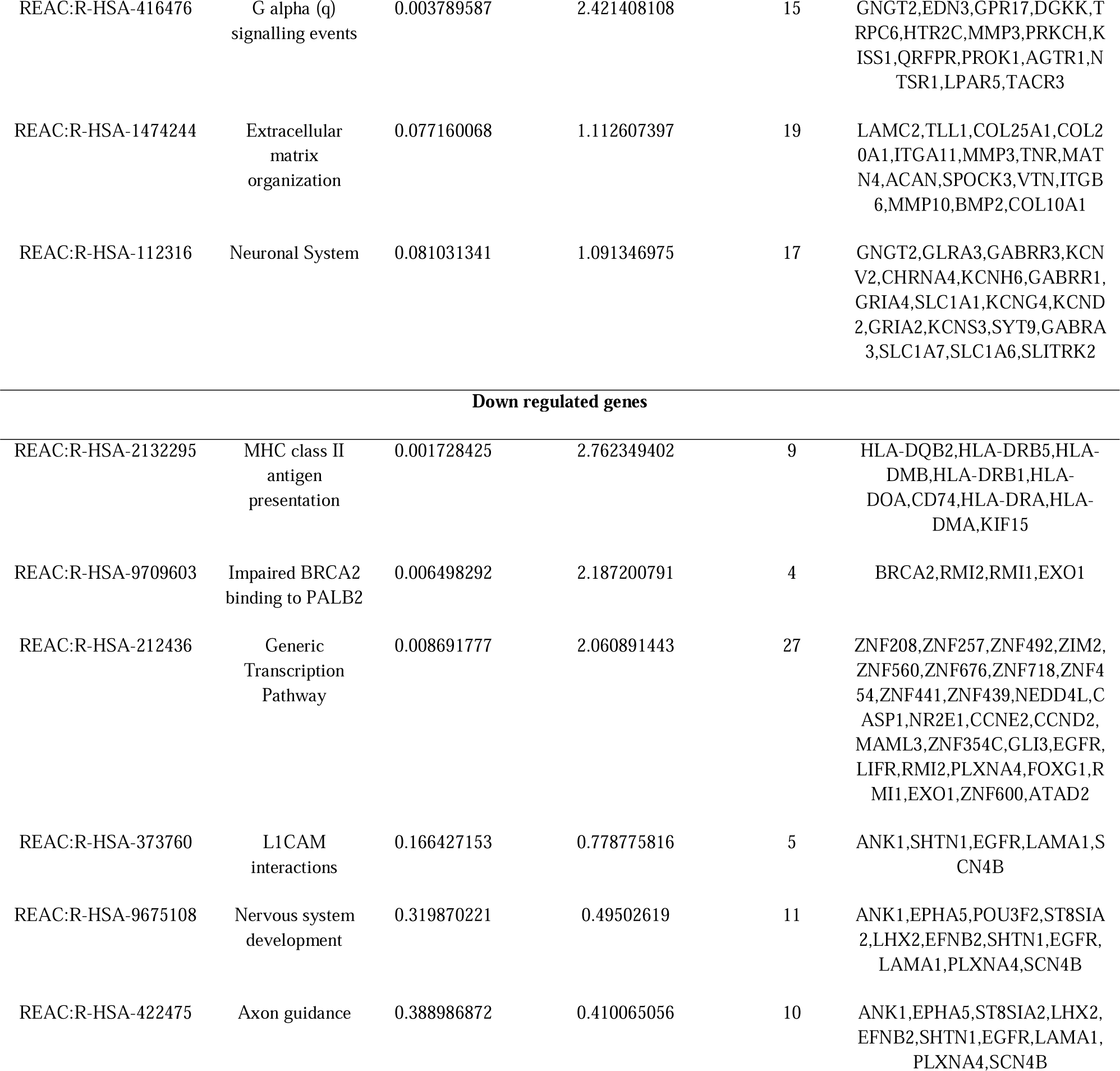
The enriched pathway terms of the up and down regulated differentially expressed genes

### Construction of the PPI network and module analysis

The IID database was used to identify the PPI pairs. As revealed in Fig 3 , 7583 nodes (genes) and 15228 edges (interactions) were established in the constructed PPI network. According to the node degree, betweenness, stress and closeness value, the hub genes were determined. The results shown that LRRK2, MTUS2, HOXA1, IL7R, ERBB3, EGFR, TEX101, WDR76, NEDD4L and COMT were the hub gene with the highest node degree, betweenness, stress and closeness (Table 4). Furthermore, the 2 significant modules were extracted from the PPI network. Module 1 contained 13 gene nodes and 24 edges (Fig. 4A). Module 2 contained 19 genes nodes and 38 edges (Fig. 4B). The functional and pathway analyses of genes involved in two significant modules were analyzed using g:Profiler. Results showed that genes in module 1 was mainly enriched in multicellular organismal process. Module 2 was mainly enriched in cellular anatomical entity, multicellular organismal process and protein binding.

**Fig. 3.**
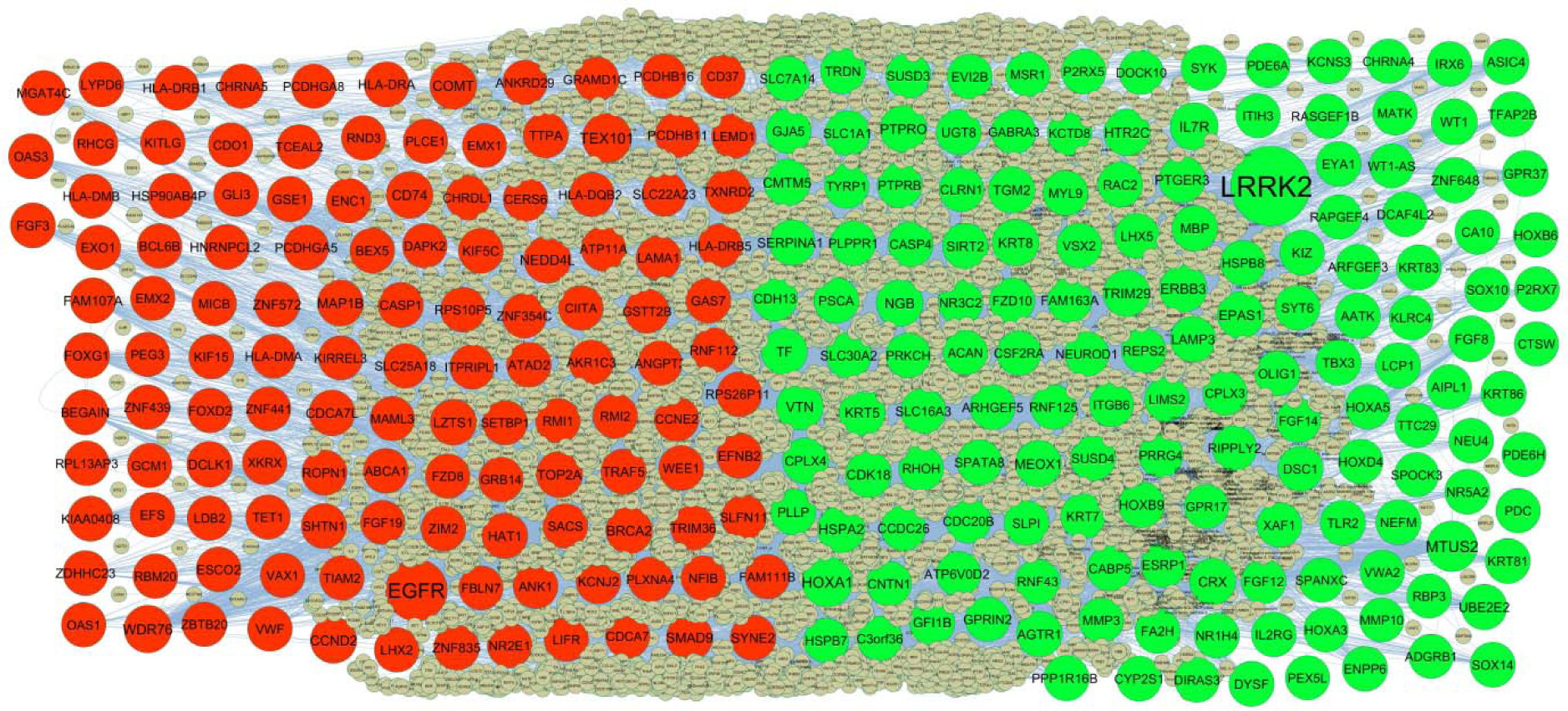
PPI network of DEGs. Up regulated genes are marked in parrot green; down regulated genes are marked in red.

**Fig. 4.**
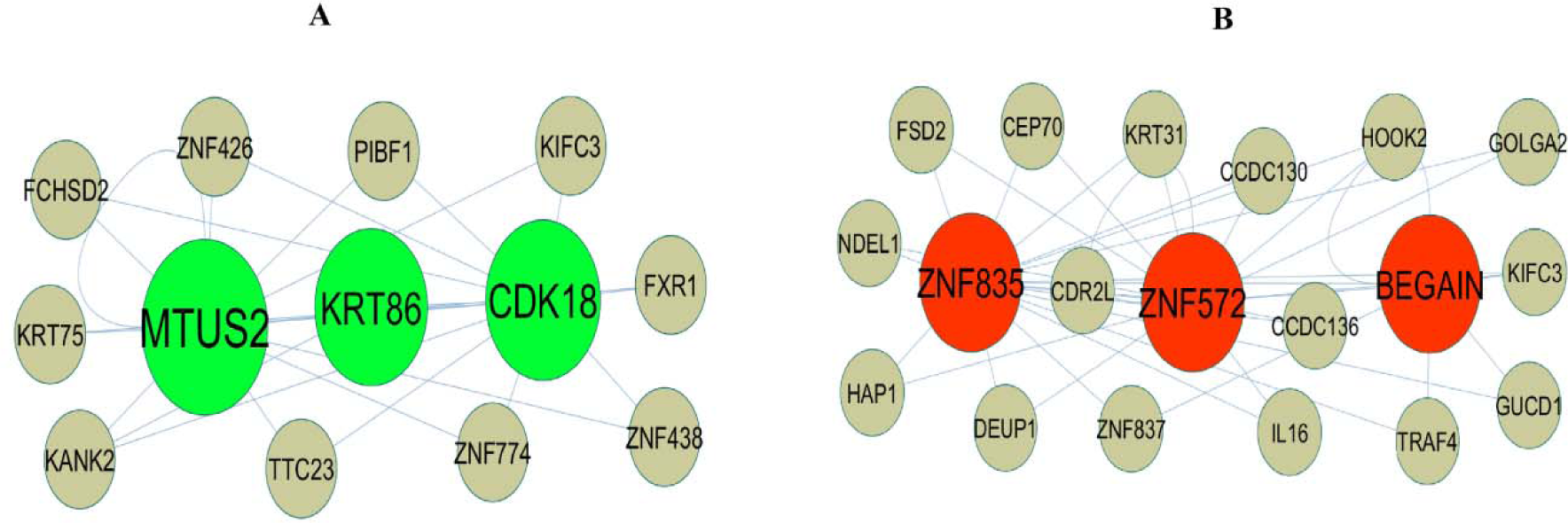
Modules selected from the PPI network. (A) The most significant module was obtained from PPI network with 13 nodes and 24 edges for up regulated genes (B) The most significant module was obtained from PPI network with 19 nodes and 38 edges for down regulated genes. Up regulated genes are marked in parrot green; down regulated genes are marked in red.

**Table 4.**
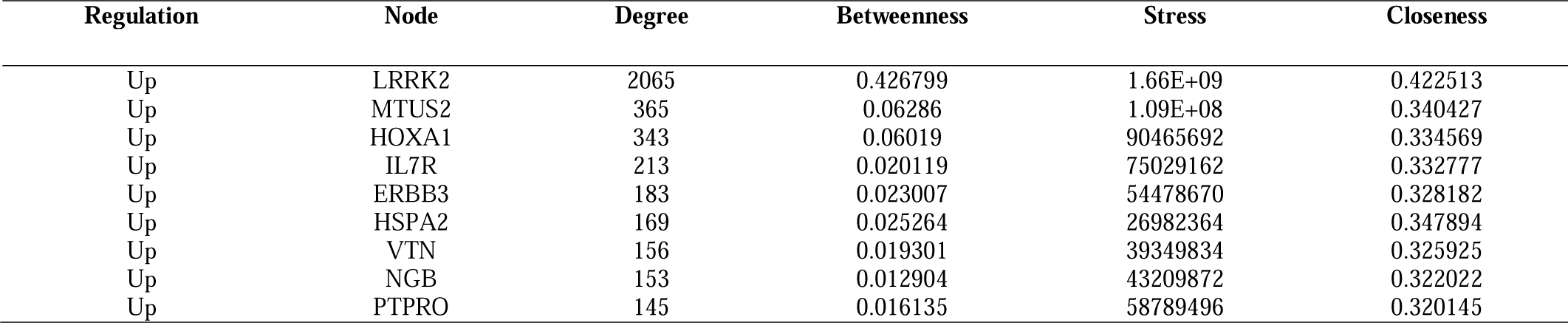
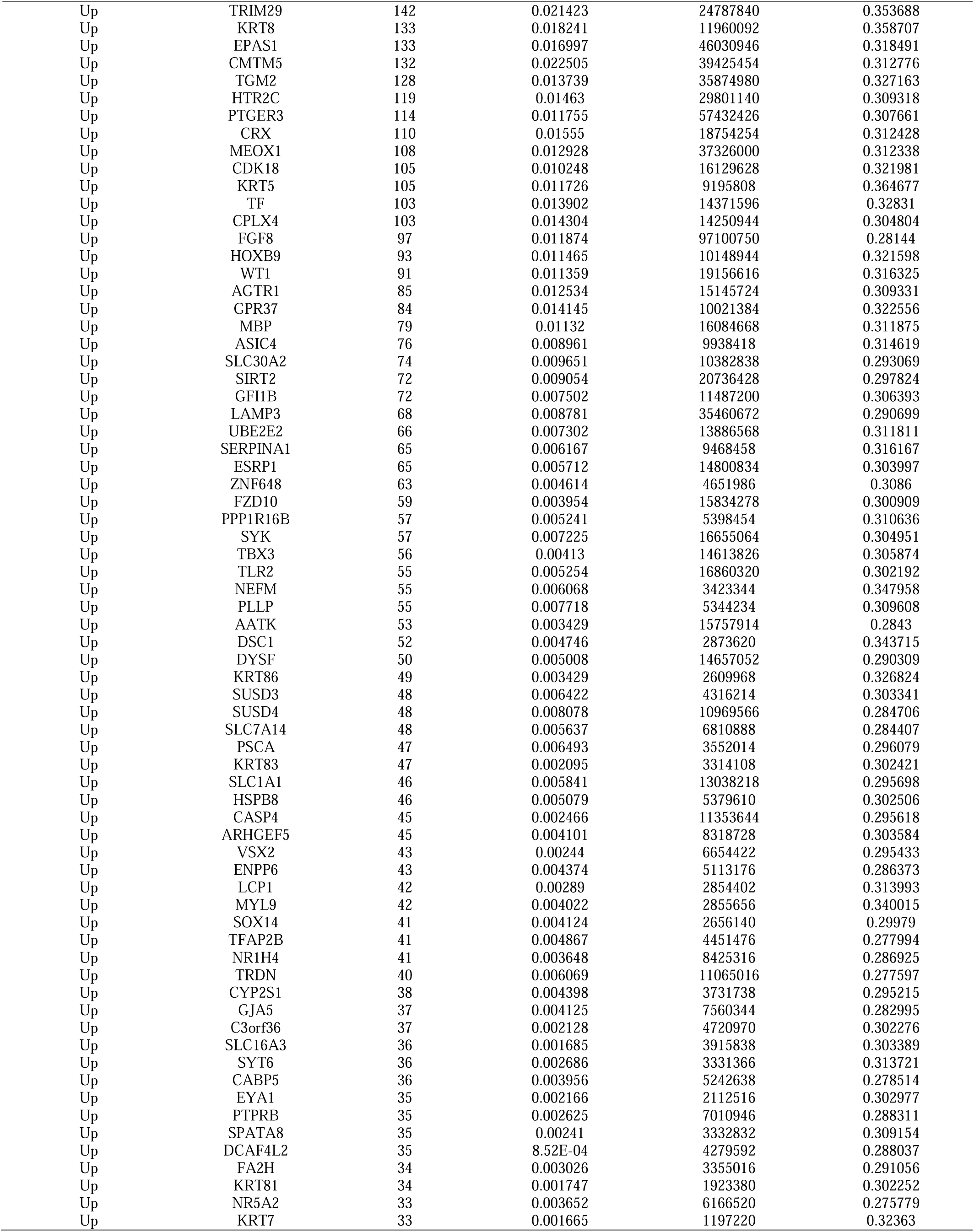
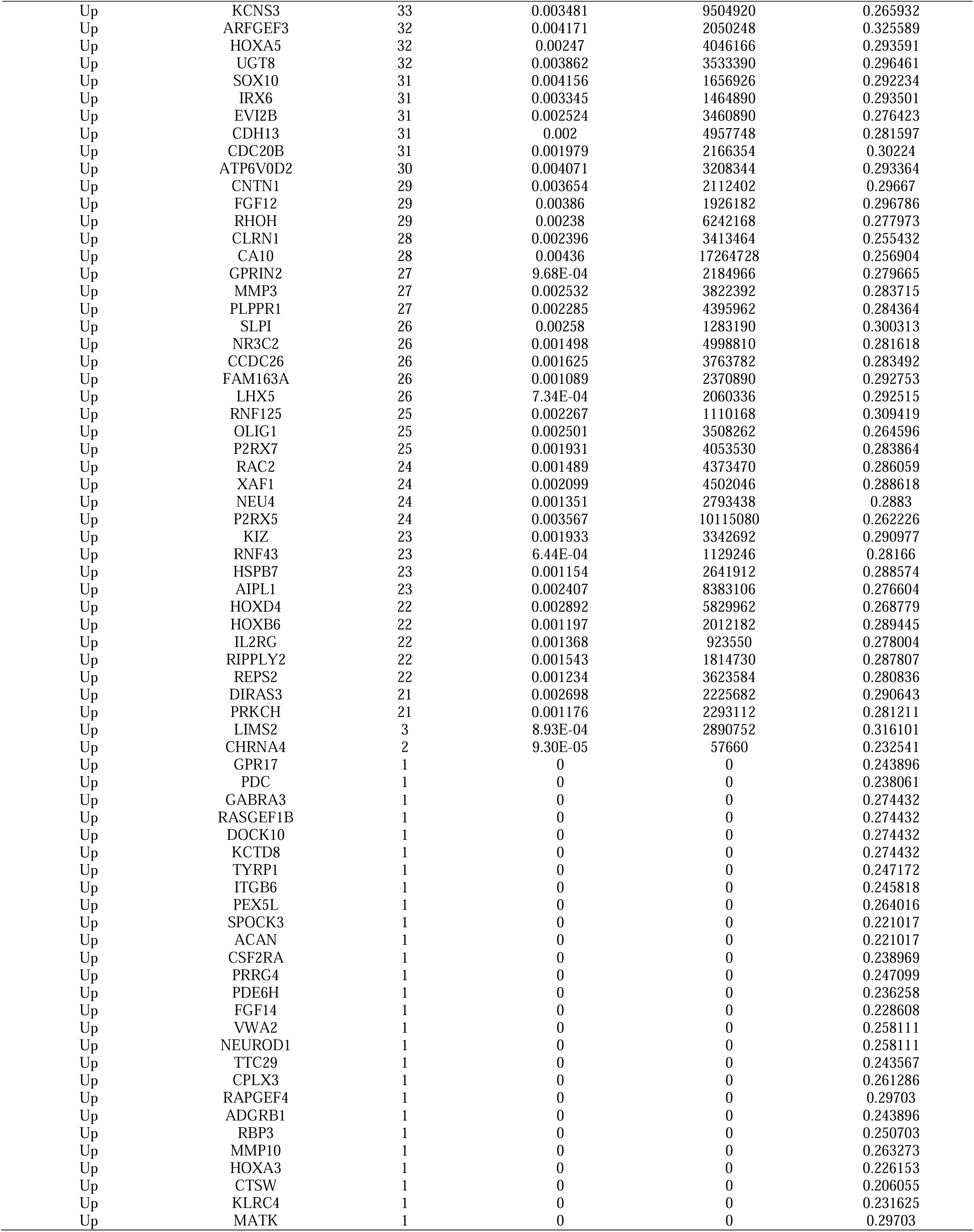
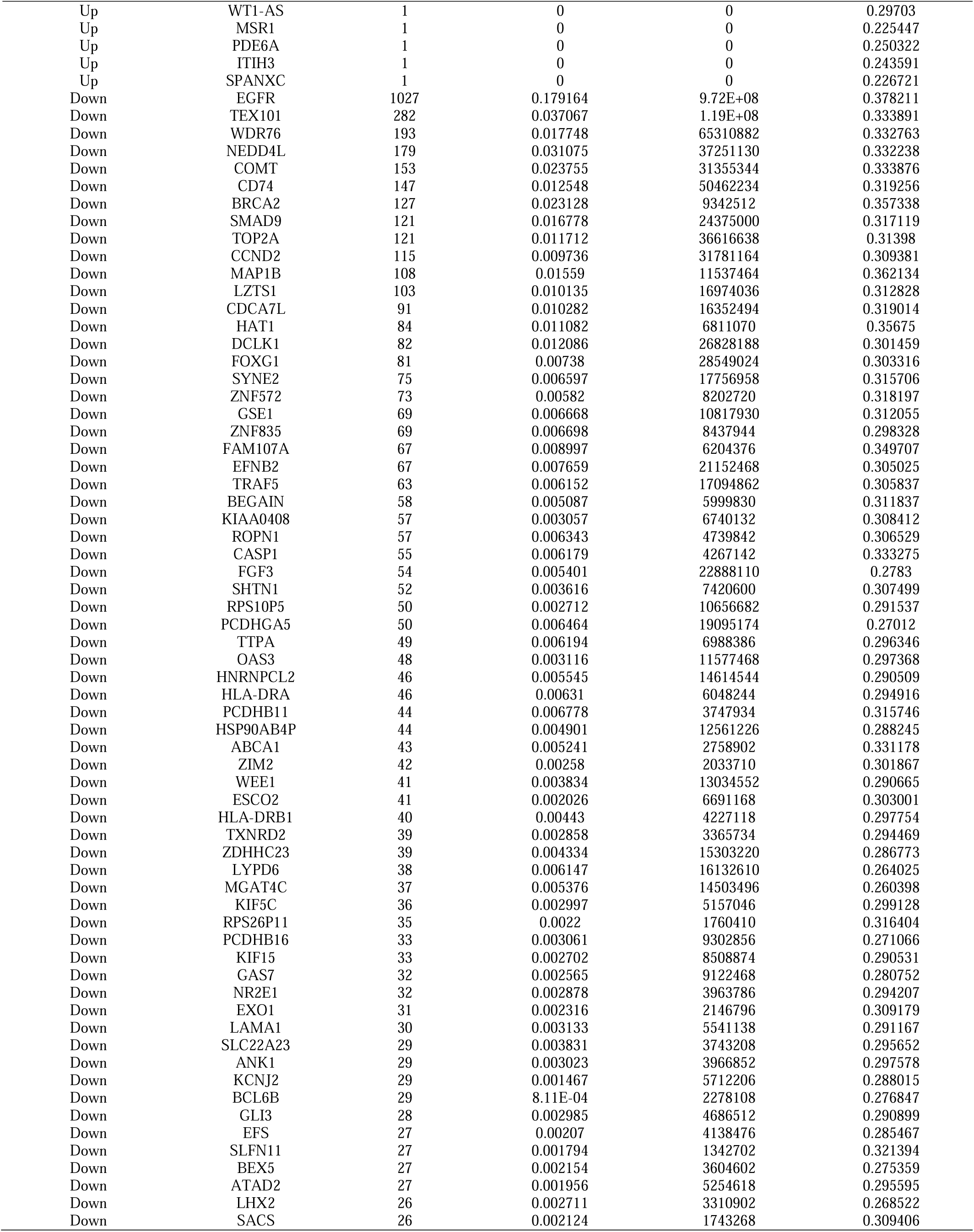
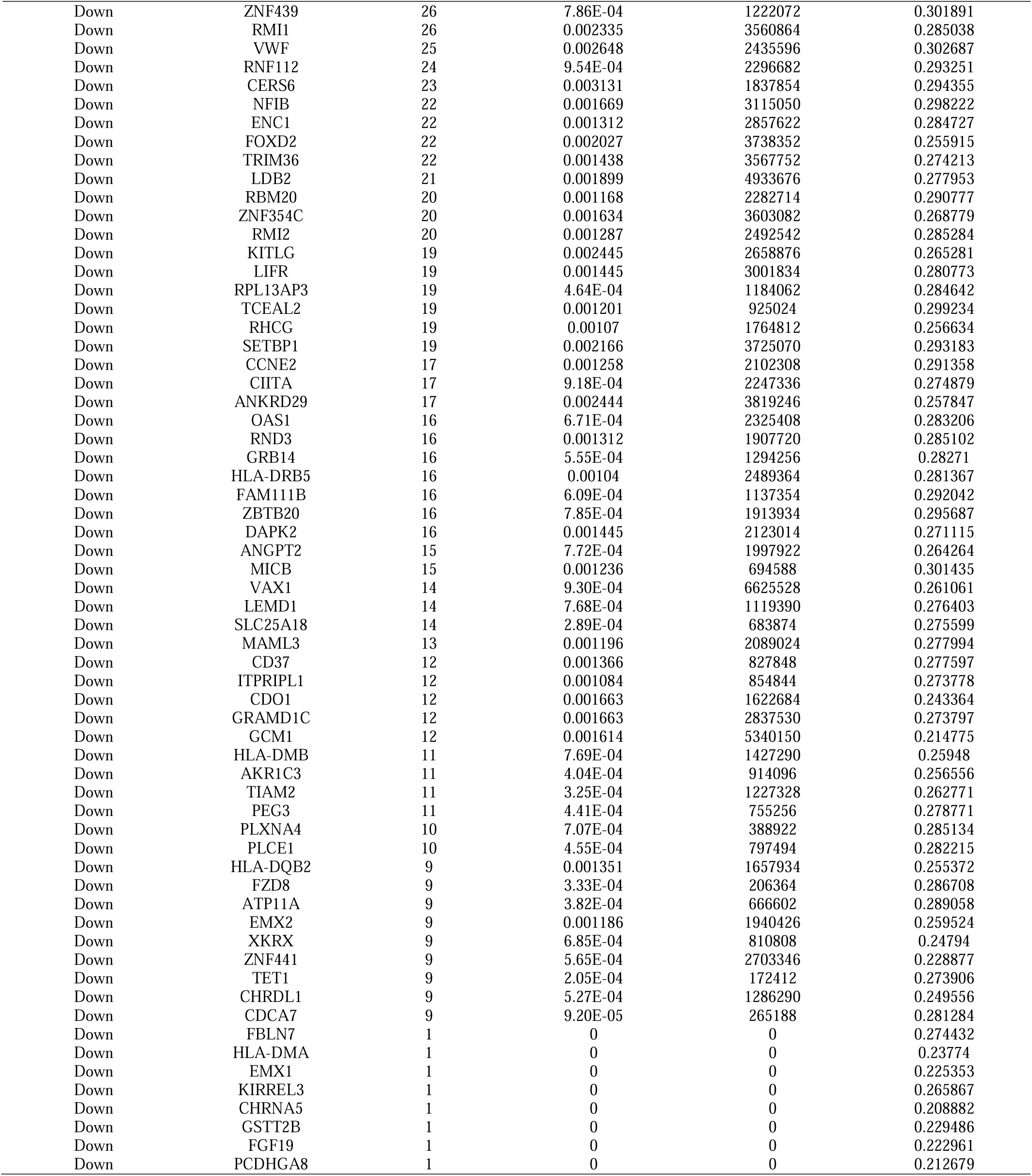
Topology table for up and down regulated genes

### Construction of the miRNA-hub gene regulatory network

MiRNA-hub gene regulatory network was generated using the miRNet web tool. The network contained 2208 nodes (miRNA: 1938; Hub Gene: 270) and 9807 edges (Fig. 5). EPAS1 was regulated by 89 miRNAs (ex: hsa-mir-1292-5p); KRT8 was regulated by 65 miRNAs (ex: hsa-mir-432-3p); ERBB3 was regulated by 53 miRNAs (ex: hsa-mir-6801-3p); HOXA1 was regulated by 45 miRNAs (ex: hsa-mir-10a); HSPA2 was regulated by 45 miRNAs (ex: hsa-mir-7977); CCND2 was regulated by 365 miRNAs (ex: hsa-mir-4521); MAP1B was regulated by 249 miRNAs (ex: hsa-mir-363-3p); TOP2A was regulated by 104 miRNAs (ex: hsa-mir-4487); EGFR was regulated by 83 miRNAs (ex: hsa-mir-21-5p); NEDD4L was regulated by 76 miRNAs (ex: hsa-mir-6807-5p) (Table 5).

**Fig. 5.**
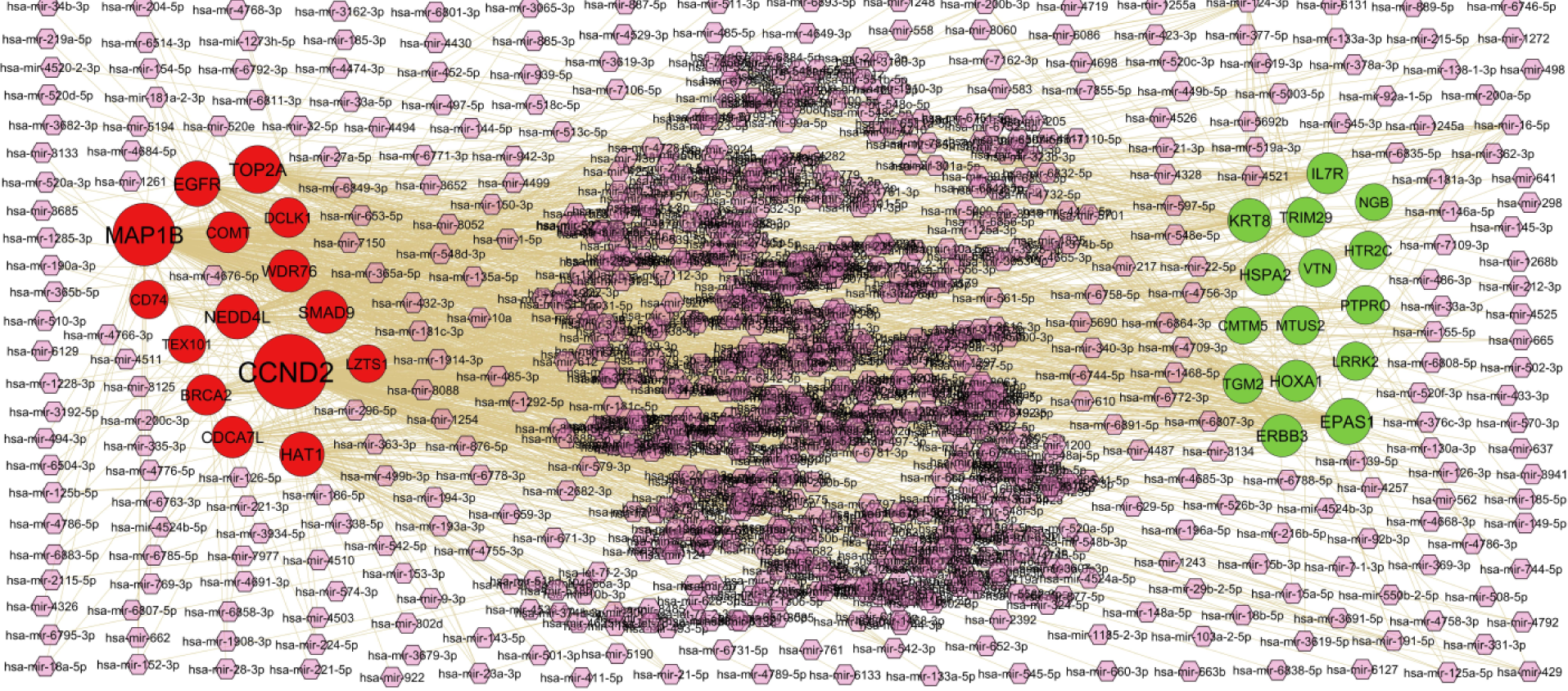
Hub gene - miRNA regulatory network. The purple color diamond nodes represent the key miRNAs; up regulated genes are marked in green; down regulated genes are marked in red.

**Table 5.** MiRNA - hub gene and TF – hub gene topology table

| Regulation | Hub Genes | Degree | MicroRNA | Regulation | Hub Genes | Degree | TF |
| --- | --- | --- | --- | --- | --- | --- | --- |
| Up | EPAS1 | 89 | hsa-mir-1292-5p | Up | EPAS1 | 49 | ESRRB |
| Up | KRT8 | 65 | hsa-mir-432-3p | Up | TGM2 | 43 | SMAD4 |
| Up | ERBB3 | 53 | hsa-mir-6801-3p | Up | HOXA1 | 37 | EED |
| Up | HOXA1 | 45 | hsa-mir-10a | Up | KRT8 | 31 | ATF3 |
| Up | HSPA2 | 45 | hsa-mir-7977 | Up | MTUS2 | 30 | TBX5 |
| Up | IL7R | 40 | hsa-mir-21-3p | Up | HSPA2 | 27 | JARID2 |
| Up | TGM2 | 27 | hsa-mir-452-5p | Up | ERBB3 | 25 | CLOCK |
| Up | TRIM29 | 27 | hsa-mir-761 | Up | PTPRO | 25 | SOX11 |
| Up | PTPRO | 18 | hsa-mir-429 | Up | VTN | 20 | PBX1 |
| Up | MTUS2 | 18 | hsa-mir-133a-3p | Up | IL7R | 16 | TCF4 |
| Up | LRRK2 | 14 | hsa-mir-149-5p | Up | LRRK2 | 16 | PPARD |
| Up | HTR2C | 12 | hsa-mir-22-5p | Up | HTR2C | 15 | TFCP2L1 |
| Up | VTN | 7 | hsa-mir-16-5p | Up | TRIM29 | 15 | YAP1 |
| Up | CMTM5 | 2 | hsa-mir-520f-3p | Up | CMTM5 | 15 | SCLY |
| Up | NGB | 1 | hsa-mir-146a-5p | Up | NGB | 5 | PAX6 |
| Down | CCND2 | 365 | hsa-mir-4521 | Down | NEDD4L | 50 | SREBF1 |
| Down | MAP1B | 249 | hsa-mir-363-3p | Down | CCND2 | 42 | SMARCA4 |
| Down | TOP2A | 104 | hsa-mir-4487 | Down | EGFR | 39 | ARNT |
| Down | EGFR | 83 | hsa-mir-21-5p | Down | MAP1B | 35 | EP300 |
| Down | NEDD4L | 76 | hsa-mir-6807-5p | Down | DCLK1 | 31 | CREB1 |
| Down | HAT1 | 73 | hsa-mir-1185-2-3p | Down | COMT | 29 | SETDB1 |
| Down | SMAD9 | 58 | hsa-mir-1273h-5p | Down | CDCA7L | 27 | DCP1A |
| Down | WDR76 | 52 | hsa-mir-6858-3p | Down | BRCA2 | 26 | MYCN |
| Down | CDCA7L | 45 | hsa-mir-124-3p | Down | HAT1 | 25 | CUX1 |
| Down | COMT | 40 | hsa-mir-1908-3p | Down | TOP2A | 25 | FOXM1 |
| Down | BRCA2 | 37 | hsa-mir-1245a | Down | LZTS1 | 20 | NR3C1 |
| Down | DCLK1 | 28 | hsa-mir-125b-5p | Down | WDR76 | 14 | BACH1 |
| Down | CD74 | 15 | hsa-mir-92a-1-5p | Down | CD74 | 12 | IRF8 |
| Down | LZTS1 | 9 | hsa-mir-204-5p | Down | TEX101 | 9 | RCOR3 |
| Down | TEX101 | 7 | hsa-mir-27a-5p | Down | SMAD9 | 9 | TET1 |

### Construction of the TF-hub gene regulatory network

TF-hub gene regulatory network was generated using the NetworkAnalyst web tool. The network contained 465 nodes (TF: 187; Hub Gene: 278) and 5830 edges (Fig. 6). EPAS1 was regulated by 49 TFs (ex: ESRRB); TGM2 was regulated by 43 TFs (ex: SMAD4); HOXA1 was regulated by 37 TFs (ex: EED); KRT8 was regulated by 31 TFs (ex: ATF3); MTUS2 was regulated by 30 TFs (ex: TBX5); NEDD4L was regulated by 50 TFs (ex: SREBF1); CCND2 was regulated by 42 TFs (ex: SMARCA4); EGFR was regulated by 39 TFs (ex: ARNT); MAP1B was regulated by 35 TFs (ex: EP300); DCLK1 was regulated by 31 TFs (ex: CREB1) (Table 5).

**Fig. 6.**
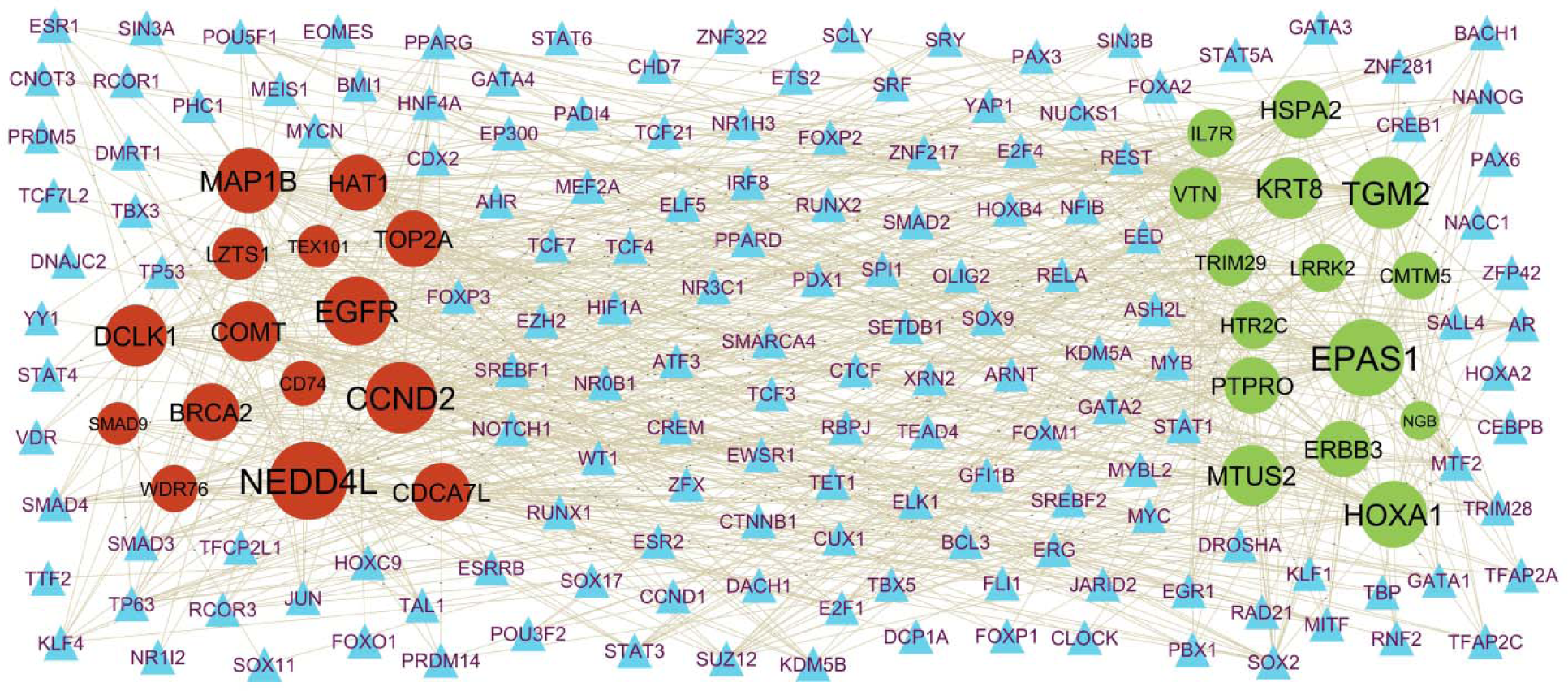
Hub gene - TF regulatory network. The blue color triangle nodes represent the key TFs; up regulated genes are marked in dark green; down regulated genes are marked in dark red.

### Receiver operating characteristic curve (ROC) analysis

To validate the diagnostic effectiveness of LRRK2, MTUS2, HOXA1, IL7R, ERBB3, EGFR, TEX101, WDR76, NEDD4L and COMT for HD by ROC analysis. The larger the AUC, the more the capability of the molecular biomarker to diagnose HD with excellent specificity and sensitivity. As shown in Fig 7, the AUC value of LRRK2 was 0.910; AUC value of MTUS2 was 0.912; AUC value of HOXA1 was 0.891; AUC value of IL7R was 0.894; AUC value of ERBB3 was 0.896; AUC value of EGFR was 0.916; AUC value of TEX101 was 0.900; AUC value of WDR76 was 0.882; AUC value of NEDD4L was 0.887; AUC value of NEDD4L was 0.907.

**Fig. 7.**
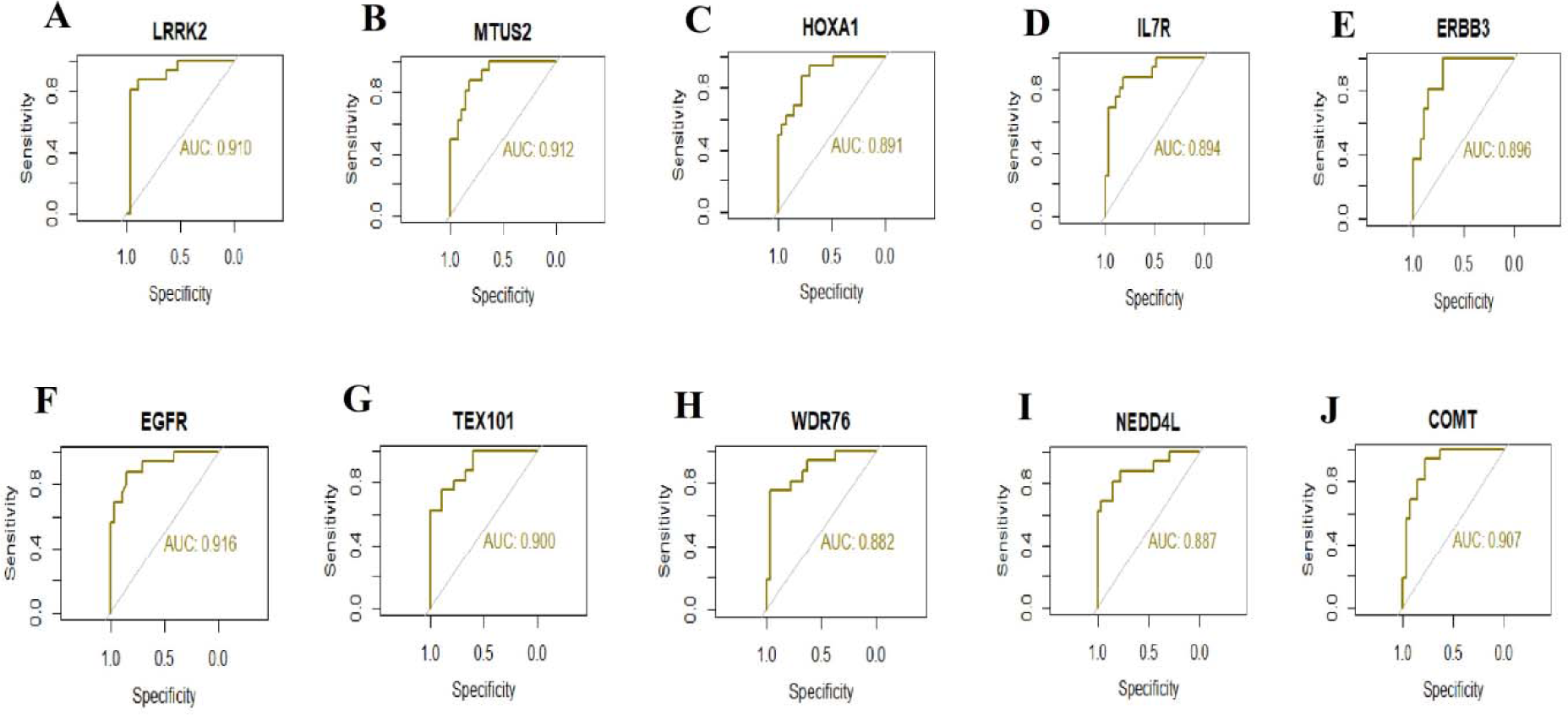
ROC curve analyses of hub genes. A) LRRK2 B) MTUS2 C) HOXA1 D) IL7R E) ERBB3 F) EGFR G) TEX101 H) WDR76 I) NEDD4L J) COMT

## Discussion

HD is the main cause of neurodegenerative disease globally [41]. Due to the extremely complex CNS disorders that occur in patients with HD, once HD has reached the terminal stage, it is often more difficult to treat than other CNS diseases. Although many investigations have studied the molecular pathogenesis of HD, it has not been clarified completely [42]. Therefore, it is necessary to identify potential biomarkers for early diagnosis and targeted therapy of HD. This investigation used NGS data to conduct bioinformatics analysis for identifying target genes and pathways involved in the occurrence and advancement of HD. The results of this investigation suggest that novel biomarkers and signaling pathways may play a key role in the pathogenesis of HD.

We first downloaded HD related NGS dataset, GSE105041 from the GEO database, which were used for analysis to obtain DEGs. A total of 958 DEGs were identified in HD patients compared with the normal control samples, including 479 up regulated and 479 down regulated genes. HSPA2 [43] might be a potential target for Alzheimer’s disease treatment. BCAS1 [44] and MAG (myelin associated glycoprotein) [45] engages in the development of multiple sclerosis. Previous studies have demonstrated that BCAS1 [46] and MAG (myelin associated glycoprotein) [47] are linked with the development mechanisms of schizophrenia. Levels of BCAS1 [48], MAG (myelin associated glycoprotein) [49], ZNF208 [50] and PEG3 [51] are increased in patients with stroke. MAG (myelin associated glycoprotein) [52] participate in pathogenic processes of Parkinson’s disease. Recent evidence indicates that BCAS1 [46], MAG (myelin associated glycoprotein) [49] and CES1P1 [53] are key to progression of inflammation. Previous research suggested some cardiac biomarkers, such as ZNF208 [54] and PEG3 [55] could be valuable in the diagnosis and prognosis of cardiovascular diseases. The above results suggest that significant DEGs might be involved in the progression of HD.

To obtain a deeper understanding into the biological functions of DEGs, GO annotation and REACTOME pathway enrichment analyses were performed. These enrichment analyses were used to explore the molecular mechanisms of enriched genes involved in the occurrence and advancement of HD. Signaling pathways include signaling by GPCR [56] and extracellular matrix organization [57] were responsible for advancement of HD. Research has shown that CBLN2 [58], MOG (myelin oligodendrocyte glycoprotein) [59], ROS1 [60], LGR6 [61], GJA5 [62], SUCNR1 [63], REN (renin) [64], VEGFC (vascular endothelial growth factor C) [65], TLR2 [66], FGF12 [67], WT1 [68], SLC22A3 [69], CSRP3 [70], PAPPA2 [71], ACAN (aggrecan) [72], SHH (sonic hedgehog signaling molecule) [73], PDC (phosducin) [74], AGTR1 [75], XDH (xanthine dehydrogenase) [76], P2RX7 [77], NTSR1 [78], SIRT2 [79], RBP4 [80], BMP2 [81], ERBB3 [82], HAS2 [83], CDH13 [84], CSMD1 [85], NR3C2 [86], EPAS1 [87], TRPC6 [88], MMP3 [89], ERAP2 [90], HLA-DRB1 [91], CD74 [92], VWF (von Willebrand factor) [93], MTTP (microsomal triglyceride transfer protein) [494], ANGPT2 [184], AKR1C3 [495], NEDD4L [496], CASP1 [497], LDB2 [498], CHRNA5 [499], CCND2 [500], BRCA2 [97], ZBTB20 [501], EFNB2 [502], CYP7B1 [503], DCLK1 [504], RBM20 [505], PLCE1 [197], protein) [94], NEDD4L [95], CASP1 [96], BRCA2 [97], EFNB2 [98], PLCE1 [99], EGFR (epidermal growth factor receptor) [100], ABCA1 [101], CRHR2 [102], RND3 [103], COMT (catechol-O-methyltransferase) [104] and SMAD9 [105] plays an important role in the pathogenesis of hypertension. Studies have revealed that NEUROD4 [106], SOX10 [107], MOG (myelin oligodendrocyte glycoprotein) [108], NR5A2 [109], GLRA3 [110], FA2H [111], SERPINA [112], HOXB9 [113], TGM2 [114], LRRK2 [115], ROS1 [116], CD22 [117], LGR6 [118], GDF3 [119], GPR17 [120], TF (transferrin) [566], PCDH11Y [567], RXFP1 [568], RELN (reelin) [569], DYSF (dysferlin) [570], RIT2 [324], OPRK1 [571], SHISA7 [572], MSR1 [573], REN (renin) [574], OLIG2 [575], CNTN1 [576], TLR2 [577], WT1 [578], SYK (spleen [121], NEUROD1 [122], RELN (reelin) [123], LCP1 [124], DYSF (dysferlin) [125], LGR5 [126], SUCNR1 [127], ASCL2 [128], HOXA5 [129], MSR1 [130], REN (renin) [131], NEU4 [132], VEGFC (vascular endothelial growth factor C) [133], OLIG2 [134], SLC1A1 [135], TLR2 [136], WT1 [137], SYK (spleen associated tyrosine kinase) [138], BMP8B [139], GDF6 [140], NKX6-1 [141], SLC22A3 [142], CADM2 [143], FREM1 [144], TWIST2 [145], WNT16 [146], KISS1 [147], TMEM176B [148], MYL9 [149], IL7R [150], MBP (myelin basic protein) [151], GPR37 [152], ACAN (aggrecan) [153], SHH (sonic hedgehog signaling molecule) [154], HOXD9 [155], PROK1 [156], AGTR1 [157], TIE1 [158], P2RX7 [159], NTSR1 [160], VTN (vitronectin) [161], SIRT2 [162], RBP4 [163], LPAR5 [164], IL2RG [165], BMP2 [166], EPAS1 [167], HAS2 [168], ERMN (ermin) [169], CSMD1 [170], NKX6-1 [171], POU2F3 [172], NR3C2 [173], TRPC6 [174], CCL25 [175], MMP3 [176], ERAP2 [177], FGF19 [178], HLA-DRB1 [179], NKX2-3 [180], CD74 [92], HLA-DRA [181], VWF (von Willebrand factor) [182], BCL6B [183], ANGPT2 [184], LRRC19 [185], TTPA (alpha tocopherol transfer protein) [186], NEDD4L [187], CASP1 [188], NR2E1 [189], PDE2A [190], CCND2 [191], TET1 [192], AFF2 [193], ZBTB20 [194], EFNB2 [195], DCLK1 [196], PLCE1 [197], EGFR (epidermal growth factor receptor) [198], ABCA1 [199], NFIB (nuclear factor I B) [200], PDPN (podoplanin) [201], PLXNA4 [202], FOXG1 [203], MAP1B [204], SLCO1B1 [205], RND3 [206], COMT (catechol-O-methyltransferase) [207], GPR39 [208], RHCG (Rh family C glycoprotein) [209], FBLN7 [210], LIFR (LIF receptor subunit alpha) [211], TRAF5 [212], HDC (histidine decarboxylase) [213], SLFN11 [214], CIITA (class II major histocompatibility complex transactivator) [215], DAPK2 [216] and CDO1 [217] plays a key role in inflammation. NEUROD4 [218], NR5A2 [219], GLRA3 [220], DMRT2 [221], NKX2-2 [222], TGM2 [223], LRRK2 [224], CD22 [225], GDF3 [226], TF (transferrin) [227], KLRC3 [228], NEUROD1 [229], KCNH6 [230], LGR5 [231], SUCNR1 [63], TFAP2B [232], MSR1 [233], REN (renin) [234], TLR2 [235], PCSK2 [236], SLC24A2 [237], HOXA3 [238], MNX1 [239], SLC22A3 [240], CADM2 [241], TSPAN8 [242], KISS1 [243], IL7R [244], MBP (myelin basic protein) [245], XAF1 [246], SHH (sonic hedgehog signaling molecule) [247], AGTR1 [248], VTN (vitronectin) [249], SIRT2 [250], RBP4 [251], ISM1 [252], BMP2 [253], SPINK1 [254], ERBB3 [255], CDH13 [256], NPY4R [257], NR3C2 [258], TRPC6 [259], CCL25 [260], MMP3 [261], FGF19 [262], HLA-DMB [263], HLA-DRB1 [264], PLEK (pleckstrin) [265], CD74 [266], VWF (von Willebrand factor) [267], MTTP (microsomal triglyceride transfer protein) [268], HLA-DMA [269], CASQ1 [270], ANGPT2 [271], CASP1 [272], NR2E1 [273], CCND2 [274], OAS1 [275], EGFR (epidermal growth factor receptor) [276], LAMA1 [277], ABCA1 [278], SLCO1B1 [205], CRHR2 [279], ANK1 [280], GPR39 [281], GRB14 [282] and TXNRD2 [283] have been known to be involved in diabetes mellitus progression. SOX10 [284], MOG (myelin oligodendrocyte glycoprotein) [285], TGM2 [286], LRRK2 [287], CD22 [288], GPR17 [120], TF (transferrin) [289], RELN (reelin) [290], DYSF (dysferlin) [291], HOXA5 [292], REN (renin) [293], CNTN1 [294], SLC1A1 [295], TLR2 [296], WT1 [297], MOBP (myelin associated oligodendrocyte basic protein) [298], MYRF (myelin regulatory factor) [299], IL7R [300], MBP (myelin basic protein) [151], SHH (sonic hedgehog signaling molecule) [301], NTSR1 [302], BMP2 [303], ERMN (ermin) [304], MMP3 [305], HLA-DRB5 [306], HLA-DMB [307], HLA-DRB1 [308], HLA-DRA [309], CASP1 [310], OAS1 [311], DCLK1 [312], EGFR (epidermal growth factor receptor) [313], COMT (catechol-O-methyltransferase) [314] and CIITA (class II major histocompatibility complex transactivator) [315] have a significant prognostic potential in multiple sclerosis. SOX10 [316], MOG (myelin oligodendrocyte glycoprotein) [317], CNGB1 [318], TGM2 [319], LRRK2 [320], CHRNA4 [321], RELN (reelin) [322], HTR2C [323], RIT2 [324], ACSL6 [325], C6 [326], REN (renin) [327], OLIG2 [328], TLR2 [329], SEMA3D [330], PCDH15 [331], FGF14 [332], TSPAN8 [333], ACAN (aggrecan) [334], SHH (sonic hedgehog signaling molecule) [335], P2RX7 [336], NTSR1 [337], RBP4 [338], TACR3 [339], ERBB3 [340], ERMN (ermin) [341], CDH13 [342], CSMD1 [343], NTF3 [344], FRMPD4 [345], GRIA4 [346], GRIA2 [347], SNCB (synuclein beta) [348], NR3C2 [349], MMP3 [350], GABRB2 [351], HLA-DRB1 [352], VWF (von Willebrand factor) [353], MTTP (microsomal triglyceride transfer protein) [354], MDGA1 [355], CASP1 [356], POU3F2 [357], LDB2 [358], NR2E1 [359], CHRNA5 [360], GAS7 [361], BRCA2 [362], ST8SIA2 [363], ZBTB20 [364], EFNB2 [365], DCLK1 [366], EGFR (epidermal growth factor receptor) [367], ABCA1 [368], GRM3 [369], COMT (catechol-O-methyltransferase) [370], LDB2 [371], EFNB2 [372] and DCLK1 [373] are molecular markers for the diagnosis and prognosis of schizophrenia. MOG (myelin oligodendrocyte glycoprotein) [374], SERPINA1 [375], LRRK2 [376], CHRNA4 [377], TF (transferrin) [121], RELN (reelin) [378], HTR2C [379], OPRK1 [380], DSCAM (DS cell adhesion molecule) [381], REN (renin) [382], OLIG2 [383], CNTN1 [384], TLR2 [385], WT1 [386], SHH (sonic hedgehog signaling molecule) [387], AGTR1 [388], SIRT2 [389], RBP4 [390], ERBB3 [391], CDH13 [392], NTF3 [344], GRIA4 [393], GRIA2 [394], NR3C2 [395], TRPC6 [396], FGF19 [397], EMX1 [398], VWF (von Willebrand factor) [399], MDGA1 [400], NEDD4L [401], CASP1 [402], TET1 [403], ZBTB20 [404], VAX1 [405], EGFR (epidermal growth factor receptor) [406], ABCA1 [407], PDPN (podoplanin) [408], SCN4B [409], MAP1B [410], CRHR2 [279], GRM3 [411], COMT (catechol-O-methyltransferase) [412], GPR39 [413], CIITA (class II major histocompatibility complex transactivator) [215], ZNF354C [414], KIF15 [415] and SLC1A1 [416] were found to be a key biomarkers to promote its occurrence and progression of depression. FA2H [417], TGM2 [418], LRRK2 [419], GPR17 [420], RELN (reelin) [421], RIT2 [324], REN (renin) [422], TLR2 [423], SYK (spleen associated tyrosine kinase) [138], GRID2 [424], SHH (sonic hedgehog signaling molecule) [425], XDH (xanthine dehydrogenase) [426], VTN (vitronectin) [427], SIRT2 [428], OAS3 [429], TTPA (alpha tocopherol transfer protein) [430], EGFR (epidermal growth factor receptor) [431], ABCA1 [432], WEE1 [433], MAP1B [434], ANK1 [435], GRM3 [436], COMT (catechol-O-methyltransferase) [437], GPR39 [438], HDC (histidine decarboxylase) [439] and TXNRD2 [440] have been proved to participate in neurodegenerative diseases. Altered expression of FA2H [441], DSCAM (DS cell adhesion molecule) [442], OLIG2 [328], PCDH15 [443], GRID2 [424], CTTNBP2 [444], AFF2 [445], SYNE2 [446] and EGFR (epidermal growth factor receptor) [447] are associated with autism spectrum disorder. FA2H [111], SERPINA1 [448], HOXB9 [113], TGM2 [449], LRRK2 [450], ROS1 [451], CFC1 [452], GDF3 [453], TF (transferrin) [121], RXFP1 [454], RELN (reelin) [455], HTR2C [456], MYL7 [457], DYSF (dysferlin) [458], GJA5 [459], IRX1 [460], TLL1 [461], SUCNR1 [462], KERA (keratocan) [463], TFAP2B [464], HOXA5 [465], ACSL6 [466], DSCAM (DS cell adhesion molecule) [467], MSR1 [468], REN (renin) [64], VEGFC (vascular endothelial growth factor C) [469], TLR2 [470], PCSK2 [471], FGF12 [472], SLC22A3 [142], HSPB7 [473], CSRP3 [474], KLRD1 [475], PTPRO (protein tyrosine phosphatase receptor type O) [476], IL7R [477], CUX2 [478], ACAN (aggrecan) [479], SHH (sonic hedgehog signaling molecule) [480], MEOX1 [481], AGTR1 [482], VTN (vitronectin) [483], SIRT2 [79], RBP4 [484], IL2RG [485], EPAS1 [486], CDH13 [487], TRPC6 [488], MMP3 [261], PCDHGA3 [489], FGF19 [490], TBX18 [491], HLA-DRB1 [492]. CD74 [493], VWF (von Willebrand factor) [267], MTTP (microsomal triglyceride transfer protein) [494], ANGPT2 [184], AKR1C3 [495], NEDD4L [496], CASP1 [497], LDB2 [498], CHRNA5 [499], CCND2 [500], BRCA2 [97], ZBTB20 [501], EFNB2 [502], CYP7B1 [503], DCLK1 [504], RBM20 [505], PLCE1 [197], EGFR (epidermal growth factor receptor) [276], ABCA1 [506], PDPN (podoplanin) [507], SCN4B [508], SLCO1B1 [509], CRHR2 [510], RND3 [511], COMT (catechol-O-methyltransferase) [512], GPR39 [513], GRB14 [514], HDC (histidine decarboxylase) [515], DAPK2 [516], SMAD9 [517], TXNRD2 [518] and KIF15 [519] are altered expressed in cardiovascular diseases. Previous studies have proven that the FGF8 [520], SERPINA1 [521], LRRK2 [522], CHRNA4 [523], TF (transferrin) [524], TRDN (triadin) [525], NR0B1 [526], RELN (reelin) [527], HTR2C [528], RIT2 [529], REN (renin) [530], TLR2 [531], MOBP (myelin associated oligodendrocyte basic protein) [532], TNR (tenascin R) [533], ADGRB1 [534], GPR37 [535], SHH (sonic hedgehog signaling molecule) [536], XDH (xanthine dehydrogenase) [537], SIRT2 [538], BMP2 [539], HAS2 [540], CSMD1 [541], NTF3 [542], SNCB (synuclein beta) [543], MMP3 [544], HLA-DRB5 [545], HLA-DRB1 [546], HLA-DRA [547], VWF (von Willebrand factor) [548], AKR1C3 [549], CASP1 [550], CHRNA5 [551], TET1 [552], LRRN3 [553], EGFR (epidermal growth factor receptor) [554], PLXNA4 [555], FOXG1 [556], MAP1B [557], ANK1 [558], ATP10B [559], RND3 [560], COMT (catechol-O-methyltransferase) [561], LIFR (LIF receptor subunit alpha) [562] and TRAF5 [212] are involved in Parkinson’s disease. Recent studies have shown that SERPINA1 [563], LRRK2 [522], CHRNA4 [564], CD22 [565], TF (transferrin) ssociated tyrosine kinase) [579], FGF14 [580], INSC (INSC spindle orientation adaptor protein) [581], BARHL1 [582], PRKCH (protein kinase C eta) [583], SLC24A4 [584], TMEM176B [585], MBP (myelin basic protein) [586], ACAN (aggrecan) [587], SHH (sonic hedgehog signaling molecule) [588], SIRT2 [589], RBP4 [590], ISM1 [591], HOXB6 [592], CDH13 [593], NTF3 [594], TRPC6 [595], MMP3 [596], HLA-DRB5 [597], HLA-DRB1 [546], CD74 [598], VWF (von Willebrand factor) [599], MDGA1 [600], ANGPT2 [601], NEDD4L [602], CASP1 [603], CHRNA5 [604], CCND2 [605], BRCA2 [606], TET1 [607], ZBTB20 [608], OAS1 [609], EGFR (epidermal growth factor receptor) [610], ABCA1 [611], PLXNA4 [612], FOXG1 [613], WEE1 [433], MAP1B [614], ANK1 [615], GRM3 [436], COMT (catechol-O-methyltransferase) [616], SLC10A4 [617], GPR39 [618] and LIFR (LIF receptor subunit alpha) [562] could promote Alzheimer’s disease progression. SERPINA1 [619], LRRK2 [620], CHRNA4 [621], TLR2 [622], MOBP (myelin associated oligodendrocyte basic protein) [623], AATK (apoptosis associated tyrosine kinase) [624], MBP (myelin basic protein) [625], SHH (sonic hedgehog signaling molecule) [626], XDH (xanthine dehydrogenase) [627], RBP4 [628], GRIA2 [629], NR3C2 [630], HLA-DRB5 [631], HLA-DRA [631], NEDD4L [632], CASP1 [633], EFNB2 [634], CYP7B1 [635], EGFR (epidermal growth factor receptor) [636], MAP1B [637], GRM3 [638] and AFF2 [639] plays a central role in amyotrophic lateral sclerosis. Aberrant expression of TGM2 [640], NEUROD1 [641], PRKG2 [642], RIT2 [324], TLR2 [643], MYRF (myelin regulatory factor) [642], SIRT2 [644], MMP3 [645], CASP1 [646], POU3F2 [647] and COMT (catechol-O-methyltransferase) [648] have been observed in HD. SERPINA1 [521], LRRK2 [649], TF (transferrin) [650], REN (renin) [651], TLR2 [652], MOBP (myelin associated oligodendrocyte basic protein) [653], SYK (spleen associated tyrosine kinase) [654], MBP (myelin basic protein) [625], SHH (sonic hedgehog signaling molecule) [655], RBP4 [590], SNCB (synuclein beta) [543], MMP3 [656], VWF (von Willebrand factor) [657], EGFR (epidermal growth factor receptor) [658], ABCA1 [659] and GPR39 [618] are associated with the clinical stages of dementia. Altered expression of LRRK2 [660], GPR17 [661], TF (transferrin) [662], NEUROD1 [663], RELN (reelin) [664], LCP1 [665], TFAP2B [666], HOXA5 [667], MSR1 [668], REN (renin) [669], OLIG2 [670], TLR2 [671], WT1 [672], SLC22A3 [673], CADM2 [674], PRKCH (protein kinase C eta) [675], MBP (myelin basic protein) [676], XAF1 [677], GPR37 [678], SHH (sonic hedgehog signaling molecule) [679], VTN (vitronectin) [680], SIRT2 [681], RBP4 [682], BMP2 [683], CDH13 [684], NR3C2 [685], TRPC6 [686], MMP3 [687], FGF19 [688], HLA-DRB1 [689], CD74 [690], VWF (von Willebrand factor) [691], ANGPT2 [692], CHRDL1 [693], AKR1C3 [694], NEDD4L [695], CASP1 [696], PDE2A [697], GAS7 [698], TET1 [699], EFNB2 [700], EGFR (epidermal growth factor receptor) [701], ABCA1 [702], PDPN (podoplanin) [703], SLCO1B1 [704], COMT (catechol-O-methyltransferase) [705] and GPR39 [706] have been reported to be associated with stroke. CHRNA4 [707], RELN (reelin) [708], VEGFC (vascular endothelial growth factor C) [709], FRMPD4 [710], SCN4B [711], SLCO5A1 [712] and GPR39 [713] are involved in the development of epilepsy. RELN (reelin) [714], HTR2C [379], GABRR1 [715], RIT2 [324], DSCAM (DS cell adhesion molecule) [381], TLR2 [716], PCDH15 [331], TSPAN8 [333], LHX5 [717], MBP (myelin basic protein) [718], CUX2 [719], ACAN (aggrecan) [334], SHH (sonic hedgehog signaling molecule) [720], P2RX7 [721], CACNA2D4 [722], CDH13 [392], CSMD1 [723], NTF3 [724], GRIA4 [725], GRIA2 [725], MMP3 [726], GABRB2 [727], HLA-DRB1 [728], VWF (von Willebrand factor) [353], MDGA1 [355], LDB2 [729], NR2E1 [730], BRCA2 [362], ST8SIA2 [731], KITLG (KIT ligand) [732], GRM3 [733] and COMT (catechol-O-methyltransferase) [412] could act as diagnosis and prognosis biomarkers for bipolar disorder. However, further investigations are needed to explore and confirm the potentially significant GO terms and signaling pathways for HD and to achieve a comprehensive understanding of this process.

Based on the PPI network constructed and modules analysed by the online database IID, we identified hub genes. LRRK2 [115], HOXA1 [734], IL7R [150], EGFR (epidermal growth factor receptor) [198], NEDD4L [187] and COMT (catechol-O-methyltransferase) [207] have been identified in inflammation. LRRK2 [224], IL7R [244], ERBB3 [255] and EGFR (epidermal growth factor receptor) [276] altered expression has been closely associated with diabetes mellitus. LRRK2 [287], IL7R [300], EGFR (epidermal growth factor receptor) [313] and COMT (catechol-O-methyltransferase) [314] have been revealed to be altered expressed in multiple sclerosis. LRRK2 [320], ERBB3 [340], EGFR (epidermal growth factor receptor) [367] and COMT (catechol-O-methyltransferase) [370] have been identified to be involved in the development of schizophrenia. Previous studies have shown that LRRK2 [376], ERBB3 [391], EGFR (epidermal growth factor receptor) [406], NEDD4L [401] and COMT (catechol-O-methyltransferase) [412] might influence the prognosis in depression. LRRK2 [419], EGFR (epidermal growth factor receptor) [431] and COMT (catechol-O-methyltransferase) [437] have been demonstrated to be regulated in neurodegenerative diseases. LRRK2 [450], HOXA1 [735], IL7R [477], EGFR (epidermal growth factor receptor) [276], NEDD4L [496] and COMT (catechol-O-methyltransferase) [512] are associated with cardiovascular diseases development. Altered expression of LRRK2 [522], EGFR (epidermal growth factor receptor) [554] and COMT (catechol-O-methyltransferase) [561] promotes Parkinson’s disease. A previous study reported that the LRRK2 [522], MTUS2 [736], EGFR (epidermal growth factor receptor) [610], NEDD4L [602] and COMT (catechol-O-methyltransferase) [616] genes were associated with Alzheimer’s disease. LRRK2 [620], EGFR (epidermal growth factor receptor) [636] and NEDD4L [632] are critical in amyotrophic lateral sclerosis development. In many clinical studies, LRRK2 [649] and EGFR (epidermal growth factor receptor) [658] have been linked to the start of dementia. LRRK2 [660], EGFR (epidermal growth factor receptor) [701], NEDD4L [695], COMT (catechol-O-methyltransferase) [705] and CDK18 [737] are involved in the regulation of stroke. ERBB3 [82], EGFR (epidermal growth factor receptor) [100], NEDD4L [95] and COMT (catechol-O-methyltransferase) [104] expression levels were significantly altered in hypertension. A previous study found that EGFR (epidermal growth factor receptor) [447] is positively correlated with the severity of autism spectrum disorder, suggesting its potential as a biomarker for autism spectrum disorder. COMT (catechol-O-methyltransferase) [648] is known to be involved in the development of HD. COMT (catechol-O-methyltransferase) [412] protein levels might together be helpful for diagnosing bipolar disorder. Our findings suggest that novel biomarkers include TEX101, WDR76, KRT86, ZNF835, ZNF572 and BEGAIN (brain enriched guanylate kinase associated) might influence the progression of HD. However, the exact pathogenic mechanism remains unknown. This invesstigation might provide reference for research the connection between HD and its associated complication.

MiRNA-hub gene regulatory network analyses and TF-hub gene regulatory network are formed from the interactions of hub genes, miRNAs and TFs, and participate in all steps of the life process, including biosignal transfer, gene expression control, energy and substance metabolism, and cell cycle control. Previous studies have been reported that EPAS1 [167], HOXA1 [734], TGM2 [114], CCND2 [191], MAP1B [204], EGFR (epidermal growth factor receptor) [198], NEDD4L [187], DCLK1 [196], hsa-mir-363-3p [738], SMAD4 [739], ATF3 [740], TBX5 [741] and CREB1 [742] increased the risk of inflammation development. Previous studies have reported that EPAS1 [486], HOXA1 [735], TGM2 [449], CCND2 [500], EGFR (epidermal growth factor receptor) [276], NEDD4L [496], DCLK1 [504], hsa-mir-363-3p [738], hsa-mir-21-5p [743], SMAD4 [744], ATF3 [745], TBX5 [746], SREBF1 [747], SMARCA4 [748] and CREB1 [749] were regulated in cardiovascular diseases. Studies reported that ERBB3 [255], TGM2 [223], CCND2 [274], EGFR (epidermal growth factor receptor) [276], hsa-mir-21-5p [750], ATF3 [751], SREBF1 [752], ARNT (aryl hydrocarbon receptor nuclear translocator) [753] and CREB1 [754] play important roles in diabetes mellitus. ERBB3 [340], TGM2 [319], EGFR (epidermal growth factor receptor) [367], DCLK1 [366], SMAD4 [755], SREBF1 [756] and CREB1 [757] participated in the regulation of schizophrenia. ERBB3 [391], MAP1B [410], EGFR (epidermal growth factor receptor) [406], NEDD4L [401], hsa-mir-21-5p [758] and CREB1 [759] were involved in depression. ERBB3 [82], EGFR (epidermal growth factor receptor) [100], NEDD4L [95] and SREBF1 [760] regulated the pathogenesis of hypertension. HSPA2 [43], MTUS2 [736], CCND2 [605], MAP1B [614], EGFR (epidermal growth factor receptor) [610], NEDD4L [602], hsa-mir-4487 [761], hsa-mir-21-5p [762] and CREB1 [763] were involved in regulating the occurrence of Alzheimer’s disease. TGM2 [286], EGFR (epidermal growth factor receptor) [313] and DCLK1 [312] have been shown to be associated with multiple sclerosis. Altered levels of TGM2 [418], MAP1B [434], EGFR (epidermal growth factor receptor) [431], hsa-mir-21-5p [764] and CREB1 [765] have been observed in neurodegenerative diseases patients. Some studies have shown that TGM2 [640], hsa-mir-10a [766], ATF3 [767] and CREB1 [768] plays a certain role in HD. Altered expression of MAP1B [557], EGFR (epidermal growth factor receptor) [554], SREBF1 [769] and CREB1 [770] promotes Parkinson’s disease. MAP1B [637], EGFR (epidermal growth factor receptor) [636], NEDD4L [632], hsa-mir-21-5p [771], ATF3 [772] and SREBF1 [773] were significantly altered in patients with amyotrophic lateral sclerosis. A previous study reported that the EGFR (epidermal growth factor receptor) [658], hsa-mir-21-5p [762] and SREBF1 [774] biomarkers were associated with the dementia. Previous studies have shown that EGFR (epidermal growth factor receptor) [701], NEDD4L [695], hsa-mir-21-5p [775] and ATF3 [776] are closely associated with stroke. EGFR (epidermal growth factor receptor) [447] have recently been found in autism spectrum disorder. Altered expression of hsa-mir-4521 [777] was found to promote the epilepsy. Hence, the results of miRNA-hub gene regulatory network analyses and TF-hub gene regulatory network were in accordance with experimental results on HD. The roles of novel biomarkers in the network were determined, such as KRT8, TOP2A, hsa-mir-1292-5p, hsa-mir-432-3p, hsa-mir-6801-3p, hsa-mir-7977, hsa-mir-6807-5p, ESRRB (estrogen related receptor beta), EED (embryonic ectoderm development) and EP300, which expanded our knowledge on development of HD. Hence, our findings might be a novel clue for the diagnosis and treatment of HD.

## Conclusion

Based on the current investigation, it is feasible to use bioinformatics methods to screen and identify essential molecular markers and signaling pathways related to HD. Bioinformatics methods can be used to analyze NGS data to discover disease-related molecular features such as gene expression, protein structure, and metabolite levels. In this study, the hub genes, miRNA and TFs related to HD were identified. Our findings will help to encourage further studies on the biological functions of hub genes, miRNA and TFs.

## Acknowledgement

I thanks very much to Steven Goldman, University of Rochester Medical Center, Center for Translational Neuromedicine, Goldman Lab, Rochester, NY, USA, the author who deposited their NGS dataset GSE105041, into the public GEO database.

## Ethical approval

This article does not contain any studies with human participants or animals performed by any of the authors.

## Informed consent

No informed consent because this study does not contain human or animals participants.

## Availability of data and materials

The datasets supporting the conclusions of this article are available in the GEO (Gene Expression Omnibus) (https://www.ncbi.nlm.nih.gov/geo/) repository. [(GSE105041) https://www.ncbi.nlm.nih.gov/geo/query/acc.cgi?acc=GSE105041]

## Consent for publication

Not applicable.

## Author Contributions

1. B. V. - Writing original draft, and review and editing

2. C. V. - Software and investigation

